# Hemoglobin as a peroxidase and drug target for oxidative stress-related diseases

**DOI:** 10.1101/2024.05.21.594979

**Authors:** Woojin Won, Elijah Hwejin Lee, Lizaveta Gotina, Heejung Chun, Jae-Hun Lee, Uiyeol Park, Daeun Kim, Tai Young Kim, Jiwon Choi, Yoowon Kim, Sun Jun Park, Mridula Bhalla, Jiwoon Lim, Jong-Hyun Park, Soo-Jin Oh, Hoon Ryu, Ae Nim Pae, Ki Duk Park, C. Justin Lee

## Abstract

Hemoglobin (Hb) is well-known for transporting oxygen in red blood cells within blood vessels^1^. Although Hb is also present in the brain^2^, its role remains poorly understood. Here, we show that Hb, found in astrocytes of neurodegenerative animal models and patients, displays significant antioxidant effects through its H_2_O_2_-decomposing peroxidase activity, and a small molecule enhancer boosts this activity, reducing aberrant H_2_O_2_ and mitigating H_2_O_2_-induced neurodegeneration. To counteract the harmful effects of aberrant H_2_O_2_-production in Alzheimer’s disease (AD), we developed KDS12025, a blood-brain barrier (BBB)-permeable small molecule that effectively enhances the peroxidase activity of Hb by a hundredfold, especially at a low level of Hb. KDS12025 and its analogs achieve this enhancement through its electron-donating amine group. KDS12025 reduces H_2_O_2_ levels in astrocytes, exhibits neuroprotective effects, and reverses memory impairment in AD models. Gene-silencing of Hbβ abrogates KDS12025’s impact in both culture and animal models of AD. Moreover, KDS12025 prevented the death of dopaminergic neurons in a Parkinson’s disease (PD) model without altering the oxygen-transporting function of Hb. KDS12025 extended survival and improved motor function even in the severe amyotrophic lateral sclerosis (ALS) mouse model. Our findings propose Hb as a new therapeutic target for neurodegenerative diseases, with KDS12025 emerging as a first-in-class drug candidate that enhances Hb’s peroxidase activity to reduce H_2_O_2_. Boosting Hb’s peroxidase activity with KDS12025 mitigates oxidative stress and alleviates neurodegeneration in AD, PD, and ALS with broad applicability for numerous oxidative-stress-driven diseases.

## Introduction

Hemoglobin (Hb) is a well-known heme-containing protein recognized for transporting oxygen and carbon dioxide in the bloodstream^1^. Intriguingly, Hb has also been observed in the brain, but its function remains enigmatic^2,3^. Brain Hb degradation can exacerbate neurodegenerative diseases by releasing iron and heme, leading to inflammation and the generation of reactive oxygen species (ROS)^1,3^. For example, overexpression of Hb in dopaminergic neurons or higher brain Hb levels in the elderly increases the risk of neurodegenerative diseases^4,5^. Paradoxically, in cultured astrocytes, Hb administration is protective, potentially suppressing amyloid β-mediated inflammatory activation and oxidative stress^6,7^. However, the cellular and molecular mechanism underlying the antioxidant effect of Hb is entirely unknown.

ROS are known to cause a myriad of diseases, including neurodegenerative disorders like AD^8,9^, PD^10^, ALS^11^, and various peripheral inflammatory conditions such as atherosclerosis^12^, acute pancreatitis^12^, and rheumatoid arthritis (RA)^13^. Our recent research highlights excessive H_2_O_2_ production by reactive astrocytes as a key factor in neurodegeneration in AD^8^ and PD^10^ and inflammation in RA^13^. However, developing effective treatments has been almost impossible due to the challenge of regulating H_2_O_2_ and other ROS without disrupting cellular redox balance^12,14^. Conventional antioxidants like EGCG and vitamin C suffer from poor cellular uptake and low bioavailability, making effective radical scavenging extremely difficult *in vivo* due to their high reactivity of radicals, thus severely limiting their therapeutic success^12,15^. To make matters worse, the unclear mode of action of dietary antioxidants like curcumin and resveratrol complicates their development into effective drugs^15^. Therefore, there is a desperate need to develop novel antioxidant strategies that indirectly reduce H_2_O_2_ levels without disturbing brain function or disrupting redox balance.

Based on our serendipitous finding using an HRP-based H_2_O_2_ assay, we elucidate an unprecedented antioxidative mechanism of Hb functioning as an H_2_O_2_-decomposing peroxidase and a BBB-permeable small molecule, KDS12025, acting as a hydrogen bond donor that enhances Hb’s peroxidase activity. In the brain, H_2_O_2_-induced oxidative stress reduces Hb expression, particularly in astrocytes in the hippocampus and substantia nigra pars compacta (SNpc), which is fully reversed by KDS12025 treatment. Even at very low doses, KDS12025 shows high efficacy in animal models of AD, PD, ALS, and RA, highlighting its potential as a broad-spectrum therapeutic for oxidative stress-related diseases.

### Hemoglobin shows peroxidase-like activity

Previously, we utilized 2-hydroxy-5-[2-(4-trifluoromethyl-phenyl)-ethylamino]-benzoic acid (HTPEB)^8,16^ as a putative H_2_O_2_ scavenger in animal models of AD, but its cellular and molecular mechanism was unclear. Our previous results from the horseradish peroxidase (HRP)-based H_2_O_2_ detection assay hinted that HTPEB enhances peroxidase activity^8^, paving the way for developing a new peroxidase enhancer in the field of neurodegenerative diseases. To determine the detailed molecular mechanism of HTPEB’s enhancement and to test if Hb exhibits H_2_O_2_-decomposing activity, we used H_2_O_2_ assays with Amplex Red or ROS-Glo (Fig. 1a,b). HTPEB showed H_2_O_2_ scavenging activity in the Amplex Red assay (Fig. 1a,c), which includes HRP. The proximal (His170) and distal histidine (His42) residues interact with the iron in the heme group and H_2_O_2_ respecively^17,18^. However, HTPEB did not show activity in the ROS-Glo assay (Fig. 1b,d), indicating HTPEB’s dependence on HRP. Replenishing HRP restored HTPEB’s H_2_O_2_-decomposing activity in ROS-Glo (Fig. 1e), lowering EC_50_, doubling the activity rate, and reducing H_2_O_2_ reduction time (Extended Data Fig. 1b,c). These results indicate that HTPEB’s enhancement of peroxidase activity requires HRP.

**Fig. 1.**
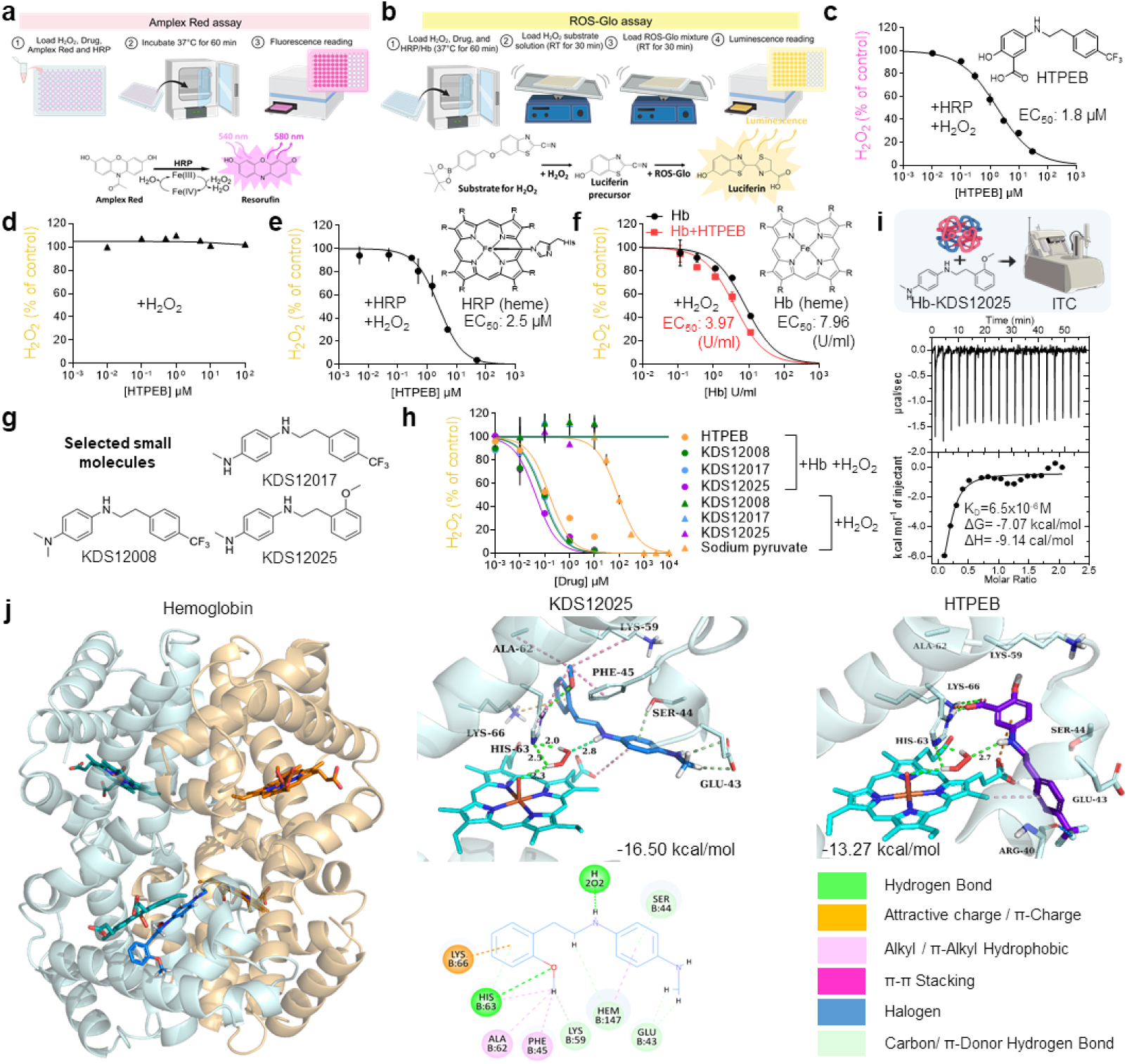
Hemoglobin’s peroxidase-like activity: developing an enhancer for decomposition of H_2_O_2_. **a,b,** Schematic diagram of HRP-dependent (**a;** Amplex Red) and HRP-independent (**b;** ROS-Glo) H_2_O_2_ assay. **c,** Dose-response curve for HTPEB of Amplex Red assay. **d,e,** Dose-response curve for HTPEB of ROS-Glo assay without (**d**) or with (**e**) replenishing HRP. **f,** Dose-response curve of Hb in ROS-Glo assay with (red square shapes) or without (black circle shapes) an HTPEB. **g,** Chemical structure of KDS derivatives small molecules (KDS12008, 17, and 25) which retains the essential *N*-phenethylaniline core. **h,** Dose-response curve for KDS derivatives with (circle) or without (triangle) Hb of ROS-Glo assay (EC_50_ with Hb in μM: HTPEB, 0.15; KDS12008, 0.08; KDS12017, 0.09; KDS12025, 0.04). **i,** ITC analysis depicting the binding interaction between Hb and KDS12025. **j,** Binding mode and calculated binding energy (ΔG_bind_) of KDS12025 (middle) and HTPEB (right) with Hb proposed by docking simulations; schematic of interactions between heme and key residues (bottom). Dose-response curve and EC50 were calculated and determined by fitting data with GraphPad Prism software. Data are presented as the mean ± s.e.m.

While HRP is a plant-derived heme-containing peroxidase and absent in animals^17^, we substituted it with animal-derived proteins similar in structure or function to HRP, like catalase (CAT), glutathione peroxidase (GPx), and Hb^19,20^ to mimic HTPEB’s enhancement of HRP activity. Like HRP, Hb, a tetrameric heme-containing protein, significantly facilitated H_2_O_2_-decomposition by 2-fold with HTPEB (Fig. 1f). Contrarily, CAT^20^ did not show enhancement with HTPEB (Extended Data Fig. 1d), possibly due to its inherent maximal catalytic activity linked to a tyrosine residue^21^. GPx, a heme-lacking and selenium-containing peroxidase^20^, showed decreased activity with HTPEB (Extended Data Fig. 1e), possibly due to competition with GPx’s substrate glutathione. These results indicate a peroxidase-like activity of Hb with enhancement by HTPEB.

### Developing potent BBB-permeable H_2_O_2_-decomposing chemical enhancers for Hb

Although HTPEB enhances Hb’s peroxidase-like activity, it suffers from poor BBB permeability and low solubility (Supplementary Table 3). To find a better enhancer, we performed structure-function analysis on different functional groups within HTPEB. We found only mesalazine enhanced Hb’s peroxidase-like activity (Supplementary Table 1), suggesting the electron-donating amine group directly bound to an aromatic ring may be critical for HTPEB’s potency. To improve the pharmacological properties of Hb-enhancers, we synthesized strategically designed HTEPB derivatives, retaining *N-*phenethylaniline core, by modifying the carboxylic acid and hydroxyl groups with an electron-donating methylamine group (Extended Data Fig. 2). This led to KDS12017 (EC_50_ = 0.09 µM) and KDS12025 (EC_50_ = 0.04 µM) with higher potency, the latter being about 4-fold more potent than HTPEB (EC_50_ = 0.15 µM) (Fig. 1g,h and Supplementary Table 2). Unlike direct H_2_O_2_ scavengers such as sodium pyruvate^22^, these compounds did not degrade H_2_O_2_ without Hb (Fig. 1h). Furthermore, KDS12025 showed high bioavailability (87.43%) and significant BBB-permeability (67.43 × 10^−6^ cm/sec) (Supplementary Table 3). KINOMEscan and off-target assays conferred its selectivity and fewer side effects than other reactive ROS scavengers (Extended Data Fig.3 and Supplementary Tables 4,5).

Next, the isothermal calorimetric assay showed significant binding between KDS12025 and Hb (ΔG_bind_ = −7.07 kcal/mol) (Fig. 1i). Molecular docking suggested a strong interaction (ΔG_bind_ = −16.5 kcal/mol) between KDS12025 and the Hbβ subunit within a wide cleft near the heme molecule, formed by residues 42-50 (Fig. 1j). Hydrophobic interactions of the methoxy group and a hydrogen bonding network between the *N*1 amine group, histidine H63, are crucial for the molecule’s activity (Fig. 1j). HTPEB and KDS12017 were also bound to this site (ΔG_bind_ of −13.27 and −12.99 kcal/mol, respectively), whereas less potent derivatives had lower ΔG_bind_ (Fig. 1j and Extended Data Fig. 4). KDS12025 and HTPEB also fit in the narrow catalytic pocket of HRP (Extended Data Fig. 5). Collectively, these findings demonstrate that *N*-phenethylaniline molecules retain the crucial amine group function and potently bind to Hbβ, with KDS12025 being the most potent enhancer of Hb’s peroxidase-like activity.

### KDS12025 enhances Hb’s peroxidase activity by 100-fold

We quantified and compared the enhancement of peroxidase activity in HRP and Hb by KDS12025, calculating the fold increase over 30 minutes between 0.5 h and 1 h reaction time (Fig. 2a). KDS12025 significantly exerted a 5-fold increase in peroxidase activity, outperforming HTPEB’s 3-fold enhancement (Fig. 2a,b). Then, we assessed Hb’s intrinsic peroxidase activity by obtaining concentration-dependence relationships (Fig. 2c). At low concentrations, Hb showed no significant peroxidase activity, whereas preincubation with KDS12025 and HTPEB significantly enhanced it (Fig. 2d). Higher Hb concentration further improved activity, with KDS12025 showing greater potency than HTPEB (Fig. 2d). When comparing Hb’s H_2_O_2_ decomposition dose-dependently, KDS12025 lowered Hb’s EC_50_ for H_2_O_2_ decomposition by 4.7-fold and its EC_20_ by 98-fold (Fig. 2e). Overall, KDS12025 remarkably enhances the H_2_O_2_-decomposing activity of Hb by about 100-fold, even at extremely low concentrations (0.04 U/ml, 0.03 μg/ml).

**Fig. 2.**
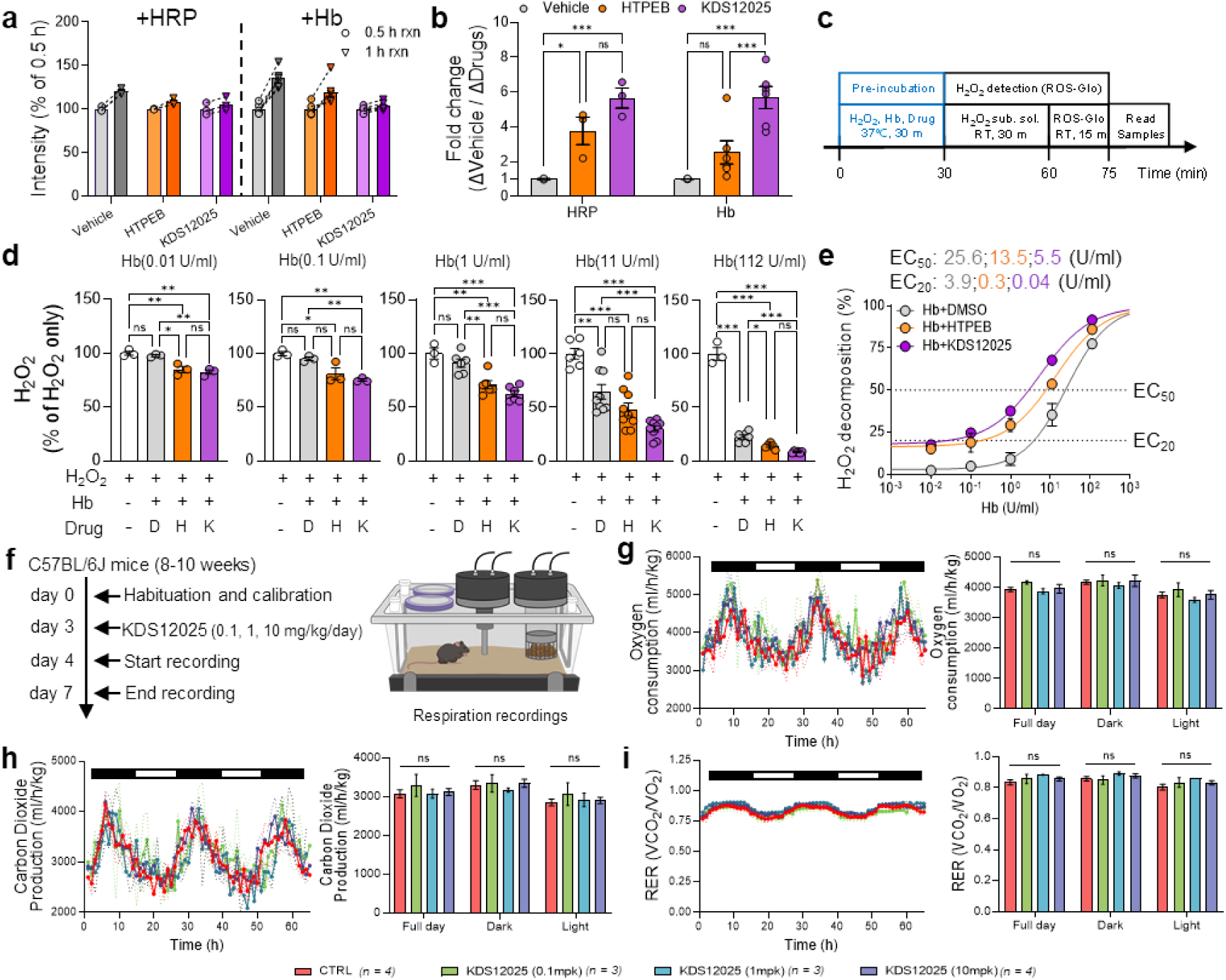
KDS12025 enhances Hb peroxidase activity. **a,** Paired comparison of H_2_O_2_ decomposition by HRP and Hb with drug treatment over 30 minutes using ROS-Glo assay. **b,** Fold change in H_2_O_2_-decomposing facilitation by HRP and Hb with drug treatment. **c,** Timeline of experiments investigating whether Hb has intrinsic peroxidase activity and the extent of peroxidase enhancement by the drug with pre-incubation. **d,** Results of investigating Hb’s peroxidase function and the drug’s (D, DMSO; H, HTPEB; K, KDS12025) effect at various Hb concentrations. The drugs were prepared as 200x stock solutions (final concentration of 10 μM). **e,** Dose-response curve of Hb’s peroxidase activity indicating EC_20_ and EC_50_ values of H_2_O_2_-decomposition at 0.01 U/ml of Hb **f,** Timeline of PhenoMaster experiments with the administration of KDS12025 at different concentrations (0.1, 1, 10 mg/kg/day). **g-i,** Measurement of oxygen consumption (**g**), carbon dioxide production (**h**), and respiratory exchange ratio (RER; **i**) during the night (dark) and day (white) cycles, with a summary graph provided. EC_20_ and EC_50_ were calculated and determined by fitting data with GraphPad Prism software. Data are presented as the mean ± s.e.m. *P < 0.05, **P < 0.01, ***P < 0.001; ns, not significant. Additional statistics are provided in Supplementary Table 7.

Subsequently, we tested whether Hb functions as catalase, an H_2_O_2_-decomposing enzyme producing oxygen as a byproduct and found that Hb is not a catalase and that KDS12025 does not facilitate CAT (Extended Data Fig. 6). To assess if KDS12025 interferes with Hb’s oxygen-carrying capability in the blood, we examined respiratory functions of animals using an automated phenotyping system (Fig. 2f). This analysis showed that KDS12025 at 0.1, 1, and 10 mg/kg/day did not affect oxygen consumption, carbon dioxide production, and the respiratory exchange ratio during dark and light cycles (Fig. 2g-i), as well as energy expenditure, food, or drink consumption (Extended Data Fig. 7). Taken together, these results imply that KDS12025 potently enhances Hb’s peroxidase activity without altering its respiratory function or the metabolism in mice.

### KDS12025 effectively reduces AD-like pathology *in vitro* and *in vivo*

Since primary cultured astrocytes have been shown to express Hb^2^, we tested whether KDS12008, KDS12017, and KDS12025 enhance the H_2_O_2_-decomposing peroxidase activity in astrocytes. Using H_2_O_2_-sensors, DCFDA and oROS-G^23^, we found that oligomerized Aβ induced H_2_O_2_ levels in a concentration-dependent manner in astrocytes, and KDS12025 effectively reduced Aβ-induced H_2_O_2_ with an EC_50_ of 0.4 μM, while 10 mM sodium pyruvate did not (Extended Data Fig. 8a-f). Moreover, long-term live-cell confocal imaging showed that Aβ-or putrescine(a precursor of MAO-B-dependent H_2_O_2_-production^13,24^)-induced aberrant H_2_O_2_ was significantly reduced by KDS12025 treatment within 24 hours (Extended Data Fig. 8g-j). Similarly, we tested whether KDS12008, KDS12017, and KDS12025 alleviate AD-like symptoms in APP/PS1 mice and found that KDS12025 (at 3 and 10 mg/kg/day, 7 times, intraperitoneal injection) most effectively improved memory impairment and normalized reactive astrocytes and microglia (Extended Data Fig. 9). These results indicate that KDS12025 is the most effective enhancer *in vitro* and *in vivo*.

### The broad therapeutic effect of KDS12025 in AD, PD, ALS, and RA mouse models

To explore the broader therapeutic potential of KDS12025 in treating AD, PD, and ALS, as well as peripheral inflammatory diseases like RA, we conducted comprehensive studies in animal models of each disease. To investigate if KDS12025 alleviates neurodegeneration, we used the previously described animal model of focal DTR-expressing astrocytes in APP/PS1 mice (fGiD)^8^. Passive avoidance test (PAT) and novel place recognition (NPR)^25^ test showed memory and cognitive impairments in fGiD mice, which were significantly recovered by KDS12025 (3 mg/kg/day, 16 times, intraperitoneal injection) without altering locomotion (Fig. 3a-c and Extended Data Fig. 10a). Severe reactive astrocytes and a decreased number of NeuN (signs of H_2_O_2_-dependent neurodegeneration) in fGiD mice were reverted to control levels by KDS12025 (Fig. 3d-f). Furthermore, KDS12025 effectively restored astrogliosis, aberrant astrocytic GABA, 8-OHdG (oxidative stress marker), and tonic GABA inhibition current^26^ in APP/PS1 to wild-type levels (Extended Data Fig. 11b-i; Extended Data Fig. 9k-n). Even KDS12025 at 0.1 mg/kg/day also reversed memory impairment in the GiD mouse model of AD (Extended Data Fig. 10c). These results indicate that KDS12025 effectively and potently reverses key AD pathological features, including astrogliosis, oxidative stress, neurodegeneration, and memory impairment.

**Fig. 3.**
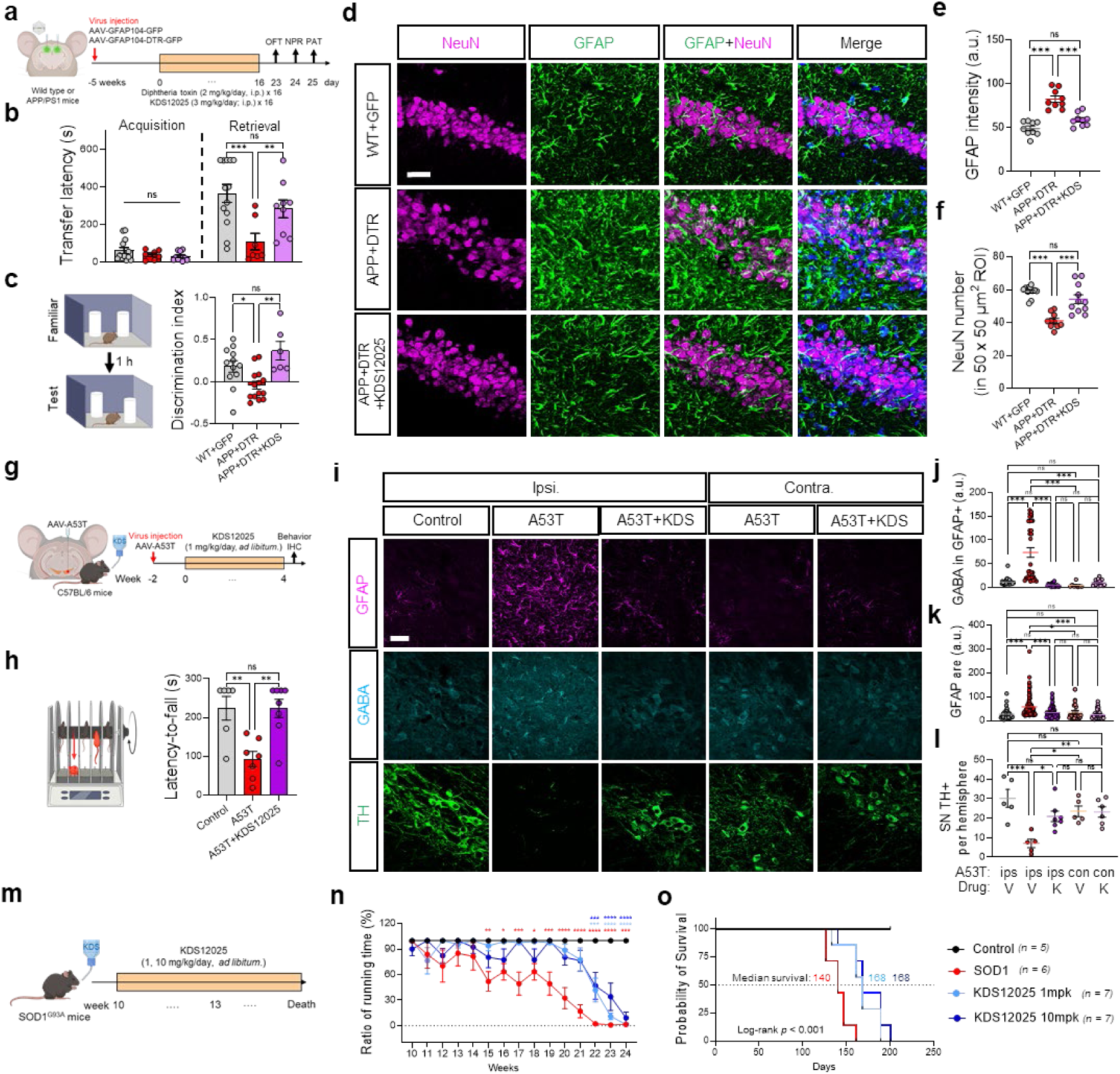
The broad therapeutic effect of KDS12025 in AD, PD, and ALS models. **a,** Schematic timeline of reactive astrocytes in APP/PS1 mice using the focal expression of DTR (fGiD) in astrocytes and intraperitoneal (i.p.) injection of KDS12025. **b,** Transfer latency to enter the dark chamber of control, fGiD, and fGiD+KDS mice. **c,** Discrimination index in NPR test. **d,** Representative images of the hippocampus CA1 for NeuN and GFAP in WT, fGiD, and fGiD+KDS mice. Scale bar, 10 μm. **e,** Mean intensity of GFAP. **f,** Quantification of neurodegeneration with NeuN number per 50 x 50 μm^2^. **g,** Schematic timeline of A53T virus-induced PD model and KDS12025 treatment (drinking *ad libitum*). **h** Schematic diagram of rotarod test (left) and latency-to-fall of control, A53T, and A53T+KDS mice. **i,** Representative images of the SN region with ipsilateral and contralateral side for GFAP, GABA, and TH of control, A53T, A53T+KDS mice. Scale bar, 10 µm. **j,** Mean intensity of GABA in GFAP-positive area. **k,** Measurement of GFAP-positive area. **l,** Quantification of TH-positive neurons per hemisphere in SN. **m,** Schematic timeline of SOD1^G93A^ mice with KDS12025 treatment (drinking *ad libitum*). **n,** Ratio of running time of control, SOD1, and SOD1+KDS (1, 10 mg/kg/day) mice in the rotarod test. **o,** Probability of survival of control, SOD1^G93A^, and SOD1^G93A^+KDS (1, 10 mg/kg/day) mice. Data are presented as the mean ± s.e.m. *P < 0.05, **P < 0.01, ***P < 0.001; ns, not significant. Additional statistics are provided in Supplementary Table 7.

Next, to test KDS12025’s effectiveness in other neurodegenerative diseases such as PD, we used a previously described A53T overexpression model of PD^27^ and treated with KDS12025 (1 mg/kg/day, 4 weeks, drinking *ad libitum*) (Fig. 3g). We observed a significant motor impairment of A53T mice, which was fully rescued by KDS12025 (Fig. 3h). Tyrosine hydroxylase (TH)-positive dopaminergic neuronal death, reactive astrocytes, and abnormally elevated astrocytic GABA levels in the ipsilateral SNpc of the A53T model compared to the contralateral side were effectively reversed by KDS12025 treatment (Fig. 3i-l and Extended Data Fig. 12).

Given the lack of small molecule drugs against ALS that extend survival, we next examined the severe animal model of ALS, SOD1^G93A^ mice to evaluate the effect by KDS12025 (1 and 10 mg/kg/day, drinking *ad libitum*) (Fig. 3m). KDS12025 significantly delayed the motor impairment by more than 7 weeks and extended the median survival from 140 to 168 days (Fig. 3n,o). This is the first-in-class small molecule that shows a significant extension of survival in this severe animal model of ALS. Additionally, we tested the effect of KDS12025 on an animal model of inflammation-associated RA^13^ and found that KDS12025 significantly mitigated the RA pathology (Extended Data Fig. 13a-m). Altogether, these results highlight KDS12025 as a potential broad-spectrum therapeutic for various neurodegenerative and H_2_O_2_-associated inflammatory diseases.

### Reduced astrocytic Hbβ in neurodegenerative diseases and its reversal by KDS12025

Although KDS12025 has shown effectiveness, its connection to brain Hb remains elusive. To assess the significance of the brain Hbβ and its clinical relevance in AD, we performed immunohistochemistry on postmortem hippocampal tissues from normal subjects and AD patients, focusing on Hbβ due to computational modeling results and previous evidence of its significant impact on cognitive impairment^28^. We observed the prevalent expression of Hbβ in astrocytes, which was significantly reduced in AD patients compared to normal subjects (Fig. 4a,b). These findings parallel the reduced expression pattern of Hbβ in the AD mouse model and human brain tissue (Extended Data Fig. 14). Furthermore, a similar reduction in Hbβ expression was found in the A53T mouse model of PD (Extended Data Fig. 14), providing its potential as a molecular target for KDS12025.

**Fig. 4.**
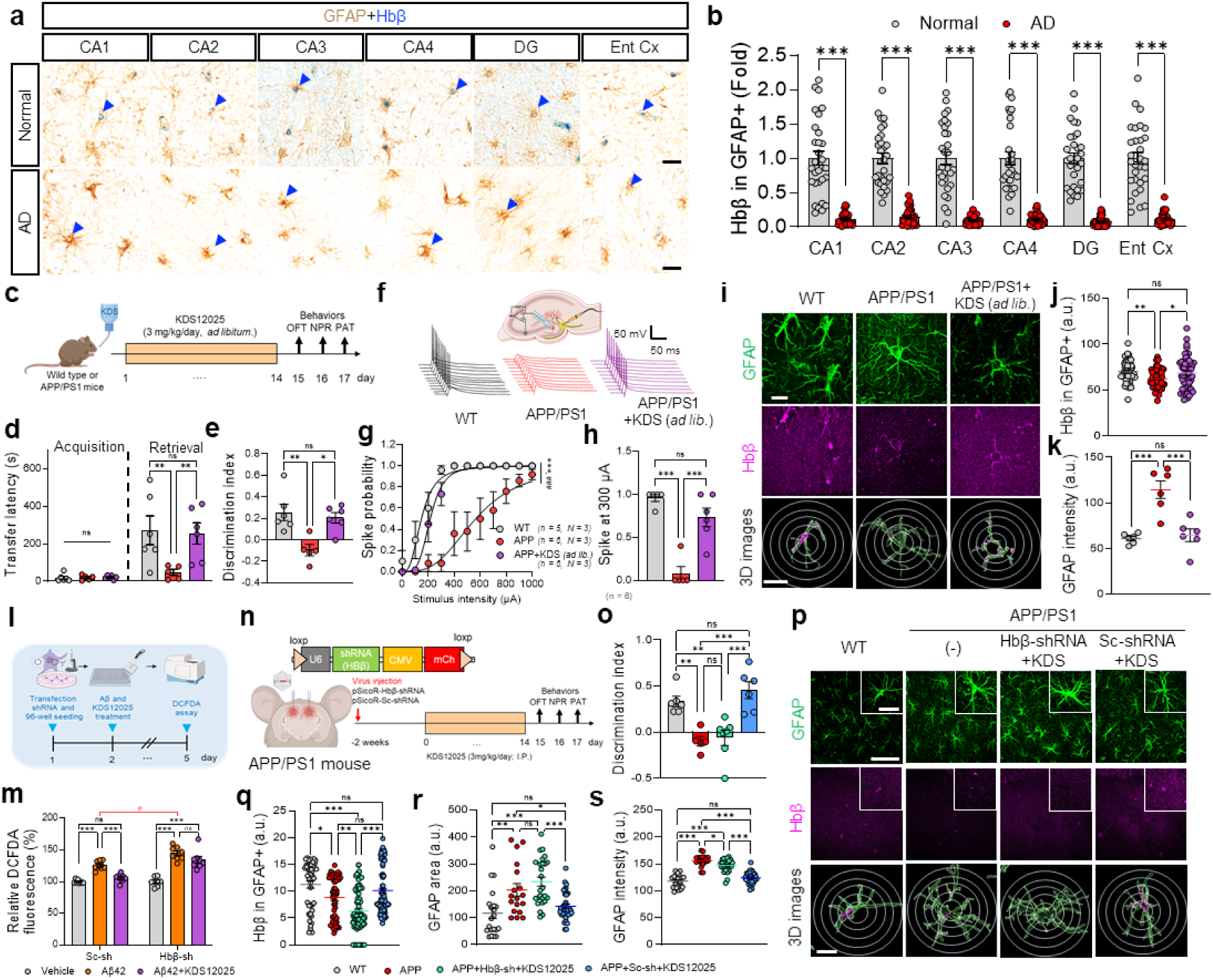
Hbβ is decreased in the hippocampus of AD patients and necessary for KDS12025 action. **a,** Representative images of postmortem hippocampal tissues for GFAP and Hbβ in normal subjects *(n = 3)* and AD patients *(n = 3)*. Scale bar, 20 μm. **b,** Quantification of Hbβ intensity in GFAP of each hippocampal sub-region **c,** Schematic timeline of APP/PS1 mice and KDS12025 treatment (3 mg/kg/day; drinking *ad libitum*), followed by performing behaviors. **d,** Transfer latency to enter the dark chamber of WT, APP, and APP+KDS mice. **e,** Discrimination index in NPR test. **f,** Schematic of spike probability with electrical stimulation of perforant pathway stimulation in the hippocampus and a representative trace of evoked EPSPs in WT, APP, and APP+KDS mice. **g,** Spike probability within stimulus intensity of 100-1000 μA. **h,** Comparison of spike probability at 300 μA. **i,** Representative Lattice-SIM images of the hippocampus for GFAP and Hbβ of WT, APP, and APP+KDS mice. Representative 3D images from Imaris software (green, GFAP; magenta, Hbβ) and Sholl analysis (circles). Scale bars, 20 μm (main); 10 μm (Imaris). **j,** Mean intensity of Hbβ in GFAP. **k,** Mean intensity of GFAP. **l,** Schematic timeline of transfection Hbβ-shRNA with Aβ (5 μM) and KDS12025 (10 μM). **m,** Relative H_2_O_2_ levels in Sc- or Hbβ-shRNA transfected with the vehicle, Aβ, and Aβ+KDS. The ‘^#^’ symbol (red) compares Aβ-induced H_2_O_2_ between Sc- and Hbβ-shRNA. **n,** Schematic timeline of injecting an AAV carrying pSicoR-Hbβ (or Sc)-shRNA-mCherry into the *stratum radiatum* of the hippocampus in APP/PS1 mouse. **o,** Discrimination index from the NPR test. **p,** Representative images of the hippocampus for GFAP and Hbβ in WT, APP, APP+Sc-shRNA+KDS, and APP+shRNA+KDS. Representative 3D images from Imaris software. Scale bars, 50 μm (main); 10 μm (inset); 5 μm (Imaris). **q,** Mean intensity of Hbβ in GFAP. **r,** GFAP-positive area. **s,** Mean intensity of GFAP. Data are presented as the mean ± s.e.m. *,^#^P < 0.05, **P < 0.01, ***P < 0.001; ns, not significant. Additional statistics are provided in Supplementary Table 7.

We then investigated KDS12025’s effect on astrocytic Hbβ in the APP/PS1 mice. As the intraperitoneal injection (Fig. 3a and Extended Data Fig. 9), oral administration of KDS12025 via drinking *ad libitum* (3 mg/kg/day for 14 days) effectively reversed memory impairment in APP/PS1 mice (Fig. 4c-e). Consistently, KDS12025 treatment significantly restored the spike probability of synaptically induced action potential firing in dentate gyrus granule cells^29^ of the APP/PS1 mice to the wild-type level (Fig. 4f-h). Immunohistochemistry showed a significant decrease in astrocytic Hbβ in the APP/PS1 mice (Fig. 4i,j), with a minimal basal expression in pyramidal neurons (Extended Data Fig. 11a). Remarkably, KDS12025 reverted astrocytic Hbβ level to that of wild-type (Fig. 4i,j), suggesting that H_2_O_2_ negatively regulates astrocytic Hbβ. Consistently, astrocytic Hbβ, rather than Hbβ in TH-positive neurons, showed a similar pattern in the animal model of PD (Extended Data Fig. 15). These results suggest that KDS12025’s effect of alleviating AD- and PD-like symptoms is associated with astrocytic Hbβ.

### Astrocytic Hbβ is necessary for the action of KDS12025 in the hippocampus

To determine astrocytic Hbβ as the KDS12025’s molecular target, we developed and used short hairpin RNA (shRNA) to gene-silence Hbβ in primary cultured astrocytes (Fig. 4l and Extended Data Fig. 16a,b). KDS12025 significantly reduced Aβ-induced H_2_O_2_ levels in control scrambled shRNA (Sc-shRNA) astrocytes but not in Hbβ-silenced astrocytes, indicating that Hbβ is necessary for KDS12025’s action (Fig. 4m). Interestingly, gene-silencing of Hbβ exacerbated Aβ-induced H_2_O_2_ compared to Sc-shRNA (Fig. 4m), indicating an increased vulnerability to oxidative stress in the absence of Hbβ.

Next, we injected AAV-pSicoR-Hbβ-shRNA-mCherry virus into the hippocampus of APP/PS1 mice and observed a significantly reduced astrocytic Hbβ (Fig. 4n,p,q), indicating a successful gene-silencing. While KDS12025 treatment significantly reverted astrocytic hypertrophy and improved memory in the Sc-shRNA, it failed to revert in Hbβ-shRNA mice (Fig. 4o-s and Extended Data Fig. 16c-g). Taken together, these results indicate that hippocampal astrocytic Hbβ is necessary for the action of systemically administered KDS12025 in alleviating astrogliosis and hippocampus-dependent memory impairment in the animal model of AD.

## Discussion

In this study, we demonstrate that Hb exhibits peroxidase activity and highlight the significant decline of astrocytic Hbβ levels in the mouse and human brains of neurodegenerative diseases. We have synthesized KDS12025, a small molecule with BBB-permeability and nanomolar potency, which enhances Hb’s peroxidase activity by 100-fold, effectively decomposing aberrant H_2_O_2_ even at extremely low levels of Hb. Without affecting Hb’s primary respiratory function, KDS12025 restores astrogliosis, memory, motor function, oxidative stress, and astrocytic Hbβ levels in AD and PD mouse models. Gene-silencing of astrocytic Hbβ recapitulates the necessity for KDS12025’s H_2_O_2_-decomposing effect. KDS12025 also shows its efficacy in ALS and RA, suggesting a broader antioxidant role of Hb in various inflammatory diseases. Therefore, KDS12025 offers a novel approach for effectively targeting oxidative stress in diseases like AD, PD, ALS, and RA (Extended Data Fig. 17).

Our study provides compelling lines of evidence that Hb exhibits antioxidative function through H_2_O_2_-decomposing peroxidase activity, not catalase-like activity, aligning with previous plasma study^30^. Previous quantum mechanics studies suggest that Arg38 and distal His42 are crucial for proton transfer between the H_2_O_2_ and water in the first step of the HRP cycle^18,31^(Extended Data Fig. 18a,b). In contrast, Hbβ lacks H-bond donating residues near distal His63^18^, which may explain Hb’s lower peroxidase activity than HRP. It is possible that KDS12025 enhances Hb’s peroxidase activity by 100-fold, with its aniline group acting as a missing polar residue that assists in coordinating H_2_O_2_ and H_2_O to the iron atom (Extended Data Fig. 18c). This enhancement may improve Hb’s thermodynamic kinetic ability to reduce H_2_O_2_. Further research is needed to test this exciting possibility of how KDS12025 interacts with Hb, H_2_O_2_, and H_2_O (Extended Data Fig. 18d).

H_2_O_2_ significantly contributes to neurodegenerative diseases^8,12,14^, exacerbating protein aggregation like Aβ (Extended Data Fig. 17) and impairing Aβ clearance^32^. Traditional small molecule antioxidants have failed to alleviate oxidative stress *in vivo* due to poor brain compatibility, high-dosage requirements, and high reactivity, limiting clinical use^12,14^. Hence, KDS12025 would be the first-in-class drug candidate that enhances Hb’s H_2_O_2_-decomposing peroxidase activity *in vivo* without directly scavenging H_2_O_2_, offering a new therapeutic approach. It boosts Hb’s capacity to decompose H_2_O_2_ up to 20%, whereas Hb shows virtually no peroxidase activity without KDS12025. This unique feature circumvents the limitations of traditional H_2_O_2_-scavengers like sodium pyruvate (800 mg/kg)^33^ and *N*-acetylcysteine (200 mg/kg)^34^. Indeed, we observed a potent effect in alleviating AD symptoms with an extremely low dose of 0.1 mg/kg/day, even by drinking (Extended Data Fig. 10c). Thus, KDS12025 promises an effective treatment option for neurodegenerative diseases by enhancing Hb’s antioxidative properties at low doses with minimal side effects.

Our findings also propose that oxidative stress influences astrocytic Hbβ in the hippocampus and SNpc, leading to decreased levels in AD and PD, similar to reduced brain Hbβ mRNA and protein levels in AD and PD patients^28,35,36^. Paradoxically, while Hbβ increases in the cerebrospinal fluid (CSF) of mild cognitive impairment patients^37^, acute H_2_O_2_ exposure in astrocytes increases Hbβ mRNA and opens its promoter regions (Extended Data Fig. 16h-j), suggesting initial antioxidant defense. However, Hbβ eventually seems to leak into the CSF in neurodegenerative states, depleting Hbβ in the brain^37^. Based on those observations, we hypothesize that excessive H_2_O_2_, if unresolved in time, leads to irreversible severe astrogliosis and chronic oxidative stress that further depletes astrocytic Hbβ, perpetuating a vicious cycle to accelerate AD progression. In other words, once the reactive astrocytes become severely reactive and engage in the vicious cycle, the condition may become irreversible, where targeting and removing Aβ plaques might be ineffective, particularly in late-stage AD. These new concepts can provide plausible explanations for the recent setbacks of the immunotherapy-based drug candidates that attempt to remove Aβ plaques. In the PD model, as previously known, Hbβ is also observed in TH-positive neurons^2^. Although we did not find any changes in its levels, we observed significant changes in astrocytic Hbβ, suggesting that astrocytic Hbβ plays a more critical role in neuronal vulnerability to oxidative stress and neurodegeneration in our experiments. In contrast, KDS12025 can liberate the hippocampal and SNpc astrocytes from the vicious cycle to enter into a virtuous cycle of recovered astrocytic Hbβ levels and decompose H_2_O_2,_ which further reduces oxidative stress and accelerates recovery from AD, as well as PD. KDS12025 can counteract the decline of antioxidative functions in neurodegenerative diseases, offering a viable therapeutic strategy where natural defenses fall short.

Extracellular free Hb has been traditionally viewed as toxic due to its tendency to elicit inflammation and damage through ferroptosis^3,38^. However, emerging reports suggest that endogenous Hb can regulate oxidative stress^6,39^. Our study highlights the antioxidative role of endogenous Hbβ in the hippocampus and SNpc (Fig. 3 and 4) under certain conditions, especially when modulated by enhancers like KDS12025. Gene-silencing results support that the absence of Hbβ increases vulnerability to H_2_O_2_ (Fig. 4). The unexpected antioxidative role of Hbβ proposes brain Hbβ as an effective therapeutic target in neurodegenerative diseases to manage oxidative stress potently and selectively. KDS12025’s therapeutic potential can be further expanded to relieve H_2_O_2_-mediated neurodegeneration and oxidative stress in various diseases such as ischemic stroke, spinal cord injury, inflammation, and even aging, which is associated with a reduction in brain Hb levels^12^. Furthermore, because Hb is abundantly present in the blood, systemically administered KDS12025 can target the blood Hb initially. In support of this idea, the administration of erythropoietin, a hematopoietic cytokine that can upregulate Hb in red blood cells, has been shown to exert protective effects in the heart, the vasculature, and the nervous system, including AD and PD^5,40,41^. Moreover, our experiments show that LPS-induced systemic inflammation significantly increases blood H_2_O_2_ levels, and KDS12025 treatment can effectively lower this inflammation-induced H_2_O_2_ elevation (Extended Data Fig. 13n-p). Future investigations should be carried out to examine KDS12025’s efficacy in diseases related to peripheral or systemic H_2_O_2_ by targeting the blood Hb.

In summary, we have delineated the unprecedented antioxidative function of Hb, which can be boosted by KDS12025. The newly developed tools and concepts in this study should be useful in overcoming various brain and peripheral diseases whose etiology involves oxidative stress.

## Acknowledgments

We express our sincere thanks to Yejin Cho, Juyeon Chae, Suyeon Yellena Kim, and Eunjin Shin for providing technical assistance in producing AAV viruses, optimizing enzyme assays, and capturing Lattice SIM images. We also extend our gratitude to the Research Solution Center (RSC) at the Institute for Basic Science (IBS) for their management of the animal facilities and for providing access to Imaris software. Special thanks to Dr. Andre Berndt from the University of Washington for generously supplying the oROS-G plasmid. Additionally, we acknowledge BioRender.com for facilitating the creation of some of the illustrative images presented in our work.

## Funding

This work was supported by the following agencies: the Institute for Basic Science (IBS), Center for Cognition and Sociality (IBS-R001-D2 to C.J.L.); the Korea Health Technology R&D Project through the Korea Health Industry Development Institute (KHIDI) and Korea Dementia Research Center (KDRC), funded by the Ministry of Health & welfare and Ministry of Science and ICT, Republic of Korea (HU23C0018 to K.D.P.; RS-2023-KH137130 to H.R.), and the Korea Institute of Science and Technology (KIST) Institutional Program (2E32851 to K.D.P.); the National Research Foundation (NRF) grant funded by the Korea government (MSIT) (RS-2023-00261784 to A.N.P.; 2022R1A2C3013138 to H.R.). Supercomputing resources were provided by the Korea Institute of Science and Technology Information (KISTI-HPC) (KSC-2023-CRE-0420 to A.N.P.).

### Author contributions

Conceptualization: WW, EHL, HC, KDP, CJL

Methodology: WW, EHL, LG, ANP, KDP, CJL

Investigation: WW, EHL, LG, HC, DK, MB, JC, TK, UP, JL, SO, CJW, YK, SJP

Funding acquisition: ANP, KDP, CJL

Supervision: HR, ANP, KDP, CJL

Writing: WW, EHL, LG, ANP, KDP, CJL

## Competing interests

W.W., E.H.L., K.D.P., C.J.L. as well as IBS and KIST, are inventors on a patent for the novel aromatic compounds (KR-10-2643543-0000) and have a pending PCT application (PCT/KR2021/016542). The other authors have no competing interest to declare.

## Data and materials availability

All data are available in the main text or the supplementary materials.

Supplementary Information is available for this paper.

---

**Extended Data Fig. 1.**
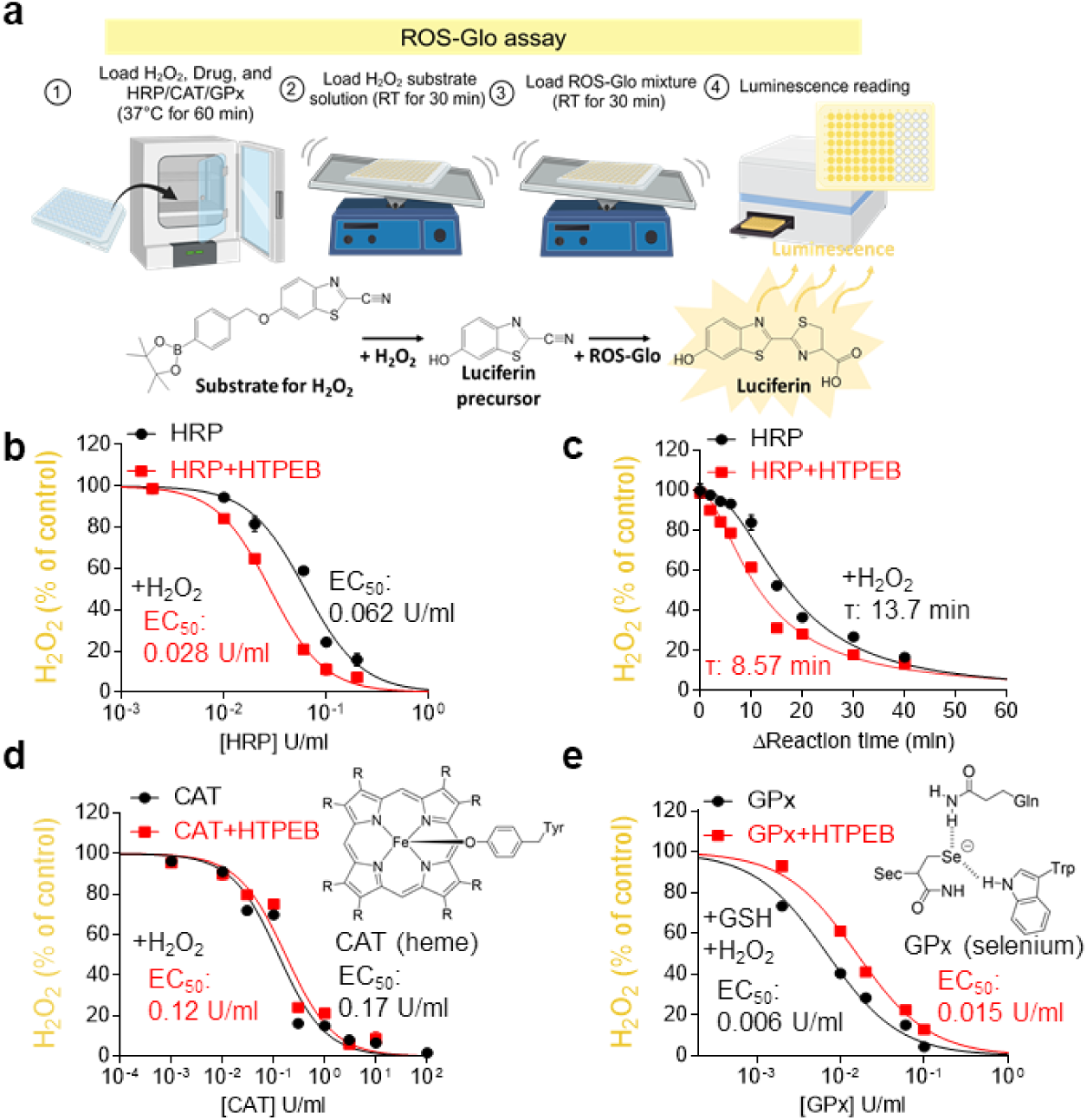
H_2_O_2_ assays using various peroxidase families. To test whether HTPEB enhances H_2_O_2_-decomposition with HRP, CAT, and GPx, we used ROS-Glo H_2_O_2_ assay. **a,** Schematic diagram of ROS-Glo H_2_O_2_ assay. **b,** Dose-response curve of HRP in ROS-Glo assay with (red square shapes) or without (black circle shapes) an HTPEB (10 μM). **c,** Reaction time-response curve for HRP with (red square) or without (black circle) HTPEB. **d,** Dose-response curve of CAT (heme as a cofactor) in ROS-Glo assay with (black circle) or without (red square) an HTPEB. **e,** Dose-response curve of GPx (selenium as a cofactor) in ROS-Glo assay with (black circle) or without (red square) an HTPEB. These results indicate that HTPEB enhances the H_2_O_2_-decomposing activity of the heme-containing protein like HRP, not the selenium-containing protein. The dose-response curve and EC_50_ were calculated and determined by fitting data with GraphPad Prism software. Data are presented as the mean ± s.e.m.

**Extended Data Fig. 2.**
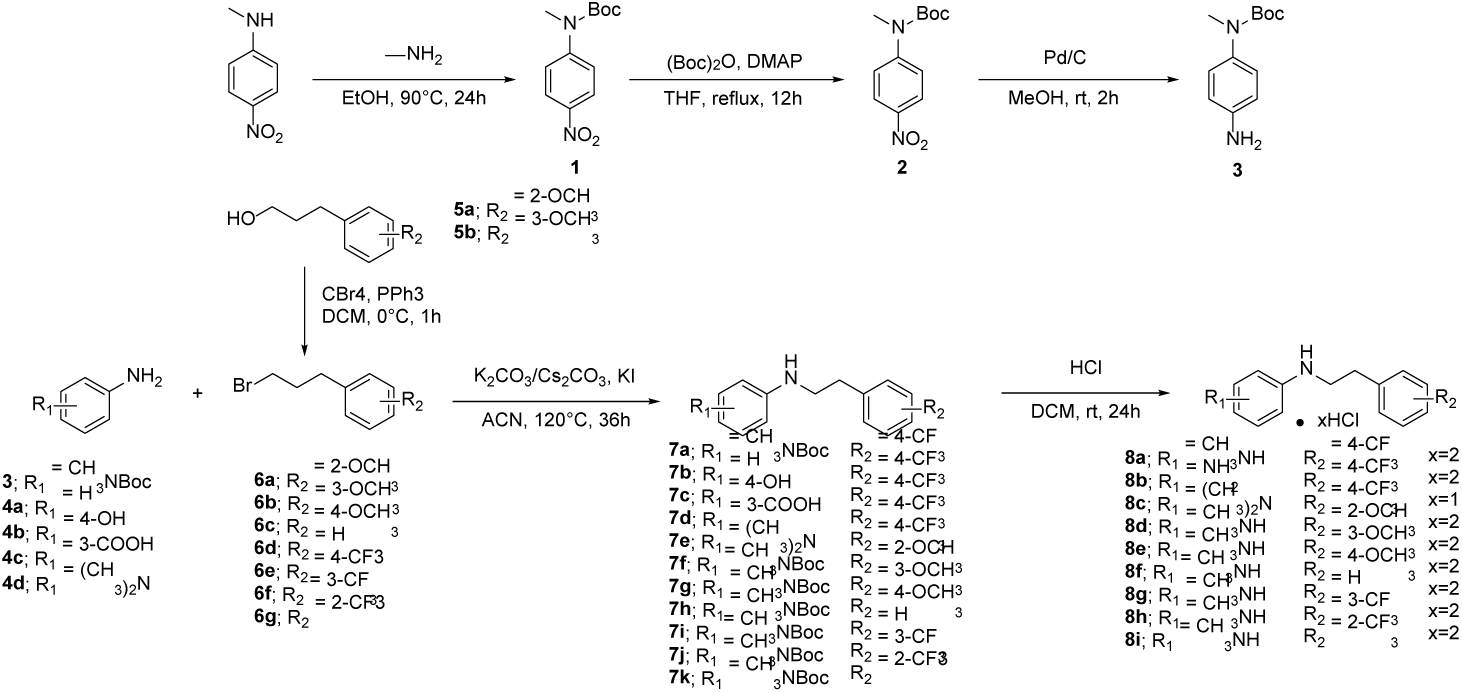
General Procedure for preparation of KDS compounds. Synthetic procedures for preparation of intermediates and KDS compounds.

**Extended Data Fig. 3.**
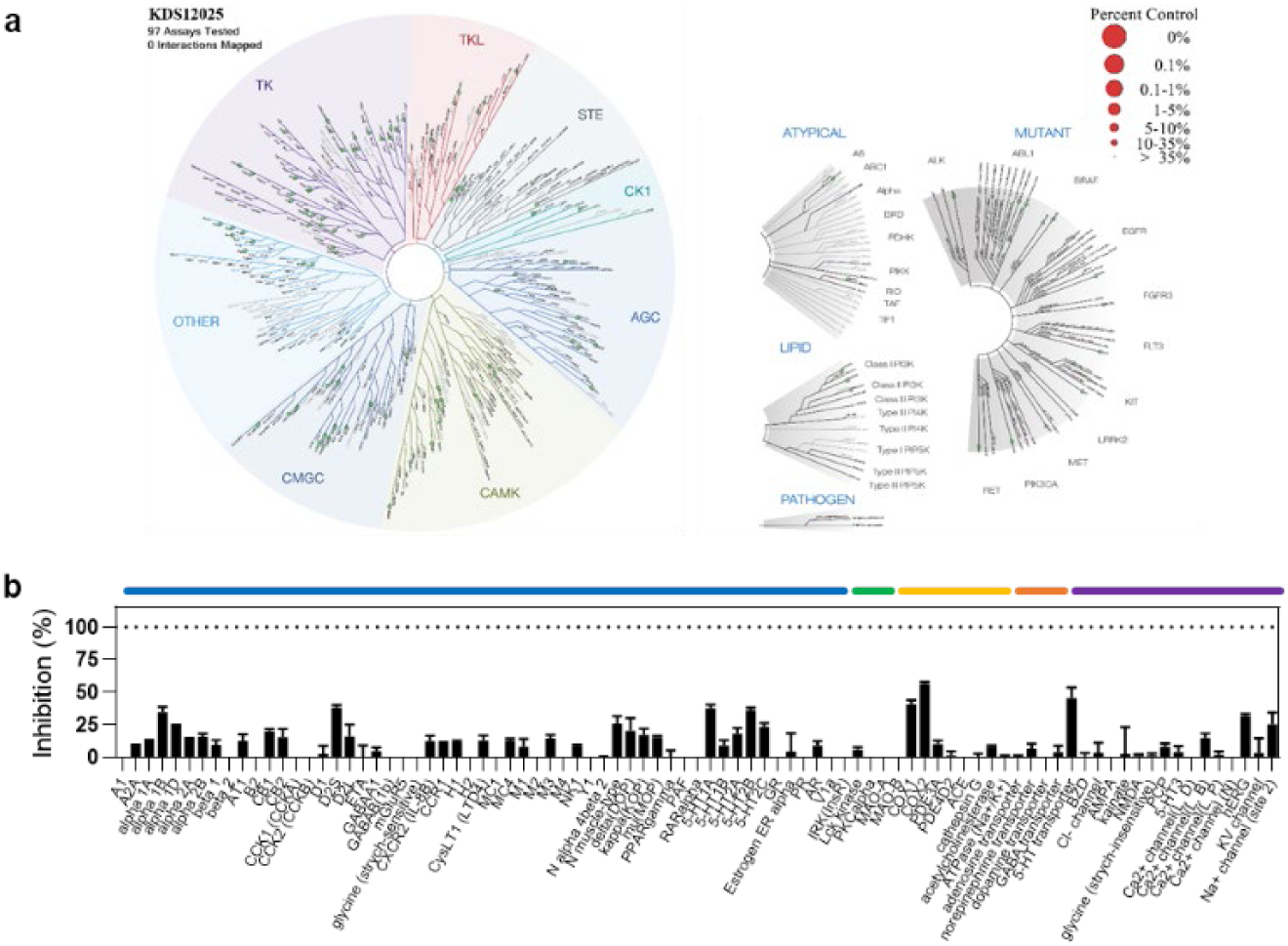
KINOMEscan and Off-target Selectivity Screening Results for KDS12025 at 1000 nM. To test the off-target of KDS12025, we employed screening assays. **a,** Schematic diagram of KINOME*scan*^TM^ screening results categorizing human kinases and disease-associated mutant variants. Competitive binding assays for 97 human kinases were performed at 1000nM KDS12025, and the amount of inhibition through the control ligand reaction is expressed in the size of the colored circle (green/red). Zero interactions mapped means no meaningful responses with ≥ 50% inhibition. TK, Tyrosine Kinase; TKL, Tyrosine Kinase Like; STE, Yeast STE-MAPK family; CK1, Casein Kinase 1; AGC, PKA, PKG, PKC family; CAMK, Calmodulin/Calcium regulated kinases; CMGC, CDK, MAPK, GSK3 and CLK; see Supplementary Table 4 for detailed results. **b,** The off-target selectivity for 87 primary molecular targets at 1000 nM of KDS12025, including G protein-coupled receptors (blue line), kinases (green line), non-kinase enzymes (yellow line), transporters (orange line), and various channels (purple line); see Supplementary Tables 5 for detailed results. These results indicate that KDS12025 shows favorable drug-like properties.

**Extended Data Fig. 4.**
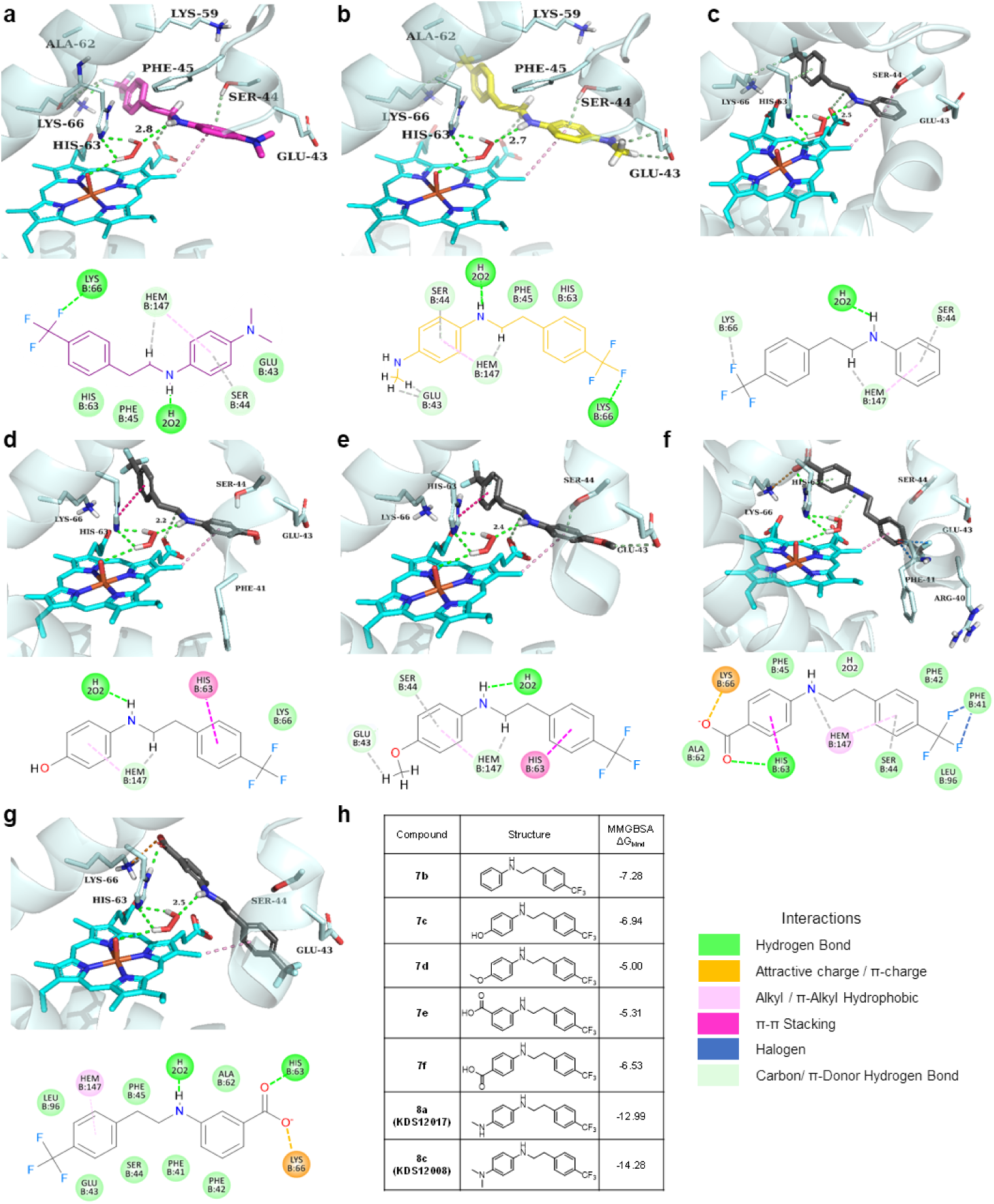
Binding modes of selected *N*-phenethylaniline compounds in Hbβ chain. To investigate whether enzyme assay results and computational modeling were correlated with each other. **a-g,** Predicted 3D binding mode and 2D interaction map of compounds KDS12017 (**a**), KDS12008 (**b**), 7b (**c**), 7c (**d**), 7d (**e**), 7f (**f**) and 7e (**g**) in the hemoglobin β-subunit. **h,** Binding energy (ΔG_bind_) values for corresponding compounds, calculated by the MMGBSA method. These results support the SAR analysis. Enzyme assay results are displayed in Supplementary Table 2, and The calculated binding energy (ΔGbind) values for each compound binding event are specified at the bottom center.

**Extended Data Fig. 5.**
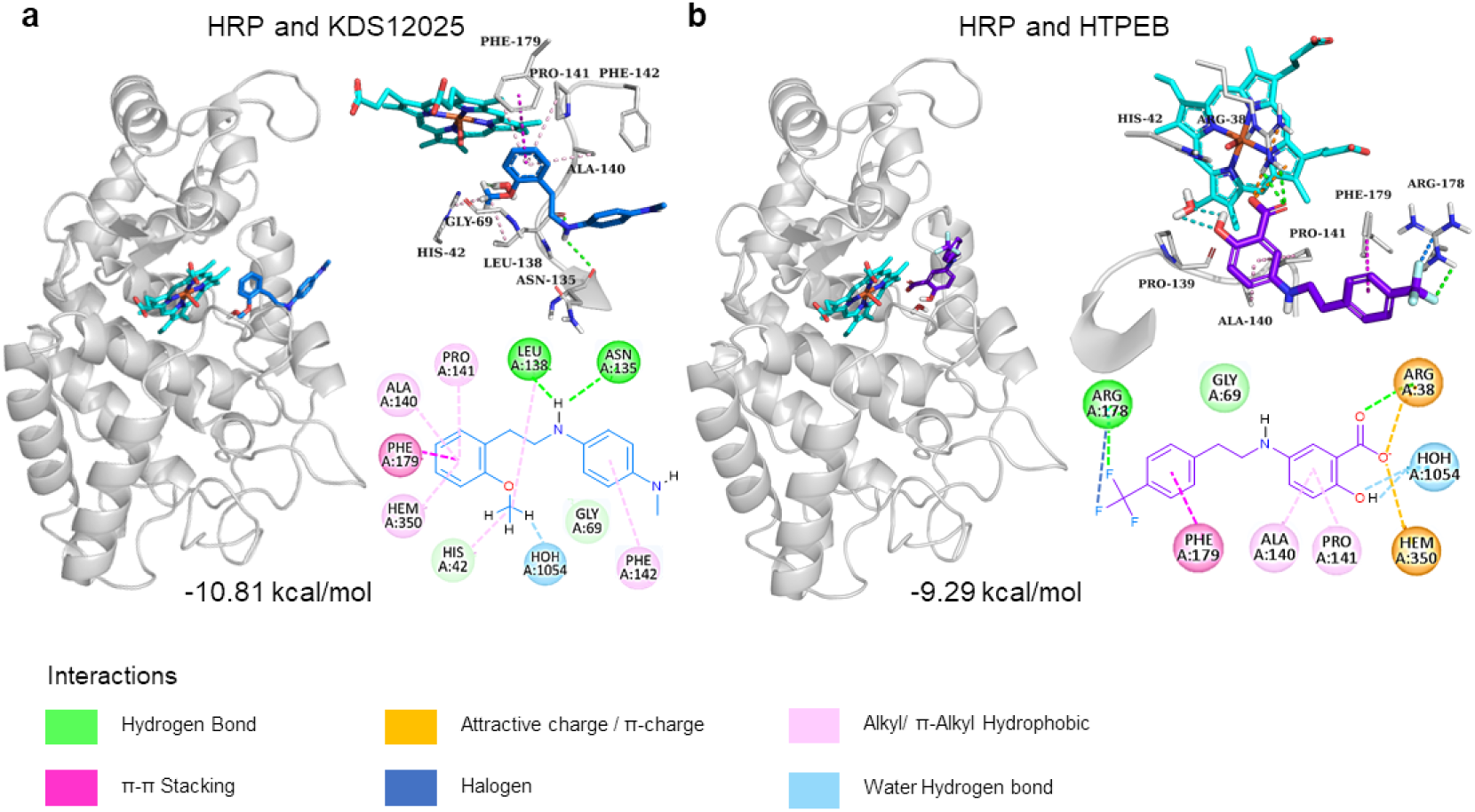
Binding modes of selected active compounds in HRP proposed by molecular docking. **a,** General view of protein-ligand binding (left), 3D interaction scheme (top right), and 2D interaction map (bottom right) of KDS12025. **b,** General view of protein-ligand binding (left), 3D interaction scheme (top right), and 2D interaction map (bottom right) of HTPEB. The calculated binding energy (ΔG_bind_) values for each compound binding event are specified at the bottom center.

**Extended Data Fig. 6.**
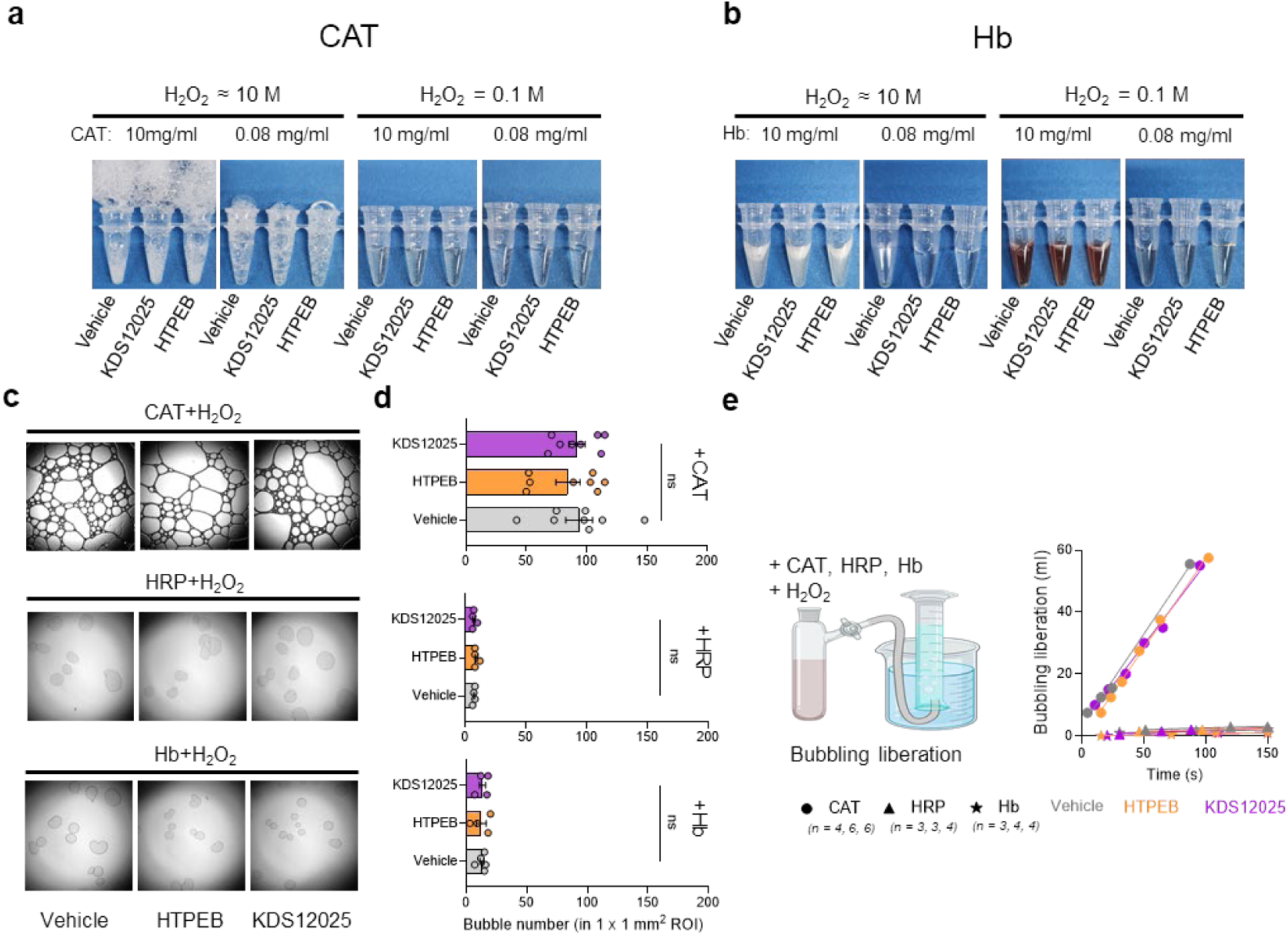
Lack of oxygen liberation by Hb. To test whether Hb might also function as CAT, which decomposes H_2_O_2_ into oxygen and water, while also evaluating the role of KDS12025 in this biochemical reaction. **a,** The assay shows oxygen release from CAT under varied conditions, with observable changes that indicate the degree of oxygen liberation. **b,** The assay shows oxygen release from Hb under varied conditions, with observable changes indicating oxygen liberation. **c,** Observing bubbles generated from CAT, HRP, and Hb in reaction with H_2_O_2_ and drugs (vehicle, HTPEB, KDS12025) under a microscope. **d,** Count the bubbles generated from CAT, HRP, and Hb per 1 x 1 mm2 area. **e,** Schematic images of bubble liberation measurement from CAT, HRP, and Hb reaction with H_2_O_2_ (left) and quantified bubble liberation volume with the cylinder (right). These results demonstrate that Hb is not a catalase, and KDS12025 does not facilitate CAT activity. Data are presented as the mean ± s.e.m. ns, not significant. Additional statistics are provided in Supplementary Table 7.

**Extended Data Fig. 7.**
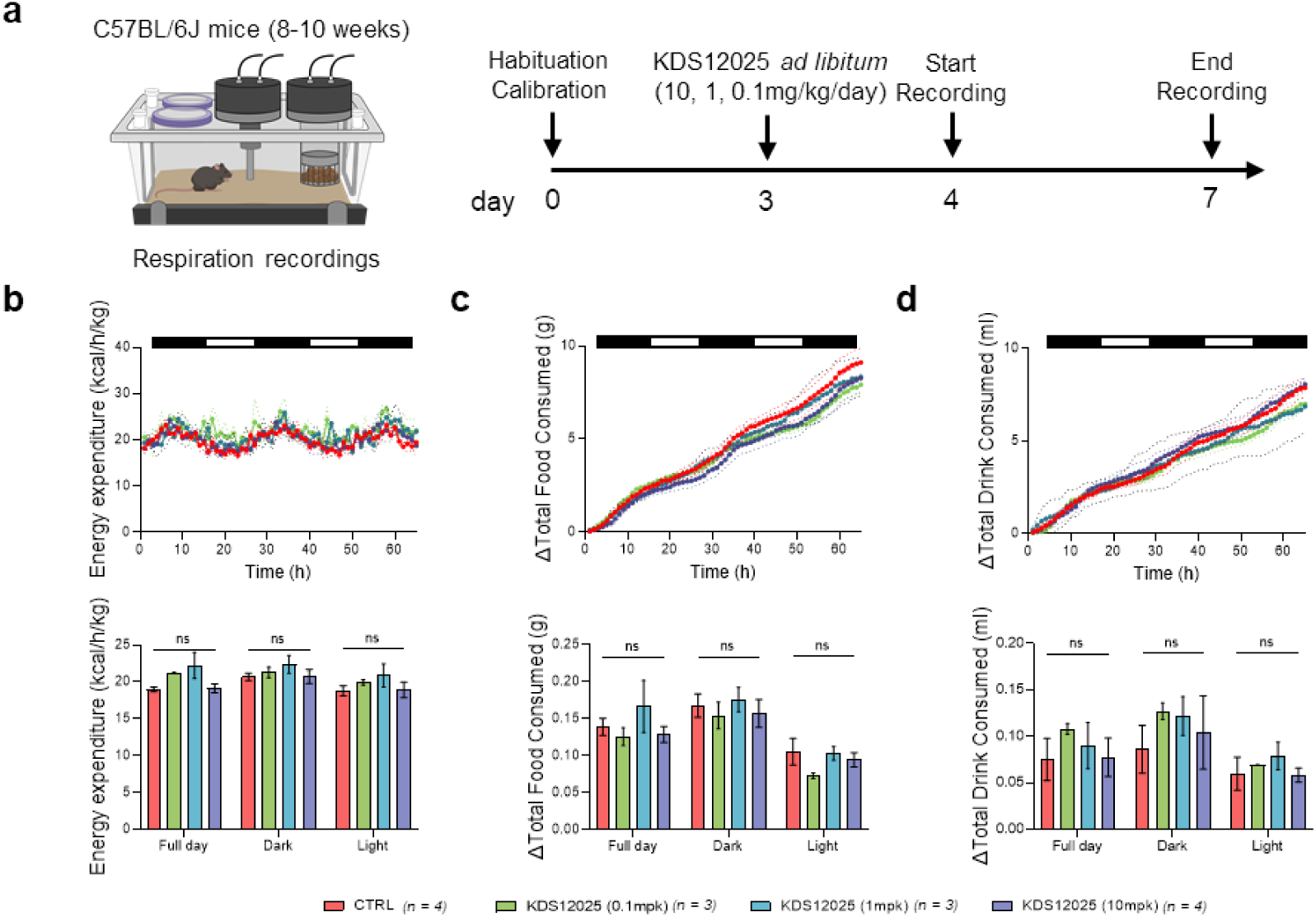
KDS12025 does not alter metabolic processes. To assess whether KDS12025 affects mice’s energy expenditure, food, or drink consumption, we employed the PhenoMaster to measure in the mouse chamber. **a,** Timeline of PhenoMaster experiments with the administration of KDS12025 (0.1, 1, 10 mg/kg/day). **b-d,** Measurement of energy expenditure (**b**), total food consumption (**c**), and total drink consumption (**d**) during the administration of KDS12025 at concentrations of 10, 1, and 0.1 mg/kg/day for both night (dark) and day (white) cycles, suggesting that KDS12025 enhances Hb’s peroxidase activity without overall metabolism in mice. Data are presented as the mean ± s.e.m. ns, not significant. Additional statistics are provided in Supplementary Table 7.

**Extended Data Fig. 8.**
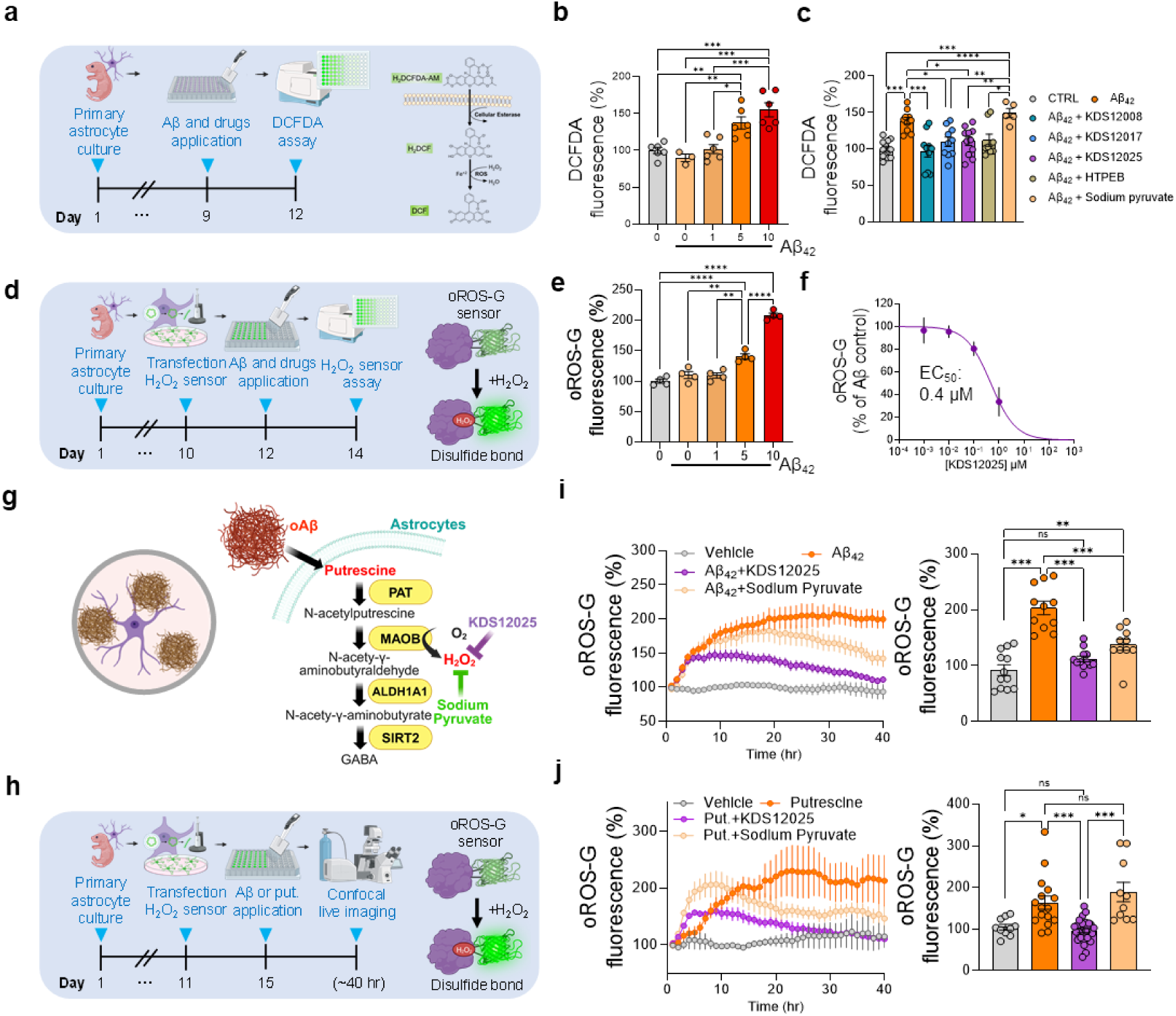
KDS12025 reduces aberrant H_2_O_2_ in astrocytes. We tested whether the KDS12008, 17, and 25 enhance the H_2_O_2_-decomposing peroxidase activity in cultured astrocytes with H_2_O_2_ dye and sensor. **a,** Timeline and schematic diagram of DCFDA-based H_2_O_2_ assay (left) and H_2_O_2_ assay in cultured astrocytes at various concentrations of Aβ_42_ (right). **b,** DCFDA assay in cultured astrocytes at various concentrations of Aβ_42_ (0, 1, 5, 10 μM). **c,** Measurement of Aβ-induced H2O2 levels in cultured astrocytes after treating KDS12008 (10 µM), KDS12017 (10 µM), KDS12025 (10 µM), HTPEB (10 µM), or sodium pyruvate (1 mM). **d,** Timeline and schematic diagram of newly developed H_2_O_2_-selective cpGFP sensor, oROS-G. **e,** oROS-G assay in cultured astrocytes at various concentrations of Aβ_42_ (0, 1, 5, 10 μM). **f,** Dose-response curve of KDS12025 within Aβ_42_ (5 µM) incubated astrocytes. **g,** Schematic diagram of Aβ- or putrescine (a precursor of MAO-B-dependent H_2_O_2_ production)-induced aberrant H_2_O_2_ in astrocytes. **h,** Timeline of 40-hour live H_2_O_2_ imaging using oROS-G probe in Aβ or putrescine incubated cultured astrocytes. **i,** 40-hour continuous H_2_O_2_ imaging using oROS-G probe following oligomerized Aβ (5 μM) treatment, with the administration of KDS12025 (10 μM) and sodium pyruvate (1 mM). **j,** Results of 40-hour continuous H_2_O_2_ measurement following putrescine (180 μM) treatment, with the administration of KDS12025 and sodium pyruvate. These results indicate that KDS12025 reduces Aβ-induced H_2_O_2_ levels to the control level in vitro within 24 hours. Data are presented as the mean ± s.e.m. *P < 0.05, **P < 0.01, ***P < 0.001; ns, not significant. Additional statistics are provided in Supplementary Table 7.

**Extended Data Fig. 9.**
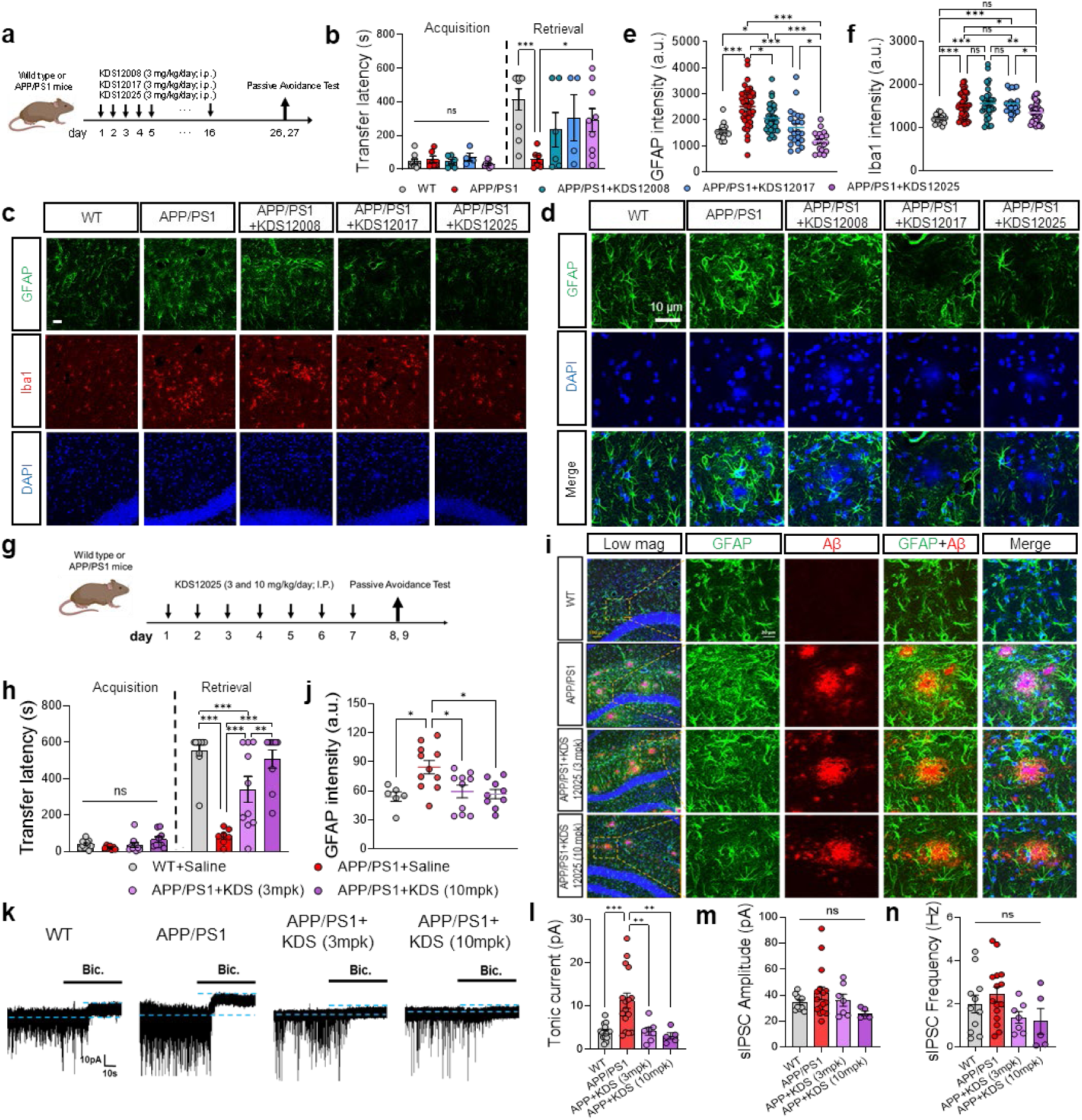
KDS12025 effectively alleviates AD pathology *in vivo.* To further investigate whether the KDS12008, 17, and 25 could alleviate AD symptoms like astrogliosis and memory impairment in the APP/PS1 mouse model. **a,** Schematic timeline of comparison KDS12008, 17, and 25 (3 mg/kg/day intraperitoneal; i.p. injection) in APP/PS1 mice. **b,** Transfer latency to enter the dark chamber where the day had a foot shock during the acquisition session in PAT. **c,d,** Representative image for GFAP, Iba1, and DAPI after administration of KDS12008, 17, and 25. Scale bar, 20 μm. **e,f,** Mean intensity of GFAP (e) and Iba1 (f) in the hippocampus. **g,** Schematic timeline of 3 and 10 mg/kg/day i.p. injection of KDS12025. **h,** Transfer latency to enter the dark chamber. **i,** Representative immunostaining image for GFAP and Aβ after KDS12025 treatment (3 and 10 mg/kg/day i.p. injection). **j,** Mean intensity of GFAP in the hippocampus. **k,** Representative trace of GABA_A_ receptor-mediated tonic GABA current, revealed by the antagonist bicuculline from the dentate gyrus granule cells of the hippocampus. **l,** Measurement of tonic GAB current from the granule cells. **m,n,** Amplitude (m), and frequency (n) of sIPSC measured from granule cells. Data are presented as the mean ± s.e.m. *P < 0.05, **P < 0.01, ***P < 0.001; ns, not significant. Additional statistics are provided in Supplementary Table 7. Data are presented as the mean ± s.e.m. *P < 0.05, **P < 0.01, ***P < 0.001; ns, not significant. Additional statistics are provided in Supplementary Table 7.

**Extended Data Fig. 10.**
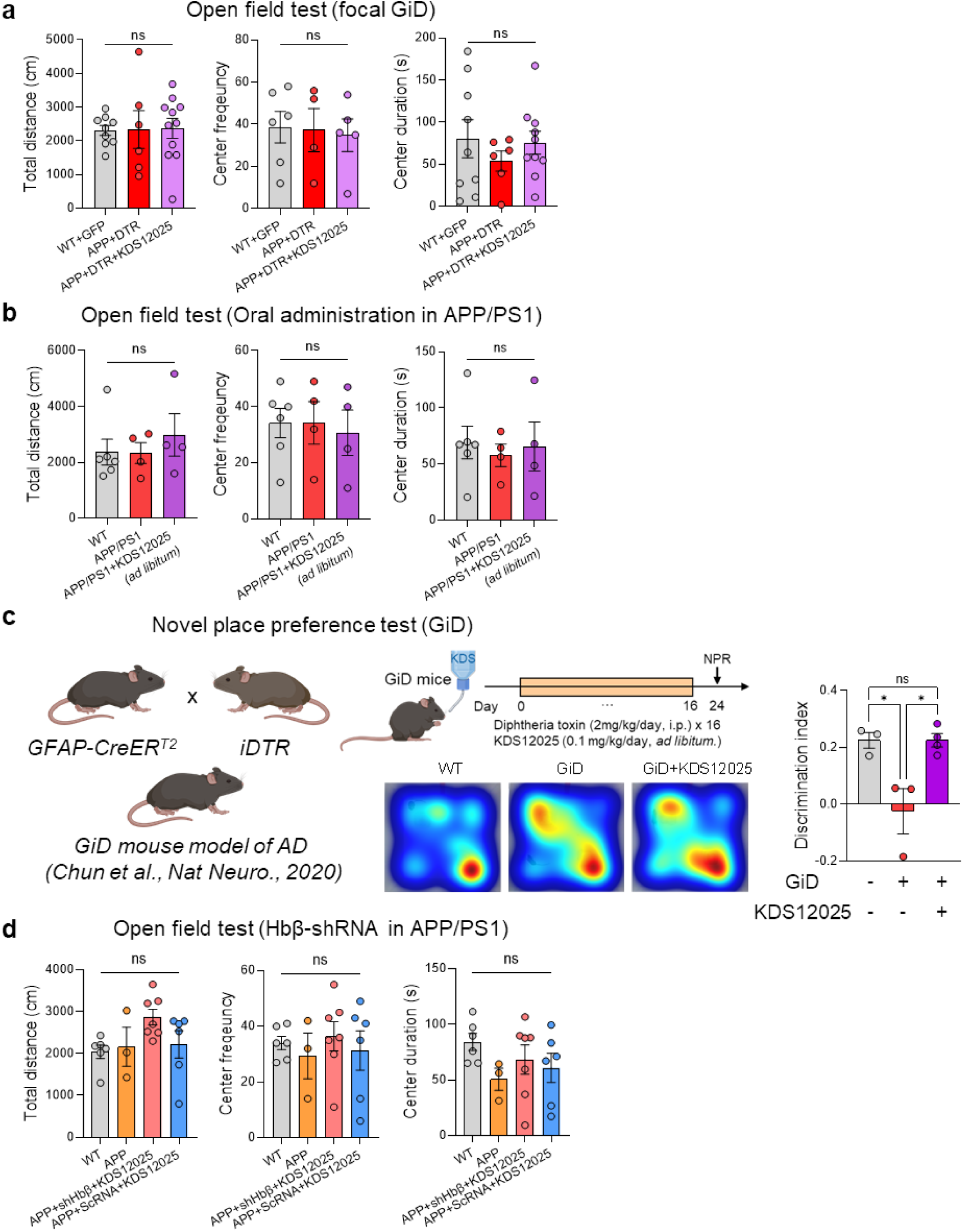
General locomotor and memory performance tests with KDS12025. To investigate whether KDS12025 affects the locomotor activity of APP/PS1 mice. **a,** Open Field Test (OFT) measuring total distance, center frequency, and center duration for focal GiD groups with KDS12025 treatment. **b,** OFT for APP/PS1 groups with KDS12025 (drinking *ad libitum*) treatment. **c,** To assess the extremely low dose of KDS12025 in an AD mouse model, we used the severe AD mouse model. Left: *GFAP-CreERT2* and *iDTR* crossbred GiD mouse model (*Chun et al., Nature Neuroescience, 2020*). Right: Results of NPR test following administration of KDS12025 (drinking *ad libitum*, 0.1 mg/kg/day) to the GiD mouse model. **d,** OFT for APP/PS1 of Hbβ silencing with shRNA with KDS12025 treatment. Data are presented as the mean ± s.e.m. *P < 0.05; ns, not significant. Additional statistics are provided in Supplementary Table 7.

**Extended Data Fig. 11.**
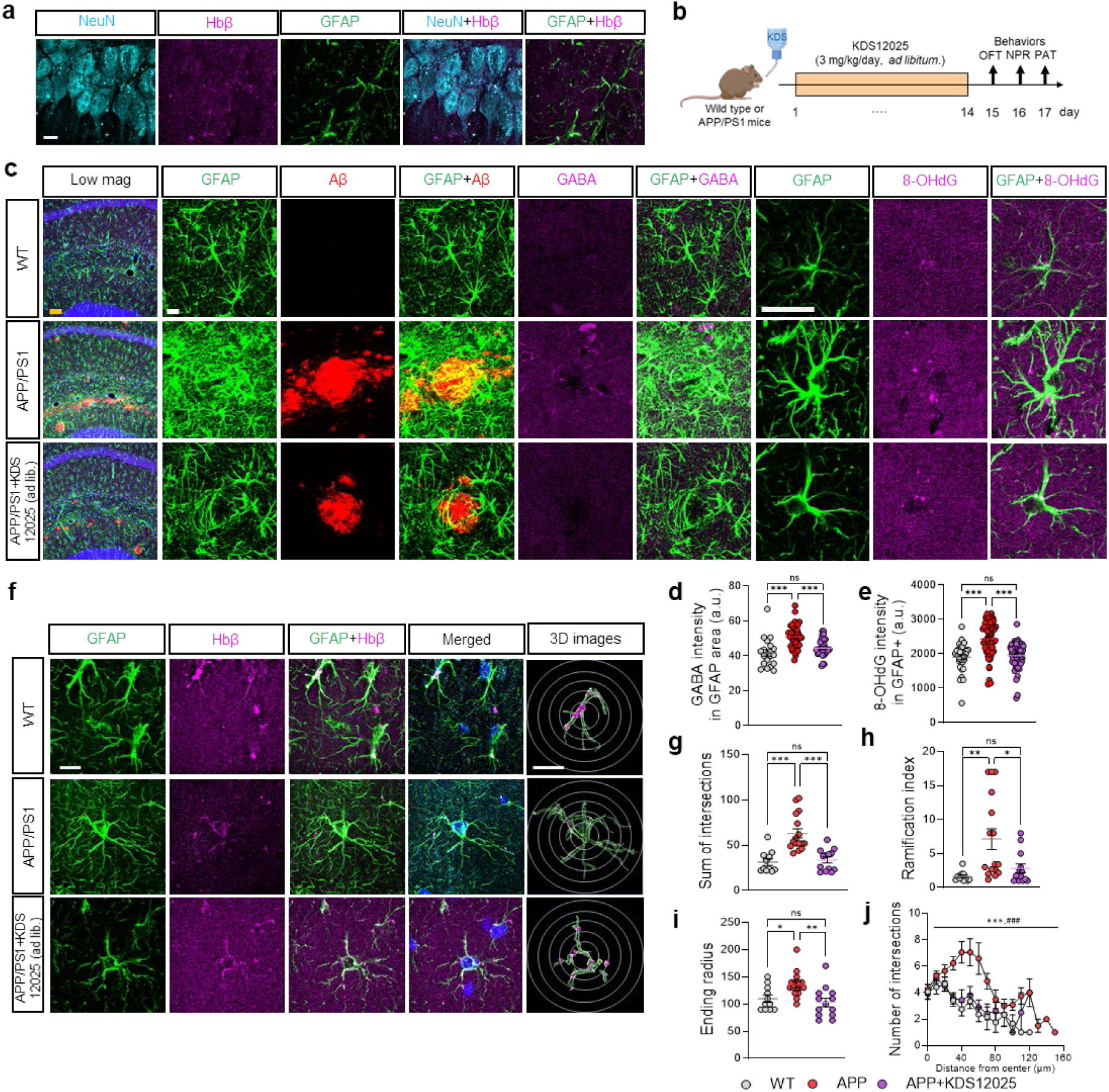
KDS12025 rescues astrocytic Hbβ, GABA, and 8-OHdG expression. **a,** Representative Lattice-SIM images showing NeuN, Hbβ, and GFAP in the pyramidal layer of the hippocampus. Scale bar, 20 μm. **b,** Schematic timeline of APP/PS1 mice and KDS12025 treatment (3 mg/kg/day; drinking *ad libitum*). **c,**Representative images of the hippocampal region for GFAP, Aβ, GABA, 8-OHdG (oxidative stress marker) of WT, APP/PS1, and APP/PS1+KDS12025 (3 mg/kg/day, drinking *ad libitum*). Scale bars, 100 μm (Low mag); 20 μm (High mag); 20 μm (8-OHdG). **d,e,** Mean intensity of GABA (d) and 8-OHdG (e) in GFAP-positive area. **f,** Representative Lattice-SIM images of the hippocampus for GFAP and Hbβ of WT, APP/PS1, and APP/PS1+KDS12025 mice. Representative 3D images from Imaris software (green, GFAP; magenta, Hbβ) and Sholl analysis (circles). Scale bars, 20 μm (main); 10 μm (Imaris). **g,** The summary graph shows the sum of intersections in astrocytes using Sholl analysis. **h,** The summary graph showing the ramification index. **i,** Summary graph showing the ending radius. **j,** The number of intersections relative to their distance from the center. Data are presented as the mean .e.m. ***P .001; ns, not significant. Additional statistics are provided in Supplementary Table 7.

**Extended Data Fig. 12.**
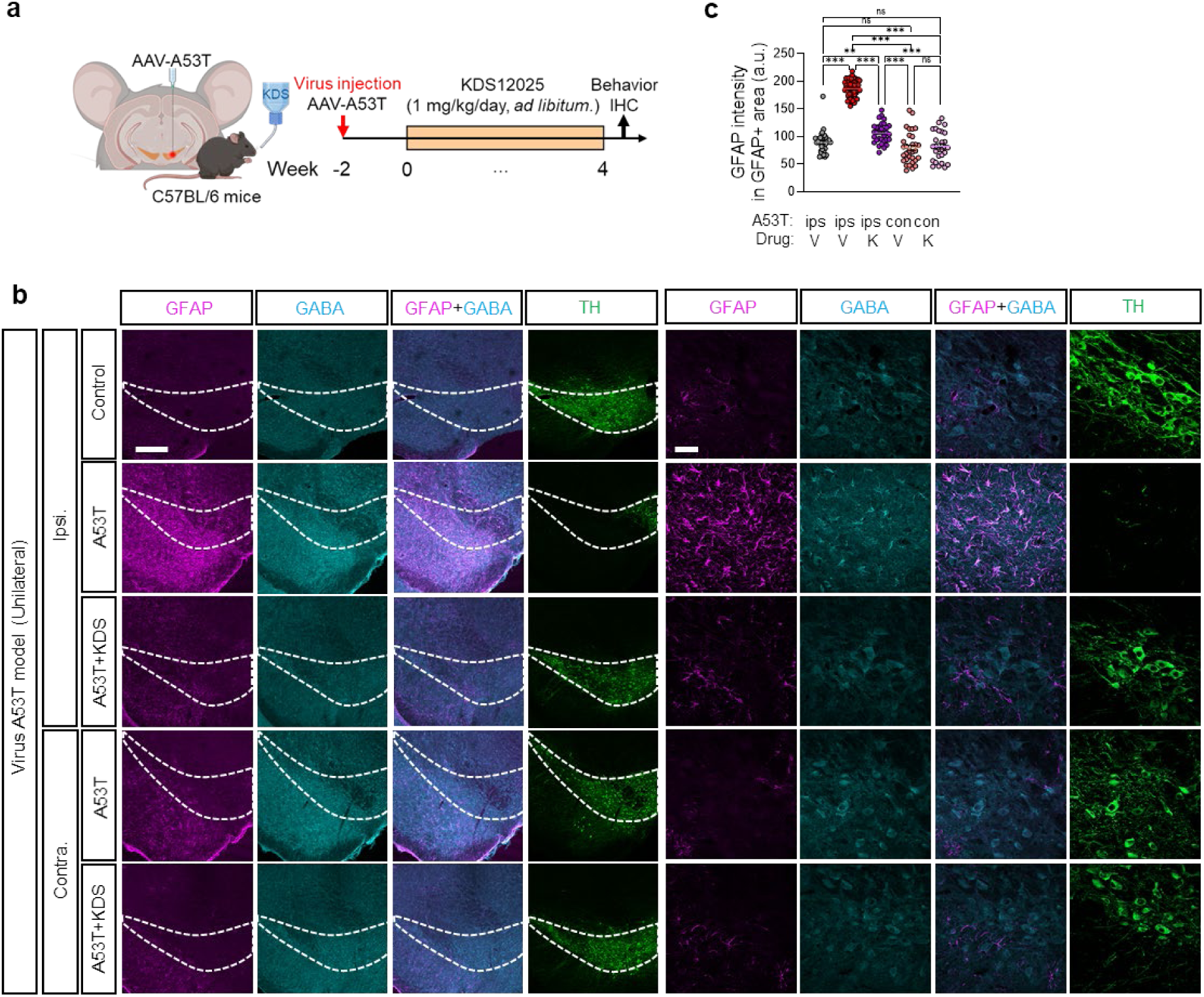
KDS12025 rescues PD pathologies in the A53T mouse model. To examine the effects of KDS12025 in the PD, we used a human A53T overexpression PD mouse model. **a,** Timeline of human A53T-induced PD mouse model (unilaterally overexpression of A53T by injecting AAV carrying human A53T virus) with KDS12025 treatment (1 mg/kg/day for four weeks, drinking *ad libitum*). **b,** Representative images of the SN region with ipsilateral and contralateral side for GFAP, GABA, and TH in control, A53T, and A53T+KDS12025 mice. Scale bars, 150 µm (left); 10 µm (right). **c,** Mean intensity of GFAP in GFAP-positive area. These results demonstrate that KDS12025 rescues the PD pathologies, including astrogliosis, TH-positive neuronal loss, and aberrant astrocytic GABA. Data are presented as the mean ± s.e.m. *P < 0.05, **P < 0.01, ***P < 0.001; ns, not significant. Additional statistics are provided in Supplementary Table 7.

**Extended Data Fig. 13.**
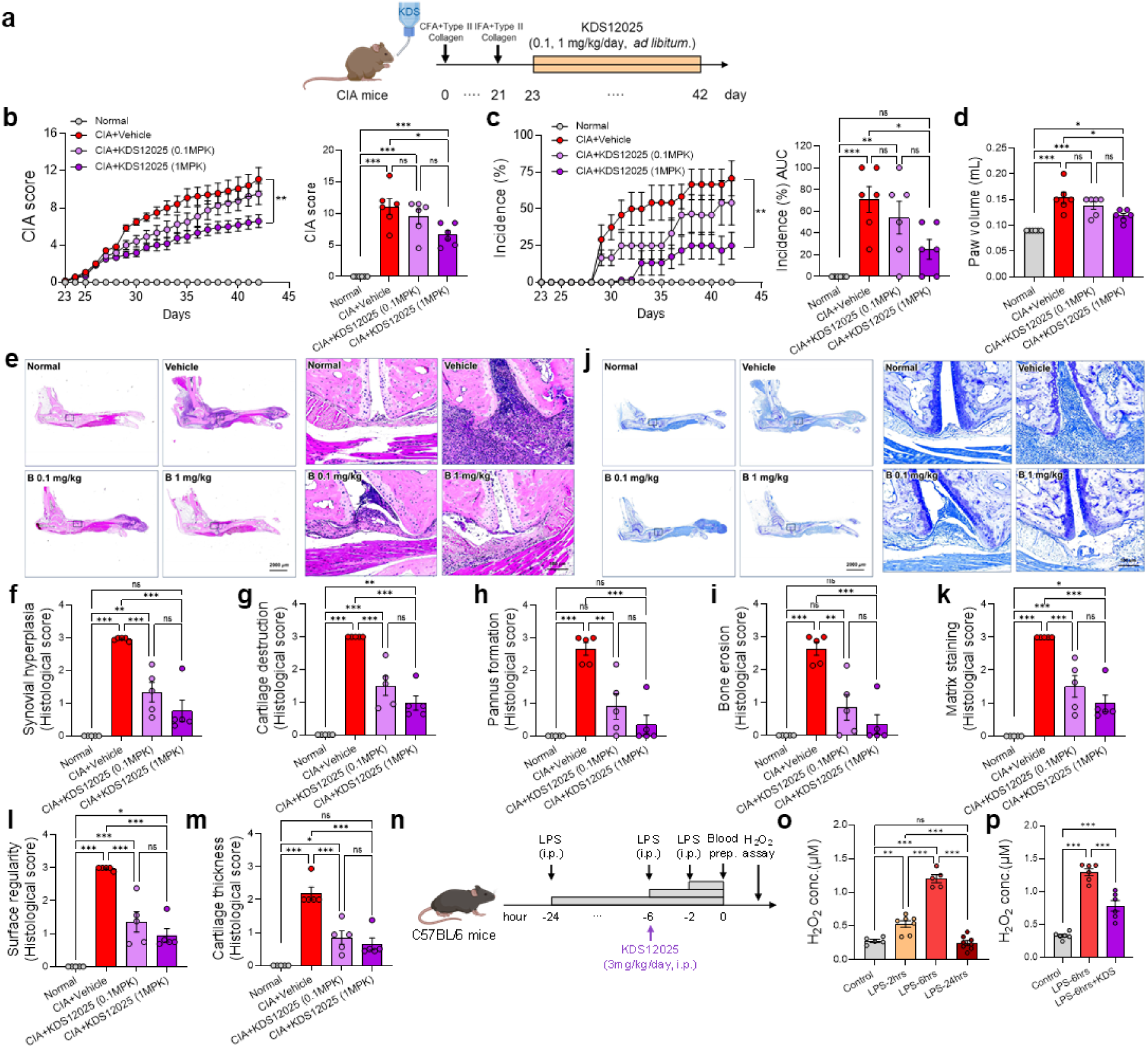
KDS12025 alleviates inflammation-associated RA. To examine the effects of KDS12025 on the inflammatory disease, we used the RA mouse model. **a,b,** Measurement of arthritis score in normal, CIA (collagen-induced arthritis)+vehicle, CIA+KDS12025 (0.1 and 1 mg/kg/day, drinking *ad libitum*) mice (left) and summarized bar graph at day 42 (right). **c,** Incidence of arthritis in normal, CIA+vehicle, CIA+KDS12025 (0.1 and 1 mg/kg/day) mice (left) and summarized bar graph at day 42 (right). **d,** Paw volume (mL) at day 42. **e,** Representative images of Hematoxylin and Eosin (H&E) staining of the joint section in normal, CIA+vehicle, CIA+KDS12025 (0.1 and 1 mg/kg/day) mice. **f-i,** Summarized bar graph showing synovial hyperplasia (f), cartilage destruction (g), pannus formation (h), and bone erosion score (i) by H&E staining analysis. j, Representative image of Safranin O staining of the joint section in normal, CIA+vehicle, CIA+KDS12025 (0.1 and 1 mg/kg/day) mice. **k-m,** Summarized bar graph showing matrix staining (k), surface regularity (l), and cartilage thickness (m) by toluidine blue staining. **n,** To measure the systemic level of H_2_O_2_, we injected an LPS to induce the inflammation acutely. Timeline of LPS (20 mg/kg, i.p. injection)-induced H_2_O_2_ levels of blood from the left ventricle of the heart in mice. Plasma was collected and measured using an Amplex red assay. **o,** Time-dependent LPS (20 mg/kg, i.p. injection, 2, 6, and 24hrs)-induced H_2_O_2_ levels in the blood. **p,** KDS12025 (3 mg/kg, i.p. injection) treatment at 6hrs of LPS (20 mg/kg, i.p. injection) and measurement of blood H_2_O_2_. These results demonstrate that KDS12025 effectively alleviates systemic H_2_O_2_. Data are presented as the mean ± s.e.m. *P < 0.05, **P < 0.01, ***P < 0.001; ns, not significant. Additional statistics are provided in Supplementary Table 7.

**Extended Data Fig. 14.**
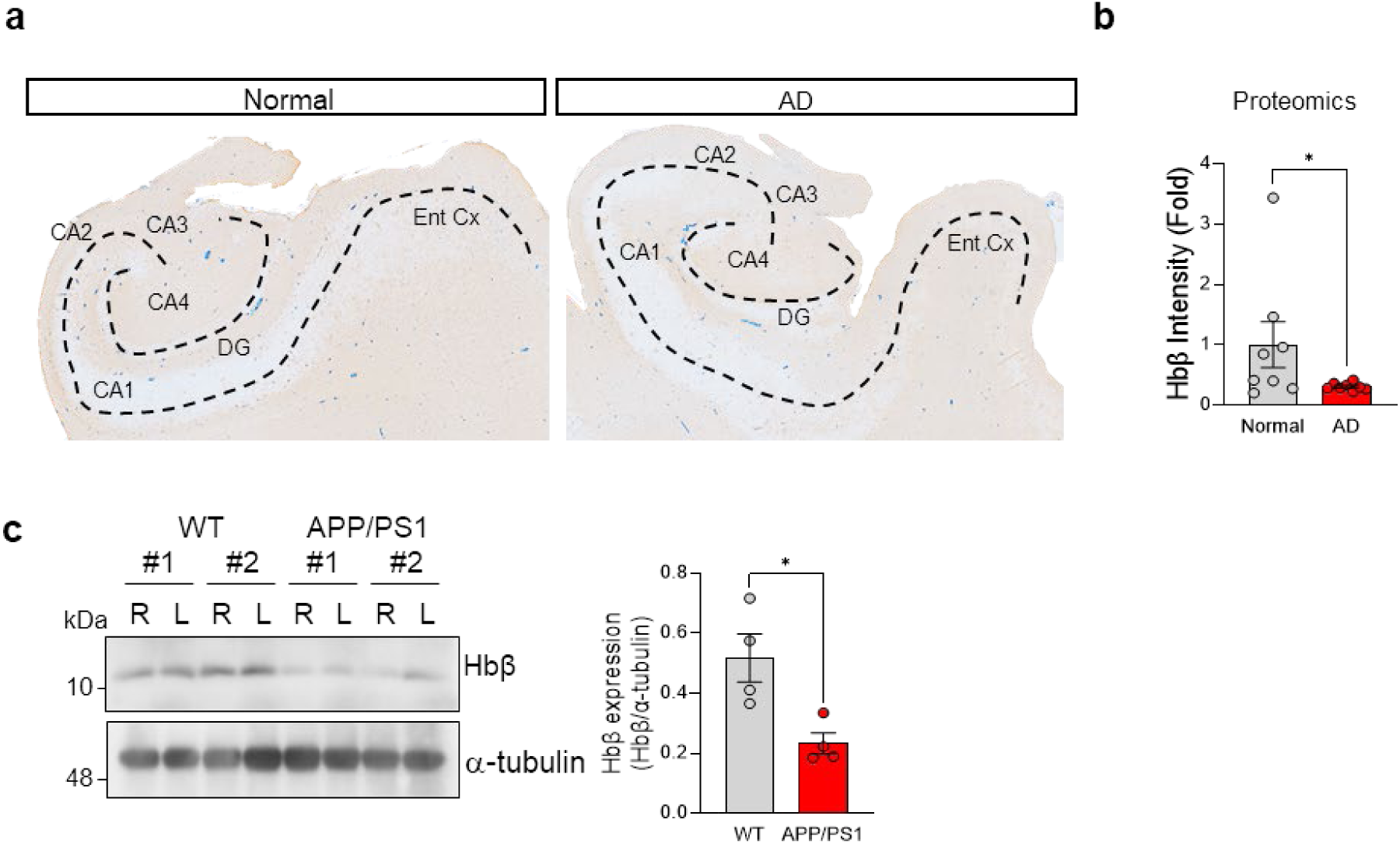
Reduced hippocampal Hbβ in AD patients and mouse model. To assess the clinical significance of the astrocytic Hbβ in AD patients, we performed proteomics on postmortem hippocampal tissues from both normal subjects *(n = 3)* and AD patients *(n = 3)*. **a,** Representative images of postmortem hippocampal tissue from normal subjects and AD postmortem brains. **b,** Hbβ proteomic data of the hippocampal postmortem brain. **c,** Protein quantification of Hbβ protein in the hippocampus of WT littermate and APP/PS1 mice. These findings parallel the reduced Hbβ expression found in the AD mouse model. Data are presented as the mean ± s.e.m. *P < 0.05. Additional statistics are provided in Supplementary Table 7.

**Extended Data Fig. 15.**
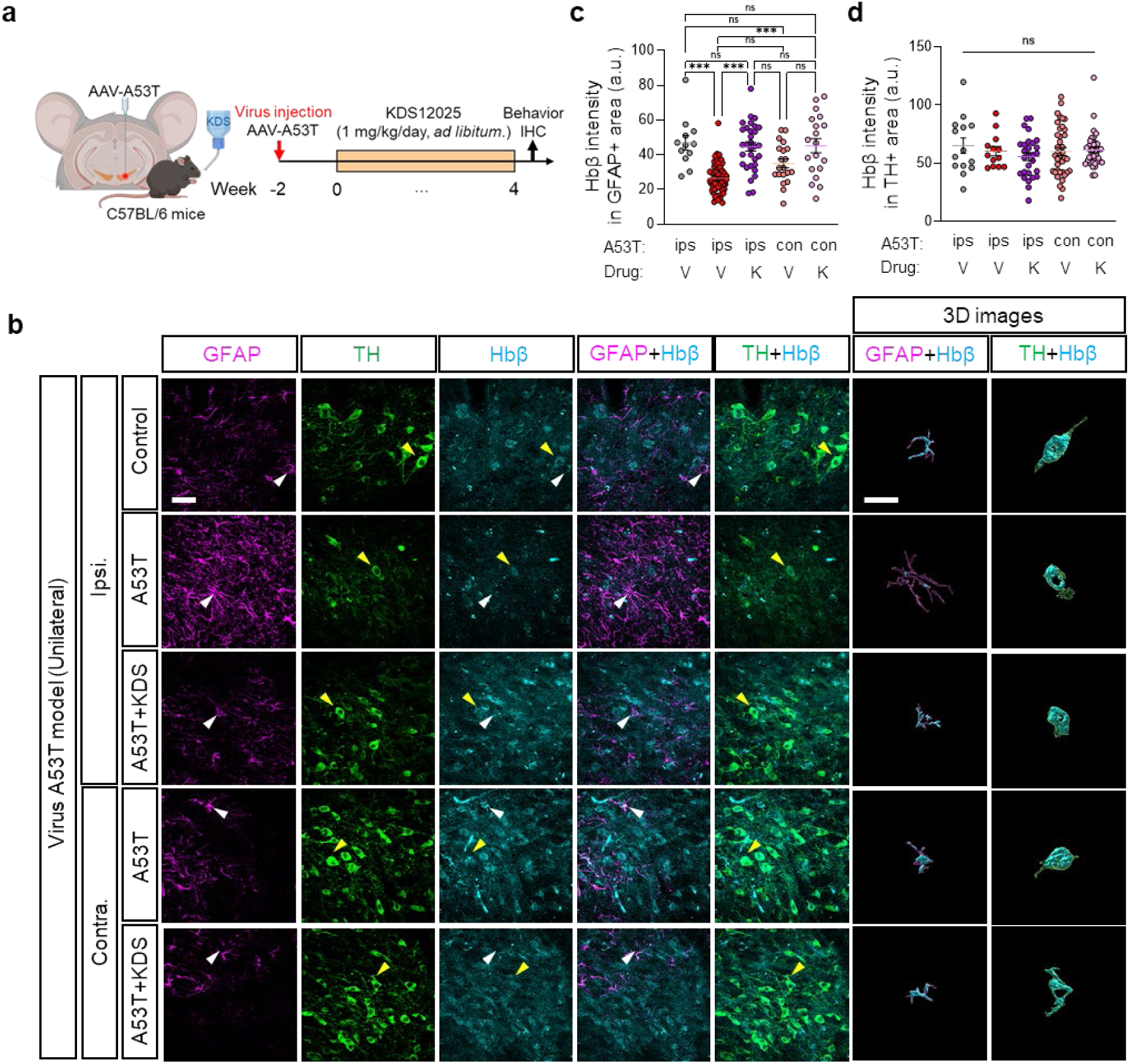
Expression of Hbβ in TH- and GFAP-positive cells in SNpc. To investigate the expression pattern of Hbβ in the PD mouse model, we performed IHC. **a,** Timeline of human A53T-induced PD mouse model (Unilaterally overexpression of A53T by injecting AAV carrying human A53T virus) with KDS12025 treatment (1 mg/kg/day for four weeks, drinking *ad libitum*). **b,** Representative images of the SN region with ipsilateral and contralateral side for GFAP, GABA, and TH in control, A53T, and A53T+KDS12025 mice. Representative 3D images from Imaris software (magenta, GFAP; green, TH; cyan, Hbβ). Scale bars, 20 μm (left); 10 μm (Imaris). **c,d,** Mean intensity of Hbβ in GFAP-positive (**c**) and TH-positive (**d**) area in the SN. Data are presented as the mean ± s.e.m. *P < 0.05, **P < 0.01, ***P < 0.001; ns, not significant. Additional statistics are provided in Supplementary Table 7.

**Extended Data Fig. 16.**
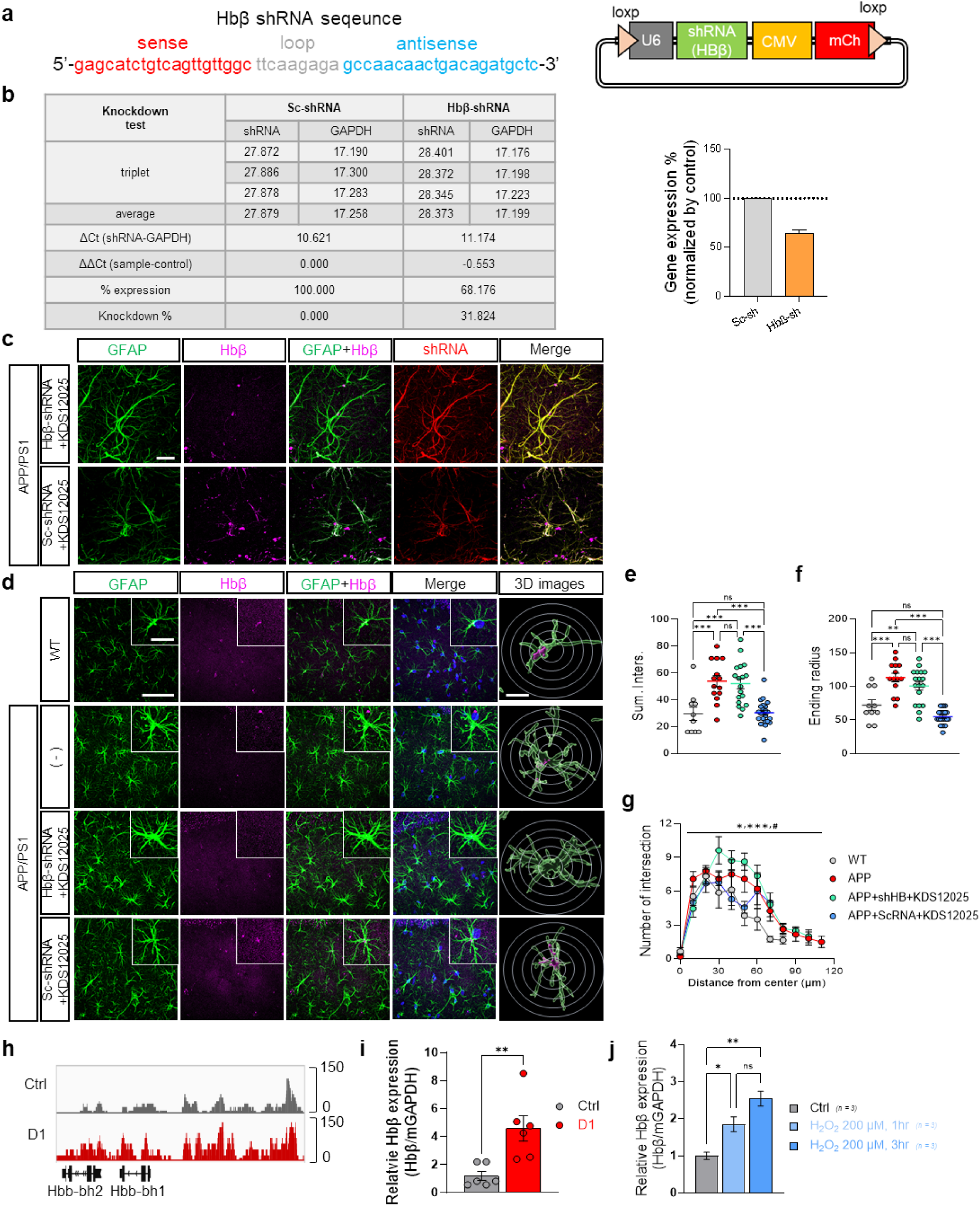
Validation of Hbβ-shRNA and genetic changes in the acute application of oxidative stress in cultured astrocytes. **a,** The sequence and map of Hbβ-shRNA used in this study. **b,** Validation of Hbβ-shRNA efficiency in cultured astrocytes via qRT-PCR, comparing the knockdown results with Sc-shRNA (left), and the bar graph shows the relative expression level. **c,** Lattice SIM imaging demonstrates Hbβ levels in astrocytes after injecting control Sc-shRNA versus Hbβ-shRNA. Scale bar, 20 μm. **d,** Representative images of the hippocampus for GFAP and Hbβ in WT, APP, APP+Hbβ shRNA+KDS12025 (3 mg/kg/day), and APP+Sc-shRNA+KDS12025 (3 mg/kg/day). Representative 3D images from Imaris software. Scale bars, 50 μm (main); 10 μm (inset); 5 μm (Imaris). **e-g,** Summary graph showing the sum of intersections (e), ending radius (f), and number of intersections (g) in astrocytes by Sholl’s analysis. **h,** Representative ATAC-seq tracks for the Hbβ locus accessibility profiles across control (Ctrl, gray) in cultured astrocytes one day after the H2O2 application (D1, red). **i,** Quantitative qRT-PCR measurement of Hbβ mRNA across Ctrl and D1 as in panel (h). **j,** Quantitative qRT-PCR measurement of Hbβ mRNA in cultured astrocytes exposed to 200 µM H_2_O_2_ for 1 hour and 3 hours. Data are presented as the mean ± s.e.m. *P < 0.05, **P < 0.01, ***P < 0.001; ns, not significant. Additional statistics are provided in Supplementary Table 7.

**Extended Data Fig. 17.**
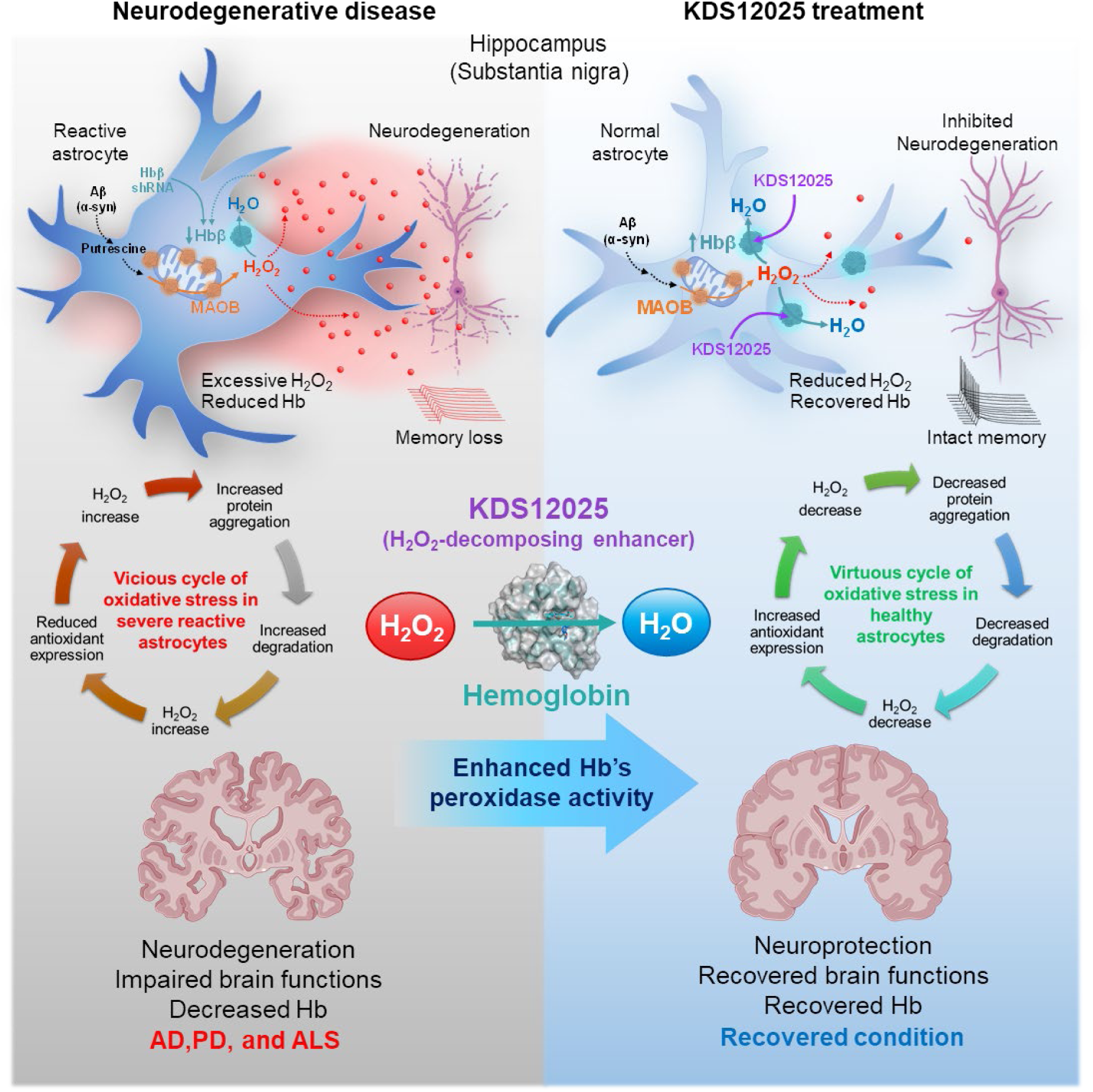
Graphical abstract. **Top left,** Neurodegenerative diseases like AD and PD involve amyloid beta or alpha-synuclein toxins triggering astrocytes to produce putrescine, which MAO-B converts to aberrant H_2_O_2_. Excessive H_2_O_2_ causes neurodegeneration in hippocampal pyramidal neurons or substantia nigra dopaminergic neurons, inhibiting neuronal firing and leading to cognitive and motor impairments. In a chronic H_2_O_2_ exposure environment, a reduction in Hbβ leads to a vicious cycle, vulnerable to oxidative stress and decreasing Hbβ levels, as seen with Hbβ-shRNA. Conversely, the H_2_O_2_-decomposing enhancer KDS12025 promotes brain Hb’s H_2_O_2_-decomposing peroxidase activity, reducing astrocytic H_2_O_2_ levels and normalizing reactive astrocytes. This prevents neurodegeneration, maintains memory and motor functions, and restores Hbβ to normal levels, indicating a return to a virtuous cycle. **Middle,** In the context of neurodegenerative diseases, severe reactive astrocytes initiate a vicious cycle of oxidative stress characterized by increased H_2_O_2_ levels. This aberrant H_2_O_2_ induces protein aggregation, further exacerbating degradation processes leading to even higher H_2_O_2_ levels. Concurrently, antioxidant levels diminish, perpetuating the vicious cycle. Conversely, KDS12025 treatment ceases this vicious cycle by reducing H_2_O_2_ by enhancing Hb’s H_2_O_2_-decomposing peroxidase activity. As a result, H_2_O_2_ levels decrease, protein aggregation is diminished, and antioxidant levels are maintained, thereby establishing a virtuous cycle. **Bottom,** In conclusion, aberrant H_2_O_2_ causes neurodegeneration and brain function impairment in AD, PD, and ALS, with reduced astrocytic Hb. KDS12025 enhances the H_2_O_2_-decomposing peroxidase activity of Hb, mitigating oxidative stress even with extremely low levels of Hb.

**Extended Data Fig. 18.**
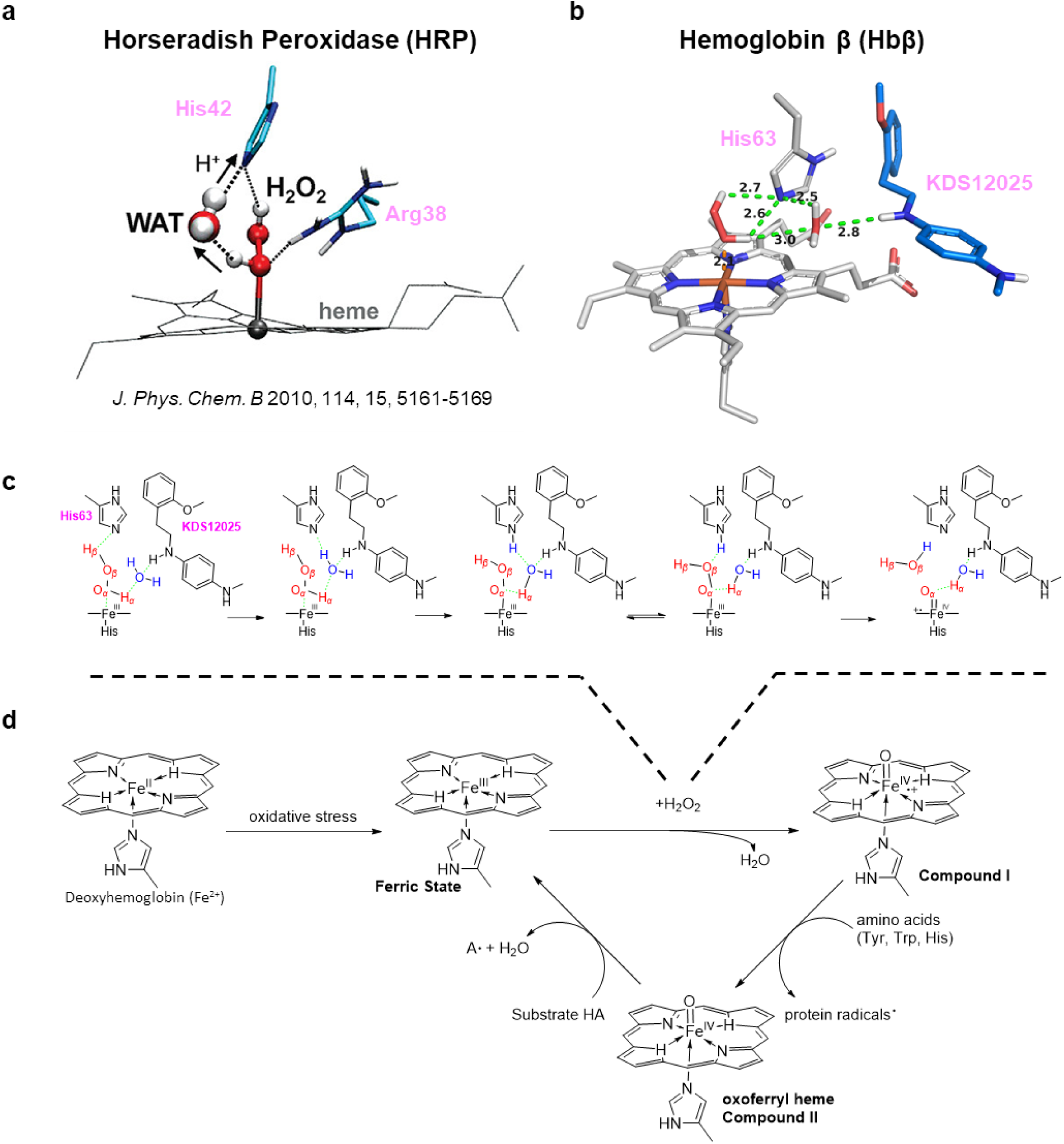
Molecular mechanisms of KDS12025 enhancement of hemoglobin’s peroxidase activity. **a,** Representative snapshot of HRP and H_2_O - H_2_O_2_ – His42 – Arg38 reactive conformation for H_2_O_2_ decomposition and subsequent activation to Por°±Fe^IV^=O (taken from *J. Phys. Chem. B* 2010, 114, 14, 5161-5169). **b,** Predicted reactive conformation of H_2_O – H_2_O_2_ – His63 – KDS12025 for H_2_O_2_ composition and subsequent activation to Por°±Fe^IV^=O (Compound I). **c,** Proposed reactive conformation mechanism for the decomposition of H_2_O_2_ (red color) for proton transfer and release of H_2_O (blue color) with KDS12025 assisting in coordinating H_2_O and subsequently H_2_O_2_ with the iron atom. **d,** General hemoglobin peroxidase cycle. KDS12025 facilitates the decomposition of H_2_O_2_ and the formation of Por°±Fe^IV^=O (Compound I). These results suggest the KDS12025’s novel mode-of-action, which can act like Arg38 in HRP.

## Online Methods

### *In vitro* H_2_O_2_ assay

Hb (Sigma-Aldrich, H7379), H_2_O_2_ (Sigma-Aldrich, H1009), and Amplex Red reagent (Invitrogen™, A12222) were obtained for the study. A 4 μM H_2_O_2_ solution was prepared in a 50 mM sodium phosphate buffer (pH 7.2). Then, 49 μl of this solution was mixed with 1 μl of the test compound in DMSO or distilled water. The mixture was combined with an assay solution of Amplex Red reagent (20 mM stock to 0.1 mM) and HRP (200U/ml stock to 0.2 U/ml) and added to the wells at 50 μl/well. After a 60-minute incubation at 37°C, resorufin production was quantified using a microplate fluorescence reader (SpectraMax iD5, Molecular Devices) at ex/em=535/580 nm.

For the ROS-Glo assay, the H_2_O_2_ substrate solution and ROS-Glo detection solution were prepared according to the manufacturer’s instructions. H_2_O_2_ and drugs were prepared as in the Amplex Red assay, with HRP, Hb, CAT, or GPx added as needed. The mixture was added to the wells at 100 μl/well and incubated for 60 minutes at 37°C. The H_2_O_2_ substrate solution was added and incubated at room temperature for 30 minutes. After adding the H2O2 substrate solution and incubating at room temperature for 30 minutes, the ROS-Glo detection solution was added and incubated for 15 minutes. Luminescence was measured using iD5 microplate readers. The U/ml was calculated based on the amount of H_2_O_2_ reduced by Hb during the reaction, using μg/ml and U/mg.

To evaluate the extent of peroxidase function facilitation by HTPEB (1 μM) and KDS12025 (1 μM) in HRP (0.2 U/ml) and Hb (10 μg/ml), we used the ROS-Glo assay over 30 and 60 minutes. The fold change was calculated by dividing the delta values during the 30-minute reaction for the vehicle over drug conditions.

To assess enzyme activity, the amount of H_2_O_2_ consumed over a specific time was quantified alongside a standard curve. For Hb’s peroxidase function, Hb was administered in doses from 0.01 to 112 U/ml, and pre-incubated with H_2_O_2_ and either DMSO, HTPEB (1 μM) or KDS12025 (1 μM) at 37°C for 30 minutes. The remaining H_2_O_2_ was measured using the ROS-Glo assay normalized against conditions without Hb. The dose-response curve, EC_50,_ and EC_20_ for the H_2_O_2_ decomposition were calculated and determined by fitting data with GraphPad Prism software.

### General Synthetic Method

Reaction progression was checked using analytical thin-layer chromatography (TLC) plates (#1.05715, Merck) and analyzed with 254 nm and 365 nm ultraviolet light. The reaction mixtures were purified by flash column chromatography using silica gel (#1.09385, Merck). Melting points were determined in open capillary tubes using a Standford Research Systems melting point apparatus and were uncorrected. Nuclear magnetic resonance (NMR) spectral data were obtained at 400MHz (^1^H) and at 100MHz (^13^C) using a BRUKER apparatus. Chemical shits (*δ*) were expressed in parts per million (ppm) from tetramethylsilane (TMS), the internal standard and coupling constants (*J*) were expressed in hertz and assigned as follows: s, singlet; d, doublet; t, triplet; q, quartet; AB_q_, AB quartet; br, broad; m, multiplet. All chemical reagents and solvents were of reagent grade, used without further purification, and purchased from commercial sources. Analytical HPLC was performed using a Waters E2695 system with a SHISEIDO capcell pak C_18_ MGII column (4.6mm x 150 mm; 5 μm). HPLC data were recorded using the following parameters: 0.1% acetic acid in H2O/ MeCN, 90/10 -> 0/100 in 10 minutes, +10 minute isocratic hold, flow rate of 1.0 mL/min, λ=254 and 280 nm. Compounds were checked by using TLC, ^1^H and ^13^C NMR and LR-MS. The TLC, NMR, and analytic data confirmed that the purity of the products was ≥ 95%.

### Preparation of *N*-methyl-4-nitroaniline (1)

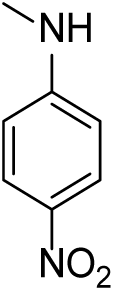

A solution of fluoro-4-nitrobenzene (1.0 g, 7.1 mmol) and methylamine (3.41 g, 109.9 mmol) in 10 mL ethanol was heated at reflux for 24 hours. The reaction was cooled and concentrated in vacuo. The resulting residue was diluted with ethyl acetate and washed with brine, dried (Na_2_SO_4_), and the solvent removed in vacuo to give 913.3 mg (85%) of the title compound as a yellow solid. R*_f_* = 0.70 (*n-*Hex 1: EtOAc 1); ^1^H NMR (400 MHz, DMSO-*d_6_*) *δ* 8.01 (d, 2H, *J* = 9.28), 7.31 (d, 1H, *J* = 4.28), 6.61 (d, 2H, *J* = 9.36), 2.80 (d, 2H, *J* = 5.00), 2.33 (s, 1H).

### Preparation of *tert*-butyl methyl(4-nitrophenyl)carbamate (2)

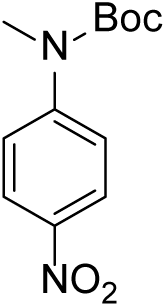

To a solution of *N-*methyl-4-nitroaniline (800 mg, 5.26 mmol) in 10 mL THF were added di-*tert*-butyl dicarbonate (1.72 g, 7.89 mmol) and 4-dimethylaminopyridine (32.1 mg, 0.26 mmol). The reaction mixture was heated at reflux for 12 hours. The reaction was cooled and concentrated in vacuo. The resulting residue was diluted with ethyl acetate and washed with brine, dried (Na_2_SO_4_), and the solvent was removed in vacuo to give 1.29 g (97%) of the title compound as a yellow oil. R*_f_* = 0.80 (*n-*Hex 3: EtOAc 1); ^1^H NMR (400 MHz, DMSO-*d_6_*) *δ* 8.20 (d, 2H, *J* = 9.20), 7.60 (d, 2H, *J* = 9.16), 3.29 (s, 3H), 1.45 (s, 9H)

### Preparation of *tert*-butyl (4-aminophenyl)(methyl)carbamate (3)

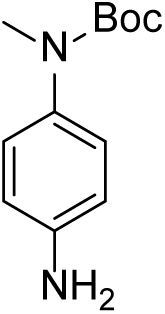

To a solution of *tert*-butyl methyl(4-nitrophenyl)carbamate (6.67g, 26.45 mmol) in 25mL methanol, was added palladium on carbon (667 mg). The reaction mixture was stirred under hydrogen for 2 hours at room temperature. The reaction was filtered through a pad of celite. The filtrate was concentrated in vacuo, to give 5.1768 g (88%) of the title compound as a yellow solid. R*_f_* = 0.05 (*n-*Hex 1: EtOAc 1); ^1^H NMR (400 MHz, DMSO-*d_6_*) *δ* 6.86 (d, 2H, *J* = 8.52), 6.50 (d, 2H, *J* = 8.60), 5.01 (s, 2H), 3.06 (s, 3H), 1.35 (s, 9H)

### Preparation of 1-(2-bromoethyl)-2-methoxybenzene (5a)

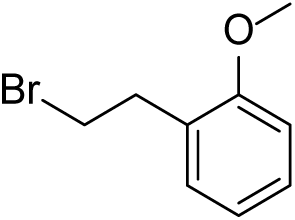

To a solution of 2-(2-methoxyphenyl)ethan-1-ol (5g, 32.85 mmol) in anhydrous dichloromethane (75 mL) was added carbon tetrabromide (15.25 g, 45.99 mmol) and stirred at 0° under nitrogen atmosphere. Triphenylphospine (10.35 g, 39.42 mmol) was added potionwise and stirred under same condition for 1 hour. The reaction mixture was concentrated in vacuo and the residue was purified by column chromatography on silica gel, eluting with *n-*hexane only and then a mixture of *n-*hexane and ethyl acetate (20:1) to afford 5.8069 g (82%) of title compound as a colorless oil. R*_f_* = 0.20 (*n-*Hex); ^1^H NMR (400 MHz, DMSO-*d_6_*) *δ* 7.24 (t, 1H, *J* = 7.98), 6.99 (d, 1H, *J* = 8.16), 6.89 (t, 1H, *J* = 7.36), 3.79 (s, 3H), 3.64 (t, 2H, *J* = 7.48), 3.09 (t, 2H, *J* = 7.44)

### Preparation of 1-(2-bromoethyl)-3-methoxybenzene (5b)

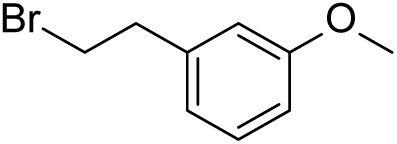

To a solution of 2-(3-methoxyphenyl)ethan-1-ol (1g, 6.57 mmol) in anhydrous dichloromethane (20 mL) was added carbon tetrabromide (3.05 g, 9.20 mmol) and stirred at 0° under nitrogen atmosphere. Triphenylphospine (2.06 g, 7.88 mmol) was added potionwise and stirred under same condition for 1 hour. The reaction mixture was concentrated in vacuo and the residue was purified by column chromatography on silica gel, eluting with *n-*hexane only and then a mixture of *n-*hexane and ethyl acetate (20:1) to afford 1.37 g (97%) of title compound as a colorless oil. R*_f_* = 0.20 (*n-*Hex); ^1^H NMR (400 MHz, DMSO-*d_6_*) *δ* 7.23 (t, 1H, *J* = 7.60), 6.88 (m, 3H), 3.75 (s, 3H), 3.74 (t, 2H, *J* = 7.2), 3.11 (t, 2H, *J* = 7.2)

### General procedure for alkylated compounds (7a-7k) (Method A)

To a solution of aniline (3, 4a-4d) (1.0-3.0 equiv) in acetonitrile, were added potassium carbonate/cesium carbonate (1.0-1.2 equiv), potassium iodide (0.1 equiv), and phenethyl bromide (6a-6g) (1.0 equiv) in a sealed tube. The reaction mixture was heated at 110°C for 36 hours. The reaction mixture was cooled and diluted with ethyl acetate and washed with brine, dried with anhydrous Na_2_SO_4_, and concentrated in vacuo. Purification by column chromatography afforded the desired compound compound.

### General procedure for final KDS compounds (8a-8h) (Method B)

To a solution of amine containing compounds (7e-7k) (1.0 equiv) was dissolved in anhydrous dichloromethane was added 4.0M hydrochloric acid in dioxane (4.0-6.0 equiv). The reaction mixture was stirred at room temperature for 48 hours. The precipitate was filtered and the filtercake was obtained to afford the desired compound

### *tert*-butyl methyl(4-((4-(trifluoromethyl)phenethyl)amino)phenyl)carbamate (7a)

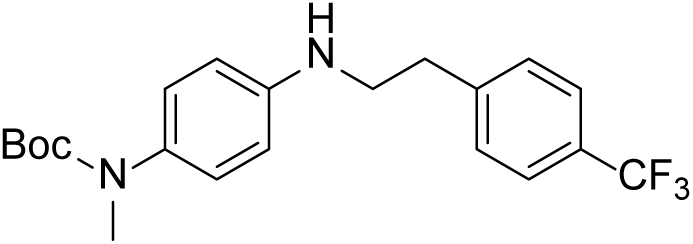

Using method A, *tert*-butyl (4-aminophenyl)(methyl)carbamate (440 mg, 1.98 mmol), 1-(2-bromoethyl)-4-(trifluoromethyl)benzene (500 mg, 1.98 mmol), potassium carbonate (821 mg, 5.94 mmol), and potassium iodide (33 mg, 0.20 mmol) in acetonitrile (10 mL) to give 7a as an off white solid (319 mg, 41%); R*_f_* = 0.45 (*n-*Hex 3: EtOAC 1); ^1^H NMR (400 MHz, DMSO-*d_6_*) *δ* 7.68 (d, 2H, *J* = 8.00 Hz) 7.53 (d, 2H, *J* = 8.00 Hz), 6.96 (d, 2H, *J* = 8.80 Hz), 6.57 (d, 2H, *J* = 8.80 Hz), 5.70 (t, 1H, *J* = 5.60 Hz), 3.28 (t, 2H, *J* = 7.20 Hz), 3.08 (s, 3H), 2.94 (d, 2H, *J* = 7.20 Hz)

### *N*-(4-(trifluoromethyl)phenethyl)aniline (7b)

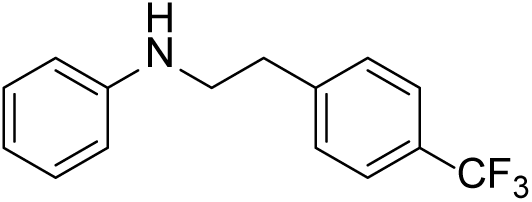

Using method A, aniline (552mg, 5.93 mmol), 1-(2-bromoethyl)-4-(trifluoromethyl)benzene (500mg, 1.98 mmol), potassium carbonate (273 mg, 1.98 mmol), and potassium iodide (33 mg, 0.20 mmol) in acetonitrile (10 mL) gave 7b as a white solid (50.4 mg, 8%); R*_f_* = 0.50 (*n-*Hex 9: EtOAC 1); HPLC purity: 7.7 min, 99.2%; mp 20-30 °C; ^1^H NMR (400 MHz, DMSO-*d_6_*) *δ* 7.65 (d, 2H, *J* = 8.12 Hz), 7.50 (d, 2H, *J* = 8.04 Hz), 7.07 (t, 2H, *J* = 8.16 Hz), 6.59 (d, 2H, *J* = 8.12 Hz), 6.53 (t, 1H, *J* = 7.24 Hz), 5.65 (t, 1H, *J* = 5.56 Hz), 3.30-3.25 (m, 2H), 2.92 (t, 2H, *J* = 7.28 Hz); ^13^C NMR (100 MHz, DMSO-*d_6_*) *δ* 149.1, 145.5, 130.0, 129.4, 127.7, 127.4, 127.1, 126.8, 126.3, 125.5 (q, *J_C-F_* = 3.57 Hz), 116.2, 112.5, 44.5 (**C**H_2_), 35.1 (**C**H_2_)

### 4-((4-(trifluoromethyl)phenethyl)amino)phenol (7c)

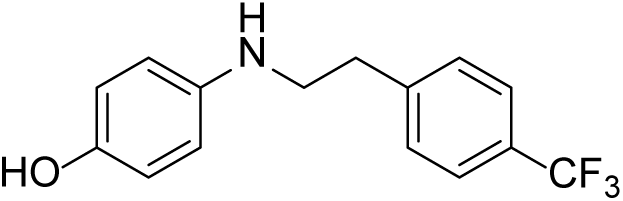

Using method A, 4-aminophenol (258.6 mg, 2.37 mmol), 1-(2-bromoethyl)-4-(trifluoromethyl)benzene (200 mg, 0.79 mmol), potassium carbonate (200 mg, 0.79 mmol), and potassium iodide (13 mg, 0.08 mmol) in acetonitrile (4 mL) gave 7c as a white solid (111 mg, 53%); R*_f_* = 0.30 (*n-*Hex 3: EtOAC 1); HPLC purity: 9.3 min, >99.9%; mp 110-120 °C; ^1^H NMR (400 MHz, DMSO-*d_6_*) *δ* 8.39 (s, 1H), 7.64 (d, 2H, *J* = 8.00 Hz), 7.49 (d, 2H, *J* = 8.00 Hz), 6.55 (d, 2H, *J* = 8.56 Hz), 6.45 (d, 2H, *J* = 8.64 Hz), 4.99 (s, 1H), 3.21-3.17 (m, 2H), 2.89 (t, 2H, J = 7.20 Hz) ^13^C NMR (100 MHz, DMSO-*d_6_*) *δ* 148.8, 145.7, 134.6, 130.0, 125.5 (q, *J_C-F_* = 4.06), 116.2, 113.9, 103.1 45.7 (**C**H_2_), 35.3 (**C**H_2_)

### 3-((4-(trifluoromethyl)phenethyl)amino)benzoic acid (7d)

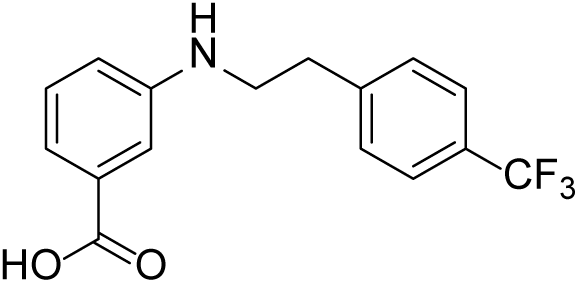

Using method A, 3-aminobenzoic acid (813 mg, 5.93 mmol), 1-(2-bromoethyl)-4-(trifluoromethyl)benzene (500 mg, 1.98 mmol), potassium carbonate (273 mg, 1.98 mmol), and potassium iodide (33 mg, 0.20 mmol) in acetonitrile (10 mL) gave 7d as a white solid (49 mg, 8%); R*_f_* = 0.30 (*n-*Hex 3: EtOAC 1); HPLC purity: 12.6 min, 98.4%; mp 82-86 °C; ^1^H NMR (400 MHz, DMSO-*d_6_*) *δ* 12.65, (s, 1H), 7.66 (d, 2H, *J* = 8.08 Hz), 7.51 (d, 2H, *J* = 7.92 Hz), 7.21-7.18 (m, 2H), 7.14 (d, 1H, *J* = 7.40 Hz), 6.83 (d, 1H, *J* = 7.08 Hz), 5.98 (s, 1H), 3.32 (t, 2H, *J* = 7.04 Hz), 2.94 (t, 2H, *J* = 6.88 Hz) ^13^C NMR (100 MHz, DMSO-*d_6_*) *δ* 166.6 (**C**(O)), 149.5, 143.8, 130.7, 130.2, 129.5, 127.6 (q, *J_C-F_* = 31.56 Hz), 125.7 (q, *J_C-F_* = 3.71 Hz), 124.9 (q, *J_C-F_* = 270.27 Hz), 118.9, 116.7, 114.5, 64.9(**C**H_2_), 34.6(**C**H_2_)

### *N*^1^*,N*^1^-dimethyl-*N*^4^-(4-(trifluoromethyl)phenethyl)benzene-1,4-diamine (7e)

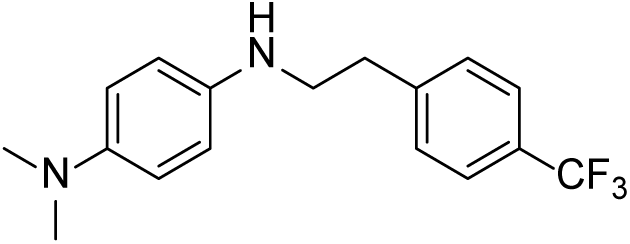

Using method A, *N^1^,N^1^*-dimethylbenzene-1,4-diamine (646 mg, 4.74 mmol), 1-(2-bromoethyl)-4-(trifluoromethyl)benzene (1.0g, 3.96 mmol), potassium carbonate (1.64 g, 11.88 mmol), and potassium iodide (66 mg, 0.40 mmol) in acetonitrile (20 mL) gave 7e as a dark brown oil (300 mg, 25%); R*_f_* = 0.37 (*n-*Hex 3: EtOAC 1) ^1^H NMR (400 MHz, DMSO-*d_6_*) *δ* 7.66 (d, 2H, *J* = 8.00 Hz), 7.50 (d, 2H, *J* = 8.00 Hz), 6.65 (d, 2H, *J* = 8.96 Hz), 6.54 (d, 2H, *J* = 8.96 Hz), 5.02 (t, 1H, *J* = 6.00 Hz), 3.22 (q, 2H, *J* = 6.44 Hz), 2.91 (t, 2H, *J* = 7.12 Hz), 2.72 (s, 6H)

### *tert*-butyl (4-((2-methoxyphenethyl)amino)phenyl)(methyl)carbamate (7f)

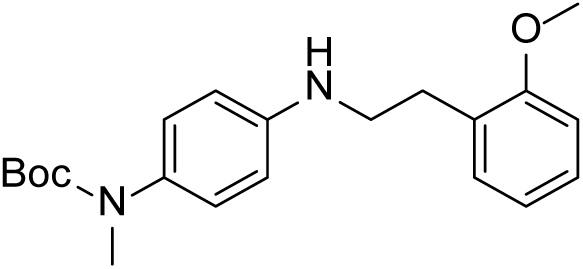

Using method A, I-butyl (4-aminophenyl)(methyl)carbamate (2.10 g, 9.45 mmol), 1-(2-bromoethyl)-2-methoxybenzene (2.03 g, 9.45 mmol), cesium carbonate (3.69 g, 11.34 mmol), and potassium iodide (157.7 mg, 0.95 mmol) in acetonitrile (20 mL) gave 7f as a yellow oil (1.21 g, 36%); R*_f_* = 0.42 (*n-*Hex 3: EtOAC 1); ^1^H NMR (400 MHz, DMSO-*d_6_*) *δ* 7.23-7.18 (m, 2H), 6.97 (d, 1H, *J* = 7.92 Hz), 6.93 (d, 2H, *J* = 8.64 Hz), 6.88 (t, 1H, *J* = 7.36 Hz), 6.54 (d, 2H, *J* = 8.76 Hz), 5.69 (t, 1H, *J* = 5.68 Hz), 3.81 (s, 3H), 3.18-3.13 (m, 2H), 3.07 (s, 3H), 2.80 (t, 2H, *J* = 6.84 Hz), 1.35 (s, 9H)

### *tert*-butyl (4-((3-methoxyphenethyl)amino)phenyl)(methyl)carbamate (7g)

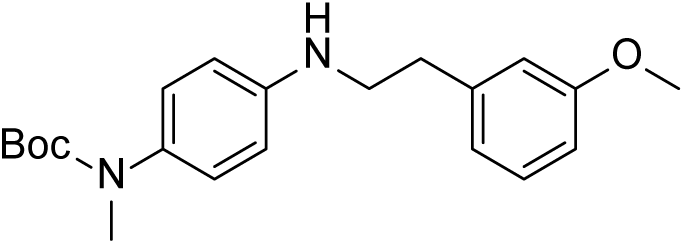

Using method A, *tert*-butyl (4-aminophenyl)(methyl)carbamate (455 mg, 2.05 mmol), 1-(2-bromoethyl)-3-methoxybenzene (400 mg, 1.86 mmol), potassium carbonate (771 mg, 5.58 mmol), and potassium iodide (32 mg, 0.19 mmol) in acetonitrile (5 mL) gave 7g as a yellow oil (244 mg, 37%); R*_f_* = 0.41 (*n-*Hex 3: EtOAC 1); ^1^H NMR (400 MHz, DMSO-*d_6_*) *δ* 7.22 (t, 1H, *J* = 8.04 Hz), 6.95 (d, 2H, *J* = 8.60 Hz), 6.86-6.85 (m, 2H), 6.80-6.77 (m, 1H), 6.55 (d, 2H, *J* = 8.72 Hz), 5.65 (t, 1H, *J* = 5.64 Hz), 3.72 (s, 3H), 3.23 (q, 2H, *J* = 6.80 Hz), 3.08 (s, 3H), 2.81 (t, 2H, *J* = 7.60 Hz), 1.36 (s, 9H)

### *tert*-butyl (4-((4-methoxyphenethyl)amino)phenyl)(methyl)carbamate (7h)

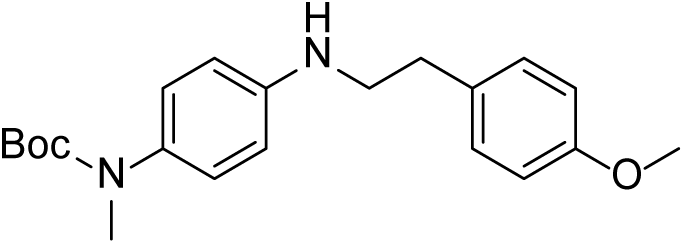

Using method A, *tert*-butyl (4-aminophenyl)(methyl)carbamate (340 mg, 1.53 mmol), 1-(2-bromoethyl)-4-methoxybenzene (300 mg, 1.39 mmol), potassium carbonate (578 mg, 4.18 mmol), and potassium iodide (23 mg, 0.14 mmol) in acetonitrile (5 mL) gave 7h as a yellow oil (244 mg, 37%); R*_f_* = 0.41 (*n-*Hex 3: EtOAC 1); ^1^H NMR (400 MHz, DMSO-*d_6_*) *δ* 7.20 (d, 2H, *J* = 8.52 Hz), 6.94 (d, 2H, *J* = 8.60 Hz), 6.87 (d, 2H, *J* = 8.52 Hz), 6.54 (d, 2H, *J* = 8.72 Hz), 5.63 (t, 1H, *J* = 5.60 Hz), 3.73 (s, 3H), 3.21-3.16 (m, 2H), 3.08 (s, 3H), 2.77 (t, 2H, *J* = 7.24), 1.36 (s, 9H)

### *tert*-butyl methyl(4-(phenethylamino)phenyl)carbamate (7i)

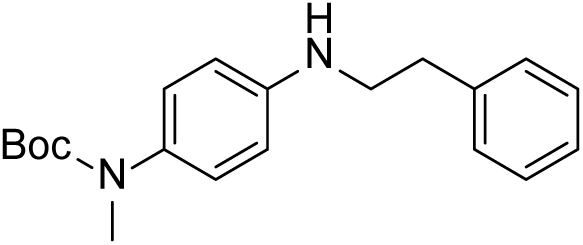

Using method A, *tert*-butyl (4-aminophenyl)(methyl)carbamate (500 mg, 2.25 mmol), (2-bromoethyl)benzene (416 mg, 2.25 mmol), cesium carbonate (879.4 mg, 2.70 mmol), and potassium iodide (37 mg, 0.23 mmol) in acetonitrile (10 mL) gave 7i as a yellow oil (160.4 mg, 20%); R*_f_* = 0.32 (*n-*Hex 9: EtOAC 1); ^1^H NMR (400 MHz, DMSO-*d_6_*) *δ* 7.32-7.27 (m, 4H), 7.22-7.18 (m, 1H), 6.93 (d, 2H, *J* = 8.68 Hz), 6.54 (d, 2H, *J* = 8.80), 5.65 (t, 1H, *J* = 5.72 Hz), 3.25-3.19 (m, 2H), 3.07 (s, 3H), 2.83 (t, 2H, *J* = 7.76), 1.35 (s, 9H)

### *tert*-butyl methyl(4-((3-(trifluoromethyl)phenethyl)amino)phenyl)carbamate (7j)

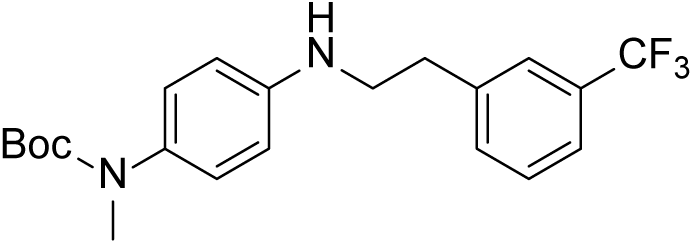

Using method A, *tert*-butyl (4-aminophenyl)(methyl)carbamate (440 mg, 1.98 mmol), 1-(2-bromoethyl)-3-(trifluoromethyl)benzene (500 mg, 1.98 mmol), potassium carbonate (821 mg, 5.94 mmol), and potassium iodide (33 mg, 0.20 mmol) in acetonitrile (7 mL) to give 7j as an off white solid (395 mg, 51%); R*_f_* = 0.44 (*n-*Hex 3: EtOAc 1); ^1^H NMR (400 MHz, DMSO-*d_6_*) *δ* 7.64 (s, 1H), 7.61-7.52 (m, 3H), 6.95 (d, 2H, *J* = 8.72 Hz), 6.56 (d, 2H, *J* = 8.80 Hz), 5.69 (t, 1H, 5.68 Hz), 3.30-3.25 (m, 2H), 3.08 (s, 3H), 2.94 (t, 2H, *J* = 7.16 Hz)

### *tert*-butyl methyl(4-((2-(trifluoromethyl)phenethyl)amino)phenyl)carbamate (7k)

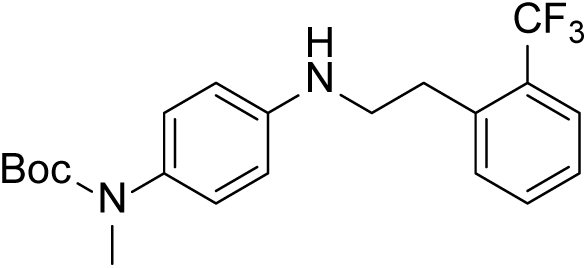

Using method A, *tert*-butyl (4-aminophenyl)(methyl)carbamate (295 mg, 1.33 mmol), 1-(2-bromoethyl)-2-(trifluoromethyl)benzene (337 mg, 1.33 mmol), potassium carbonate (552 mg, 3.99 mmol), and potassium iodide (22 mg, 0.13 mmol) in acetonitrile (7 mL) to give 7j as an off white solid (319 mg, 41%); Rf = 0.45 (*n-*Hex 3: EtOAc 1); ^1^H NMR (400 MHz, DMSO-*d_6_*) *δ* 7.73 (d, 1H, *J* = 7.72 Hz), 7.65 (t, 1H, *J* = 7.36 Hz), 7.58 (d, 1H, *J* = 7.60 Hz), 7.46 (t, 1H, *J* = 7.48 Hz), 6.97 (d, 2H, *J* = 8.64 Hz), 6.58 (d, 2H, *J* = 8.76 Hz), 5.85 (t, 1H, *J* = 5.84 Hz), 3.29-3.24 (m, 2H), 3.09 (s, 3H), 3.01 (t, 2H, *J* = 7.96 Hz), 1.37 (s, 9H)

### *N*^1^-methyl-*N*^4^-(4-(trifluoromethyl)phenethyl)benzene-1,4-diamine hydrochloride salt (8a, KDS12017)

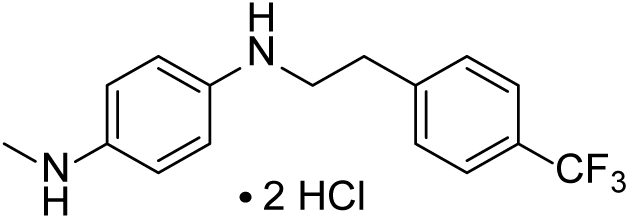

Using method B, *tert*-butyl methyl(4-((4-(trifluoromethyl)phenethyl)amino)phenyl)carbamate (316 mg, 0.80 mmol), 4.0M hydrochloric acid in dioxane (1.20 mL, 4.80 mmol) in dichloromethane (5 mL) to give 8a as an off white solid (270 mg, 92%); HPLC purity: 8.1 min, >99.9%; mp 246-250 °C; ^1^H NMR (400 MHz, DMSO-*d_6_*) *δ* 10.58 (br, 1H), 7.66 (d, 2H, *J* = 7.96 Hz), 7.50 (d, 2H, *J* = 7.92 Hz), 7.13 (s, 2H), 6.82 (s, 2H), 3.34 (t, 2H, *J* = 7.04 Hz), 2.96 (t, 2H, *J* = 7.40 Hz), 2.79 (s, 3H); ^13^C NMR (100 MHz, DMSO-*d_6_*) *δ* 144.51, 130.07, 129.81, 127.50 (q, *J_C-F_* = 31.63 Hz), 126.24, 125.62 (q, *J_C-F_* = 3.81 Hz), 123.54, 46.40 (**C**H_2_), 35.74 (**C**H_2_), 33.92 (**C**H_3_)

### *N*^1^*,N*^1^-dimethyl-*N*^4^-(4-(trifluoromethyl)phenethyl)benzene-1,4-diamine hydrochloride salt (8b, KDS12008)

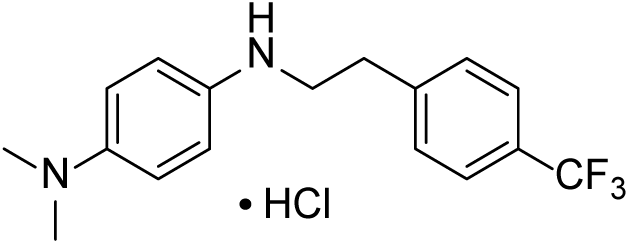

Using method B, *N*^1^,*N*^1^-dimethyl-*N*^4^-(4-(trifluoromethyl)phenethyl)benzene-1,4-diamine (300 mg, 0.97 mmol), 4.0M hydrochloric acid in dioxane (0.73 mL) in dichloromethane (7 mL) to give 8b as an off white solid; HPLC purity: 8.0 min, 99.7%; mp 216-241 °C ^1^H NMR (400 MHz, DMSO-*d_6_*) *δ* 7.89 (b, 2H) 7.66 (d, 2H, *J* = 8.00), 7.51 (d, 2H, *J* = 7.96 Hz), 7.40 (s, 2H), 6.89 (s, 2H), 3.36 (t, 2H, *J* = 7.28Hz), 3.01 (s, 6H), 2.98 (t, 3H, *J* = 7.04 Hz); ^13^C NMR (100 MHz, DMSO-*d_6_*) *δ* 144.47, 130.07, 128.94, 127.50 (q, *J_C-F_* = 31.48 Hz), 126.24, 125.62 (q, *J_C-F_* = 3.45 Hz), 123.54, 46.21 (**C**H_2_), 45.19 (**C**H_3_), 33.90 (**C**H_2_)

### *N*^1^-(2-methoxyphenethyl)-*N*^4^-methylbenzene-1,4-diamine hydrochloride salt (8c, KDS12025)

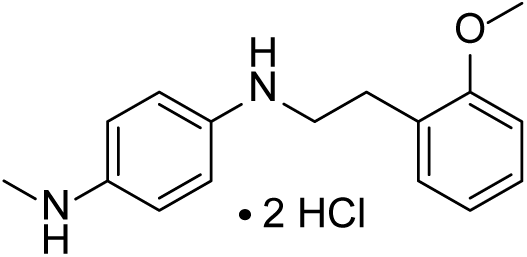

Using method B, *tert*-butyl (4-((2-methoxyphenethyl)amino)phenyl)(methyl)carbamate (586 mg, 1.64 mmol), 4.0M hydrochloric acid in dioxane (1.65 mL, 6.58 mmol) in dichloromethane (10 mL) to give 8c as a white solid (436 mg, 81%); HPLC purity: 6.8 min, 99.7%; mp 210-212 °C; ^1^H NMR (400 MHz, DMSO-*d_6_*) *δ* 7.25-7.17 (m, 3H), 7.09 (s, 2H), 6.98 (d, 1H, *J* = 8.04), 6.89 (t, 3H, *J* = 7.36 Hz), 3.80 (s, 3H), 3.24 (t, 2H, *J* = 7.12 Hz), 2.84 (t, 2H, *J* = 8.16 Hz), 2.79 (s, 3H); ^13^C NMR (100 MHz, DMSO-*d_6_*) *δ* 157.60, 130.42, 128.49, 126.16, 124.94, 124.79, 122.13, 120.86, 119.99, 111.19, 55.80(**C**H_3_), 47.45(**C**H_2_), 35.50(**C**H_2_), 27.96(**C**H_3_)

### *N*^1^-(3-methoxyphenethyl)-*N*^4^-methylbenzene-1,4-diamine hydrochloride salt (8d)

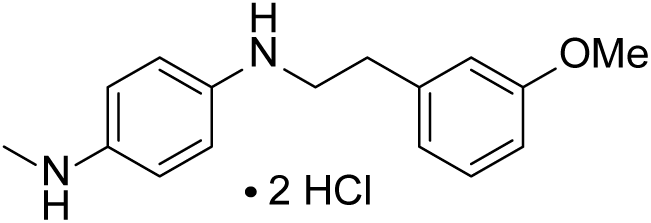

Using method B, *tert*-butyl (4-((3-methoxyphenethyl)amino)phenyl)(methyl)carbamate (240 mg, 0.67 mmol), 4.0M hydrochloric acid in dioxane (1.01 mL, 4.02 mmol) in dichloromethane (10 mL) to give 8d as a white solid (193 mg, 87%); HPLC purity: 6.7 min, >99.9%; mp 245-249 °C; ^1^H NMR (400 MHz, DMSO-*d_6_*) *δ* 9.18 (b, 4H), 7.29 (d, 2H, *J* = 7.29 Hz), 7.22 (t, 1H, *J* = 7.92 Hz), 7.16 (d, 2H, *J* = 7.84 Hz), 6.84-6.78 (m, 3H), 3.74 (s, 3H), 3.37 (t, 2H, *J* = 8.24 Hz), 2.92 (t, 2H, *J* = 7.44 Hz), 2.80 (s, 3H); ^13^C NMR (100 MHz, DMSO-*d_6_*) *δ* 159.82, 140.34, 129.92, 124.58, 124.46, 121.48, 121.33, 118.82, 114.75, 112.41, 55.44 (**C**H_3_), 48.30 (**C**H_2_), 35.34 (**C**H_2_), 33.46 (**C**H_3_)

### *N*^1^-(4-methoxyphenethyl)-*N*^4^-methylbenzene-1,4-diamine (8e)

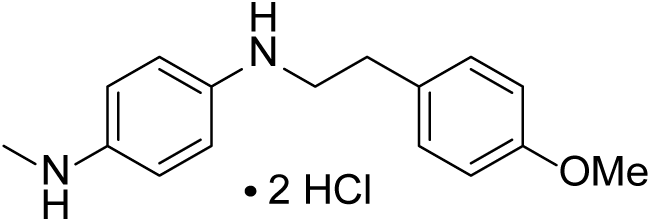

Using method B, *tert*-butyl (4-((4-methoxyphenethyl)amino)phenyl)(methyl)carbamate (586 mg, 1.64 mmol), 4.0M hydrochloric acid in dioxane (1.65 mL, 6.58 mmol) in dichloromethane (10 mL) to give 8e as a white solid (436 mg, 81%); HPLC purity: 6.6 min, 99.4%; mp 220-245 °C; ^1^H NMR (400 MHz, DMSO-*d_6_*) *δ* 10.10 (b, 4H), 7.34 (d, 2H, *J* = 8.56 Hz), 7.25 (d, 2H, *J* = 8.12 Hz), 7.18 (d, 2H, *J* = 8.48 Hz), 6.87 (d, 2H, *J* = 8.48 Hz), 3.72 (s, 3H), 3.34 (t, 2H, *J* = 8.28 Hz), 2.90 (t, 2H, *J* = 7.48 Hz), 2.81 (s, 3H); ^13^C NMR (100 MHz, DMSO-*d_6_*) *δ* 158.38, 13036, 130.12, 124.77, 124.64, 121.72, 119.86, 114.36, 55.50 (**C**H_3_), 49.24(**C**H2), 35.33(**C**H_2_), 32.28 (**C**H_3_)

### *N*^1^-methyl-*N*^4^-phenethylbenzene-1,4-diamine hydrochloride salt (8f)

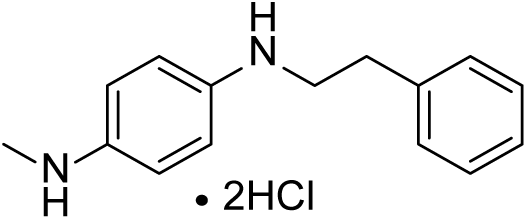

Using method B, *tert*-butyl methyl(4-(phenethylamino)phenyl)carbamate (160 mg, 0.49 mmol), 4.0M hydrochloric acid in dioxane (0.73 mL) in dichloromethane (5mL) to give 8f as an off white solid (111 mg, 76%); HPLC purity: 7.0 min, >99.9%; mp 246-247 °C; H NMR (400 MHz, DMSO-*d_6_*) *δ* 10.32 (b, 1H), 7.33-7.26 (m, 4H), 7.24-7.20 (m, 1H), 7.08 (s, 2H), 6.88 (s, 2H), 3.31 (t, 2H, *J* = 7.40 Hz), 2.87 (t, 2H, *J* = 7.92 Hz), 2.78 (s, 3H)); ^13^C NMR (100 MHz, DMSO-*d_6_*) *δ* 139.1, 129.1, 128.9, 126.8, 121.0, 117.6, 47.7 (**C**H_2_), 35.2 (**C**H_2_), 33.8 (**C**H_3_)

### *N*^1^-methyl-*N*^4^-(3-(trifluoromethyl)phenethyl)benzene-1,4-diamine hydrochloride salt (8g)

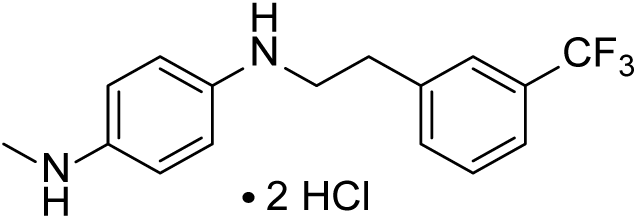

Using method B, *tert*-butyl methyl(4-((3-(trifluoromethyl)phenethyl)amino)phenyl)carbamate (395 mg, 0.34 mmol), 4.0M hydrochloric acid in dioxane (1.5 mL, 6.00 mmol) in dichloromethane (7 mL) to give 8g as an off white solid (351 mg, 96%); mp 210-246 °CHPLC purity: 7.8 min, >99.9%; mp 225-244 °C ^1^H NMR (400 MHz, MeOD-*d_4_*) *δ* 7.59-7.7.50 (m, 4H), 7.43 (d, 2H, *J*= 8.92 Hz), 7.30 (d, 2H, *J* = 6.84 Hz), 3.62-3.58 (m, 2H), 3.12 (t, 2H, *J* = 8.00 Hz), 3.30 (s, 3H); ^13^C NMR (100 MHz, DMSO-*d_6_*) *δ* 140.66, 133.55, 129.84, 125.84, 123.60, 122.06, 117.98, 47.55 (**C**H_2_), 35.80 (**C**H_2_), 33.43 (**C**H_3_)

### *N*^1^-methyl-*N*^4^-(2-(trifluoromethyl)phenethyl)benzene-1,4-diamine hydrochloride salt (8h)

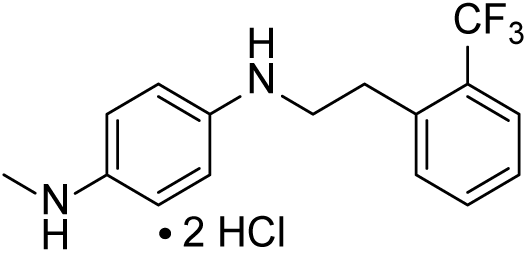

Using method B, *tert*-butyl methyl(4-((2-(trifluoromethyl)phenethyl)amino)phenyl)carbamate (80 mg, 0.20 mmol), 4.0M hydrochloric acid in dioxane (0.30 mL, 1.22 mmol) in dichloromethane (5 mL) to give 8h as an off white solid (53.1 mg, 72%); HPLC purity: 7.7 min, 99.2%; mp 221-246 °C; ^1^H NMR (400 MHz, DMSO-*d_6_*) *δ* 9.93 (b, 4H), 7.71 (d, 1H, *J* = 7.88 Hz), 7.65 (t, 1H, *J* = 7.48 Hz), 7.58 (d, 1H, *J* = 7.56 Hz), 7.46 (t, 1H, *J* = 7.48 Hz), 7.35 (d, 2H, *J* = 8.44 Hz), 7.08 (d, 2H, *J* = 7.96 Hz), 3.36 (t, 2H, *J* = 7.24 Hz), 3.11 (t, 2H, *J* = 7.20 Hz), 2.82 (s, 3H); ^13^C NMR (100 MHz, DMSO-*d_6_*) *δ* 137.36, 133.13, 132.21, 127.69 (q, *J_C-F_* = 32.55 Hz), 127.55, 126.27 (q, *J_C-F_* = 5.63 Hz), 124.91 (q, *J_C-F_* = 255.06 Hz), 122.50, 117.23, 47.13(**C**H_2_), 35.94(**C**H_2_), 30.65(**C**H_3_)

### Docking and Binding Energy calculations

Molecular modeling was performed using BIOVIA Discovery Studio software^42^ under pH 7.4 and standard protocol settings if not specified otherwise. HRP and Hb protein structures were obtained from the protein databank (PDB ID 7ATJ^43^, 2DN1^44^, respectively). The heme iron Fe(IV)O state was set up in both systems, mimicking the HRP Compound II and ferryl Hb catalytic stages. In the HRP catalytic pocket, crucial crystal water molecule HOH1054^45^ was retained. Conversely, in the Hb catalytic pocket, a hydrogen peroxide molecule was placed 2.6 Å under the porphyrin ring pyrrole and within 2.2 Å of both the oxo-iron and distal histidine (H58 and H63 in ɑ and β chains, respectively). Small molecule docking was performed using the CDOCKER protocol. In the HRP protein system, compounds KDS12025 and HTPEB were docked into the catalytic binding site, and the robustness of the docking protocol was verified through the successful re-docking of ferulic acid within 1.5 Å of the co-crystallized ligand pose. For Hb, multiple binding sites in the Hbα and Hbβ subunits of the Hb tetramer were explored, guided by prior published Hb-small molecule docking studies^46–50^.

The ligand binding energy (ΔG_bind_) for the top 5 docking poses was determined using the MM-GBSA method under the Generalized Born with Molecular Volume (GBMV) solvent model and additionally, *in situ* ligand minimization was included in the protocol, where all protein residues within 6 Å of the ligand were treated as flexible.

### Binding affinity assay

All ITC experiments were conducted using a MicroCal Auto-iTC200 (Malvern Panalytical) at the Korea Basic Science Institute (Ochang, Korea). Binding affinity was measured using 40 µl of 1 mM KDS12025 and 200 µl of 0.1 mM purified human Hb in sodium phosphate buffer (pH 7). For each titration experiment, 2 µl of KDS12025 was injected into purified human Hb for 4 s at intervals of 150 s at 25 °C. Overall, 19 injections were performed for each experiment, and the data were analyzed using MicroCal Origin 7.0 software.

### Bubbling liberation

CAT, HRP, and Hb were reacted with H_2_O_2_ to assess oxygen production through bubbling liberation. The reactions were observed under a microscope, and bubbles in a 1 x 1 mm^2^ area were quantified using ImageJ (NIH) software. The effect of DMSO, HTPEB, and KDS12025 on bubbling was evaluated. To quantitatively measure the bubbling, reactions were conducted in sealed tubes connected to an inverted cylinder submerged in a water bath to record the time and volume of the bubbles produced.

To visualize bubble formation, CAT and Hb were prepared at 10 mg/ml and 0.08 mg/ml concentrations. H_2_O_2_ was added at 10 M and 0.1 M. Vehicle, HTPEB, and KDS12025 were added to assess their effects. A similar procedure was followed for Hb. Bubble formation was visually monitored to evaluate catalytic H_2_O_2_ breakdown and oxygen release.

### Off-target selectivity assay

The possibility of KDS12025 on kinase off-target, the KINOMEscan panel of scanEDGE assay was conducted in DiscoverX (CA, USA) using a site-directed competition binding with the test compounds. The assay results are given in Supplementary Table 4 and experimental information is available on the DiscoverX web page (www.discoverx.com). Moreover, the Delta SafetyScreen87 panel assay was conducted in Eurofins Discovery (Cerep, France). The assay results are given in Supplementary Table 5, and experimental information is available on the DiscoverX web page (https://www.eurofinsdiscoveryservices.com).

### BBB-permeability test

A parallel artificial membrane permeability assay (PAMPA)^51^ was performed to validate the blood-brain barrier permeability of KDS12025. The procedure began by preparing a donor solution of the compound in a pH 7.4 donor buffer. This solution was added to the wells of a deep well plate, followed by a high-sensitivity UV plate filled with donor buffer to serve as the blank sample. An initial sample for the UV plate was taken to establish a baseline reading. The PAMPA sandwich was then assembled by adding the donor plate mixture to the donor plate well, placing a BBB-lipid coated membrane on top, and filling an acceptor plate with acceptor buffer. This setup was incubated at 25°C for a specific duration to allow diffusion. Post-incubation, a sample from the acceptor plate was taken, and the UV absorbance was measured to determine the permeated amount of the test compound. The permeability was then calculated using the PAMPA explorer program, considering the initial donor concentration and the acceptor’s concentration after incubation.

### Cultured primary astrocytes

Primary hippocampal astrocytes were obtained from postnatal day 0-2 (P0-2) C57BL/6 mice as previously described ^52^. The hippocampal tissue was dissected free of meninges, minced, and dissociated into a single-cell suspension by trituration. Cells were grown in Dulbecco’s modified Eagle’s medium supplemented with D-glucose (4500 mg/L), L-glutamine, sodium pyruvate (110 mg/L), 10% heat-inactivated horse serum, 10% heat-inactivated fetal bovine serum, and 1000 U/ml penicillin-streptomycin. Cells were maintained at 37 °C in a humidified atmosphere containing 5% CO_2_.

### Oligomerized Aβ_42_ aggregates

Amyloid-beta (Aβ)_42_ monomers (Abcam; DAEFRHDSGYEVHHQKLVFFAEDVGSNK GAIIGLMVGGVVIA) were dissolved to 1 mM in 1% ammonium hydroxide. Aβ_42_ monomers were diluted to 100 µM in Phosphate Buffered Saline (PBS). The solution was incubated at a four °C rocker for 24 hours and stored at −80°C for further use as an aggregated form of Aβ_42_ as previously described^53,54^.

### Intracellular H_2_O_2_ detection

Intracellular H_2_O_2_ levels of astrocytes were determined using 2′,7′-dichlorodihydrofluorescein diacetate (DCFDA, Thermofisher). Primary cultured astrocytes were seeded onto 96-well black plates (ibid, USA) and incubated with oligomerized Aβ_42_ in the presence of 10 µM KDS12008, 17, 25, HTPEB, and sodium pyruvate (10 mM). Then, these cells were washed with Ca^2+^ and Mg^2+^ contained PBS, followed by the application of 30 µM DCFDA and incubated at 37°C for 30 minutes. The mean fluorescence intensity was measured using iD5 microplate readers at ex/em=485/530 nm.

Dr. Andre Berndt kindly provided H2O2-specific, high sensitivity, and fast on-and-off kinetics, genetically encoded oROS-G plasmid. The oROS-G was cloned under the GFAP104 promoter and delivered into primary cultured hippocampal astrocytes using the NeoN^TM^ transfection system (Invitrogen). Astrocytes were plated in 96-well black plates (ibid) and fluorescence changes were measured using an iD5 microplate reader for endpoint analysis or imaged over 40 hours using an A1 Nikon confocal microscope for live cell imaging.

### Mouse

All male and female mice were grouped-housed in a temperature- and humidity-controlled environment with a 12-hour light/dark cycle. Handling and animal care were performed according to the Institutional Animal Care and Use Committee of the Institute for Basic Science (IBS-2023-006, Daejeon, Korea). For an animal model of AD, both sexes of 10 to 18-month-old APP/PS1 mice were maintained and used as previously described ^55,56^. For testing the novel H_2_O_2_-decomposing enhancers effect in the AD model, mice were treated by i.p. injection with KDS12008 (3 mg/kg/day), KDS12017 (3 mg/kg/day), and KDS12025 (3 and 10 mg/kg/day) once per day for one or two weeks depends on experimental protocol. All drugs were dissolved in 200 μl of saline, and the amount of each drug was calculated based on the weight (kg) of the mouse. Moreover, oral administration of KDS12025 (3 mg/kg/day) in the AD model for two weeks *ad libitum*. Both sexes of 8 to 10-week-old C57BL/6J mice were used for PhenoMaster (TSD systems, Germany). GFAP-CreERT2/iDTR (GiD) lines were maintained by crossing iDTR transgenic mice with GFAP-CreERT2. All mice were randomly distributed into different groups with matched ages.

### Passive avoidance test

Sufficiently handled mice were ready before the PAT. Mice were placed in the two-compartment (light and dark) shuttle chamber with a shock generator (Ugo Basile, Italy). On the first day, mice were placed in the light compartment for the acquisition. Mice were allowed to explore the light compartment for 60 seconds, after which the door separating the light and dark compartments was raised to allow the mice to enter the dark compartment. When the mice entered a dark compartment, the separating door was closed immediately, and an electrical foot shock (0.5 mA, 2 s duration) was delivered through the floor grid. Then, the mice were moved back to the home cage, and the retention trial was conducted 24 hours after the acquisition. On the second day, the mice were again placed in the light compartment for the retention trial. After 60 seconds, the door was raised to allow the mice to enter the dark compartment. The latency to step through the dark compartment before and after the electric shock was recorded for up to 540 seconds.

### Novel place recognition

Before the novel place recognition (NPR) test, mice underwent sufficient handling and were concurrently habituated to an open field test (OFT) in a square chamber (40 cm x 40 cm x 40 cm). After the habituation, mice were placed in the open field with two identical objects positioned in the first and second quadrants of the cage. Mice were free to explore the objects for 10 min and returned to the home cage for 1 hour. To test spatial recognition memory, one of the two identical objects was placed in a novel place (quadrant), and the mice were re-entered into the open field chamber for 10 min. The discrimination index (DI) was calculated as the percentage of time spent on the novel place over the total time spent on both the novel and the familiar place.

### Open field test

For the open field test (OFT), all the mice were placed in an acrylic container (40 cm x 40 cm x 40 cm) for a 10-minute acclimation period. The field was segmented into central (12 cm x 12 cm) and peripheral zones for analysis. Travel distance, center zone duration, and frequency were quantified using Ethovision XT software (Noldus, Netherlands).

### Respiration and metabolic analysis

For respiration and metabolic analysis, PhenoMaster (TSE systems, Germany) was employed. All C57BL/6J mice aged 8 to 10 weeks old underwent an initial chamber habituation and gas calibration on day 0. Starting on day 3, the mice were administered KDS12025 dissolved in water at dosages of 10, 1, and 0.1 mg/kg/day. Respiratory and metabolic measurements, including oxygen consumption, carbon dioxide production, respiratory exchange ratio (RER, calculated as VCO2/VO2), as well as food and drink intake and energy expenditure, were conducted from day 4. The recording ends on day 8 with data collection.

### Virus and diphtheria toxin injection

Mice were deeply anesthetized with vaporized 1% isoflurane and placed into stereotaxic frames (RWD Life Science, China). Following an incision on the midline of the scalp, a hole was drilled into the skull above the hippocampus. AAV-GFAP104-GFP or AAV-GFAP104-DTR-GFP were bilaterally microinjected into the hippocampus CA1 (AP, −2.0 mm; ML, ± 1.5 mm; DV, −1.65 mm from the bregma). Diphtheria toxin (2 mg/ml) was administered for 16 days as previously described ^56^. For shRNA knockdown, AAV-pSicoR-Hbβ-shRNA-mCherry or AAV-pSicoR-Sc-shRNA-mCherry were microinjected into the same coordinate. A total of 0.8 µl of the virus was injected using a syringe pump (KD Scientific, USA). The procedures of injection were implemented before 5 weeks of the experiments. All viruses used in this study were produced at the Institute for Basic Science Virus Facility (IBS virus facility, Korea). To investigate the 0.1 mg/kg/day effects of KDS12025, GiD mice were injected with diphtheria toxin (2 mg/ml) daily for 16 days. Concurrently, KDS12025 was administered *ad libitum* (0.1 mg/kg/day).

### α-synuclein overexpression mouse model

Stereotaxic injections were performed to deliver the AAV-CMV-A53T virus or PBS (as a control) into the right SN (AP, −3.2 mm; ML, −1.3 mm; DV, −4.0 mm from the bregma) of mice. The injections were administered at a rate of 0.2 μl/min, with a total volume of 1 μl. The procedure was conducted under general anesthesia, induced by 1% isoflurane.

Mice were habituated to the rotarod at speeds of 5, 10, and 15 rpm. The motor coordination and endurance were tested at 20 rpm for a maximum of 3 minutes. The latency to fall was recorded during the test to evaluate motor deficits associated with PD and the therapeutic effects of KDS12025.

### CIA mouse model and histological analysis

CIA was induced in DBA1 mice by intradermal injection of type II collagen emulsified in Complete Freund’s Adjuvant (CFA) on day 0, followed by a booster injection of type II collagen in Incomplete Freund’s Adjuvant (IFA) on day 21. Starting from day 23, mice were administered KDS12025 in drinking *ad libitum* at doses of 0, 0.1, or 1 mg/kg/day until day 42. To assess joint inflammation and cartilage thickness in CIA mice, joint samples were collected and fixed in 10% formalin. The samples were then decalcified, embedded in paraffin, and sectioned. For histological evaluation, sections were stained with H&E to assess inflammation, and with Toluidine Blue to evaluate cartilage thickness. The stained sections were examined under a light microscope, and images were captured for quantitative analysis of inflammatory cell infiltration and cartilage integrity.

### ALS mouse model and survival analysis

SOD1^G93A^ mice were used in the ALS mouse model. The mice were divided into three groups: Control, SOD1, and KDS12025 (1 and 10 mg/kg/day). Starting at 10 weeks of age, KDS12025 was administered via ad libitum *drinking* at doses of 1 mg/kg/day and 10 mg/kg/day. Control and SOD1 groups received standard drinking water. Motor function was evaluated weekly using the rotarod performance test. The running time ratio was recorded to monitor motor coordination and endurance over time. The survival of the mice was tracked and recorded. Median survival times were calculated for each group.

### Immunohistochemistry

Mice were deeply anesthetized with 2% avertin and positioned at the operating table. After opening up the abdomen and exposing the heart, perfused with saline followed by 4% paraformaldehyde (PFA). Decapitate the mice and the brains were excised from the skull. Excised brains were postfixed in 4% PFA overnight at 4°C and dehydrated in 30% sucrose for 48 h. Coronal hippocampal sections were cut at 30 µm-thickness in a cryostat and stored in a storage solution. The sections were fully washed in 0.1 M PBS and incubated for 1 h in a blocking solution (0.3% Triton X-100, 2% donkey and goat serum in 0.1M PBS). Next, a mixture of primary antibodies in the blocking solution was immunostained on a shaker overnight at 4°C. Sections were washed in PBS three times. After washing, sections were incubated with the corresponding fluorescent secondary antibodies for 1 h at RT. DAPI staining was done by adding a DAPI solution (1:2000) during the second washing step. After finishing the washing step, the sections were mounted on the cover slide with a mounting medium. A series of fluorescent images were obtained with an LSM900 (Zeiss, Germany) confocal microscope or Lattice SIM Elyra 7 (Zeiss, Germany). Image analysis was done by the ImageJ (NIH) program. Primary antibodies were diluted to the following amounts: chicken anti-GFAP (1:500, Millipore, ab5541), rabbit anti-amyloid beta (1:500, Abcam, ab2539), mouse anti-NeuN (1:200, Millipore, ab377), rabbit anti-Hbβ (1:200, Abcam, ab214049), mouse anti-Hbβ (1:200, Santacruz, sc-21757). Secondary antibodies from Jackson were diluted (1:500) in the blocking solution.

### Sholl analysis and neuron counting

Serial confocal sections immunolabeled with GFAP were compiled into a maximal projection. This projected image was utilized to assess the GFAP signal within the hippocampus. Utilizing the Sholl analysis plugin in ImageJ, concentric circles with 10-μm spacing were automatically generated from the nucleus to the furthest astrocytic process. This analysis quantified the intersections of GFAP-labeled processes within each circle and calculated the ramification index and the ending radius. In the focal GiD mouse model, the NeuN was quantified within a 50 μm² region using ImageJ for counting the number of CA1 pyramidal neurons.

### Imaris software

Raw image files were used for further analysis using Imaris software (Oxford Instruments). Manual surface reconstruction was performed based on masking GFAP, and Hbβ-positive signals in the stratum radiatum of the hippocampus. The parameters were surface details (0.6 μm); diameter of the largest sphere (2 μm); manual threshold value min (1000) diffusion transparency: 60%. After reconstruction from Imaris software, Hbβ-positive signals in the GFAP surface were quantified.

### Electrophysiology

Mice were deeply anesthetized with 3% isoflurane, followed by decapitation. The brain was quickly excised from the skull and submerged in a chilled cutting solution that contained 234 mM of sucrose; 24 mM of NaHCO3; 11 mM of d(+)-glucose; 10 mM of MgSO4; 2.5 mM of KCl; 12.5 mM of NaH2PO4; and 0.5 mM of CaCl2. Transverse hippocampal (300 µm thick) were prepared with a vibrating microtome (PRO7N; DSK) and transferred to an artificial cerebrospinal fluid (aCSF) solution that contained 130 mM of NaCl; 24 mM of NaHCO3; 1.25 mM of NaH2PO4; 3.5 mM KCl; 1.5 mM of CaCl2; 1.5 mM of MgCl2; and 10 mM of d(+)-glucose, pH 7.4. Slices were incubated at RT for at least 1 h before recording. All solutions were saturated with 95% O2 and 5% CO2.

For a recording of tonic GABA currents, slices were transferred to a recording chamber that was mounted on an upright Zeiss microscope and continuously perfused with the aCSF solution. The slice was viewed with a 60x water immersion objective (numerical aperture= 90) with infrared differential interference contrast optics. Cellular CMOS camera (Hamamatsu Photonics, Japan). Cellular morphology was visualized by a CMOS camera (Hamamatsu Photonics, Japan) and Imaging Workbench software (INDEC Biosystems). Whole-cell patch recordings were made from pyramidal neurons located in the DG granule cells of the hippocampus. The holding potential was −70 mV. Pipette resistance was typically 6–8 MΩ and the pipette was filled with an internal solution that contained (in mM): 135 CsCl; 4 NaCl; 0.5 CaCl2; 10 HEPES; 5 EGTA; 2 Mg-ATP; 0.5 Na2-GTP; 10 QX-314; pH-adjusted to 7.2 with CsOH (278 to 285 mOsmol). The baseline current before measuring tonic current was recorded with CNQX (20 µM) and D-AP5 (50 µM). The frequency and amplitude of spontaneous inhibitory postsynaptic currents before bicuculline (50 µM) administration were detected and measured by Mini Analysis (Synaptosoft, USA).

Dentate granule cells’ synaptic responses were elicited at a low frequency of 0.1 Hz by stimulating perforant path fibers. The stimulation was delivered over 100 ms at intensities ranging from 0 to 1,000 μA using a constant current isolation unit. A tungsten bipolar electrode was positioned in the molecular layer to stimulate the perforant path fibers. Recordings of the evoked excitatory postsynaptic potentials (EPSPs) were typically 6–8 MΩ and the pipette was filled with an internal solution that contained (in mM): 120 potassium gluconates; 10 KCl; 1 MgCl2; 0.5 EGTA; 40 HEPES; pH-adjusted to 7.2 with KOH. The spiking probability was determined by the number of spikes generated per total stimulation. Electrical signals were digitized and sampled at 10-ms intervals with the Digidata 1550B data acquisition system and the Multiclamp 700B Amplifier (Molecular Devices, USA) using the pClamp software.

### Hbβ-shRNA vector and reverse transcription-PCR

The shRNA sequence for *hbb* was designed with a Broad Institute designer (UK). shRNA for mouse *hbb* (NM_001278161.1) was targeted and shNRA sequence is 5’-gccctctgctatcatgggtaat −3’inserted into the pSico and pSicoR system^57^. Gene silencing of *hbb* was tested by reverse transcription-PCR (5’-gcacctgactgatgctgaga-3’). For gene silencing of *hbb*, adenovirus-associated virus (AAV) carrying Hbβ-shRNA was transfected into primary cultured astrocytes. 4 days after infection, the total RNA was prepared using an AllPrep RNA kit (Qiagen, Netherlands), and cDNA was synthesized by SuperScript III Reverse Transcriptase (Invitrogen, USA).

### Western blot

Western blotting was conducted following established protocols. Membranes were probed with primary antibodies targeting Hbβ and GAPDH overnight at 4 °C. Post three washes in Tris-buffered saline containing 0.05% Tween 20, and blots were incubated with HRP-conjugated secondary antibodies for 2 hours at room temperature. Bands were detected using Immobilon Western ECL solution and captured on an Image Station 4000MM.

### Immunohistochemistry with Double Chromogenic Staining

#### Human brain samples

Neuropathological examination of postmortem brain samples from normal subjects and AD patients was determined using procedures previously established by the Boston University Alzheimer’s Disease Center (BUADC). Next of kin provided informed consent for participation and brain donation. Institutional review board approval for ethical permission was obtained through the BUADC center. This study was reviewed by the Institutional Review Board of the Boston University School of Medicine and was approved for exemption because it only included tissues collected from post-mortem subjects not classified as human subjects. The study was performed in accordance with institutional regulatory guidelines and principles of human subject protection in the Declaration of Helsinki. The sample information is listed in Supplementary Table 6.

#### First staining

Paraffin-embedded human postmortem brain tissues were sectioned in a coronal plane at 10μm. BLOXALL® Blocking solution (Vector Laboratories, Burlingame, CA, USA) was used to block endogenous alkaline phosphatase. Hippocampal tissue sections were blocked with 2.5% normal horse serum (Vector Laboratories) for one h and then incubated with GFAP antibody (1:400, AB5541, Millipore, Burlington, MA, USA) for 24 h. After washing three times with PBS, tissue slides were processed with Vector ABC Kit (Vector Laboratories, Burlingame, CA, USA). The GFAP immunoreactive signals were developed with DAB chromogen (Thermo Fisher Scientific, Meridian, Rockford, IL, USA).

#### Second staining

Tissue slides stained with GFAP were incubated with hemoglobin beta (Hbβ) antibody (1:50, sc-21757, Santa Cruz) for 24 h. After reaction with secondary antibodies, sections were incubated with ImmPRESS-AP anti-mouse IgG (alkaline phosphatase) polymer detection reagent (Vector Laboratories: MP-5402) for 2 h at room temperature. A Vector Blue alkaline phosphatase substrate kit (Vector Laboratories: SK-5300) was used to develop HBβ signals. Double-stained tissue slides were gradually processed back to Histo-clear (HS-200) through an increasing ethanol gradient [70%, 80%, 90%, 95%, and 100% (1 time)] and subsequently mounted. The chromogenic signals of GFAP (brown) and Hbβ (blue) were examined under light microscopy (BX63) (Olympus, Japan) equipped with a high-definition (1920 x 1200 pixel) digital camera (DP74) (Olympus, Japan).

---

**Supplementary Table 1.** SAR analysis of HTPEB.

| Cmpd. | Structure | HB-EC<sub>50</sub> | HRP-EC<sub>50</sub> |
| --- | --- | --- | --- |
| Salicylic Acid |  | >10 | >10 |
| p-Cresotic acid |  | >10 | >10 |
| Mesalazine |  | 0.2129 | 3.604 |
| 2-(4-(Trifluoromethyl)phenyl)ethylamine |  | >10 | >10 |

<sup>a</sup> Structure of HTPEB

**Supplementary Table 2.**
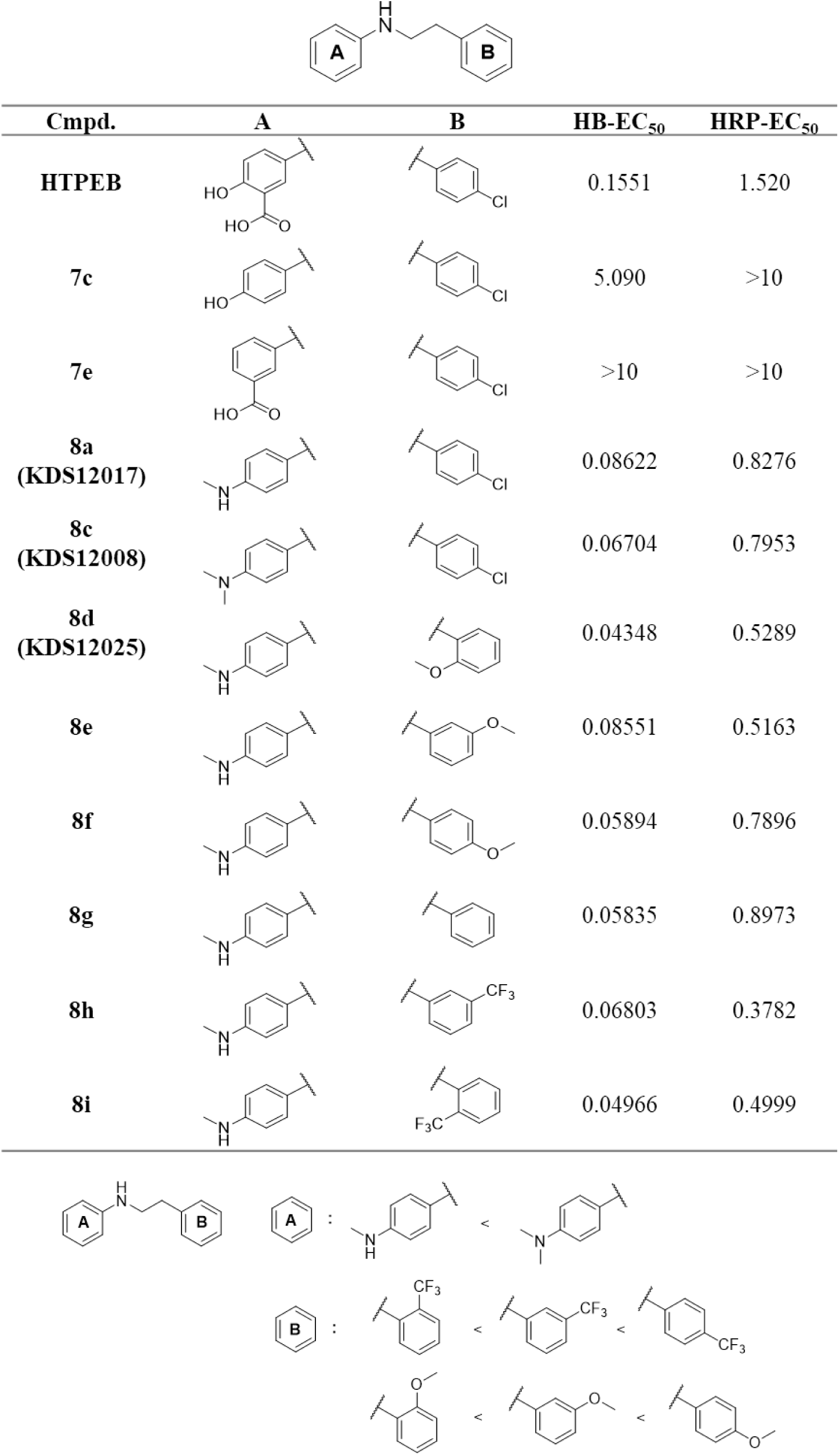
Effects of synthesized compounds on H_2_O_2_ assay.

| Cmpd. | A | B | HB-EC<sub>50</sub> | HRP-EC<sub>50</sub> |
| --- | --- | --- | --- | --- |
| HTPEB |  |  | 0.1551 | 1.520 |
| 7c |  |  | 5.090 | >10 |
| 7e |  |  | >10 | >10 |
| 8a<br>(KDS12017) |  |  | 0.08622 | 0.8276 |
| 8c<br>(KDS12008) |  |  | 0.06704 | 0.7953 |
| 8d<br>(KDS12025) |  |  | 0.04348 | 0.5289 |
| 8e |  |  | 0.08551 | 0.5163 |
| 8f |  |  | 0.05894 | 0.7896 |
| 8g |  |  | 0.05835 | 0.8973 |
| 8h |  |  | 0.06803 | 0.3782 |
| 8i |  |  | 0.04966 | 0.4999 |

**Supplementary Table 3.**
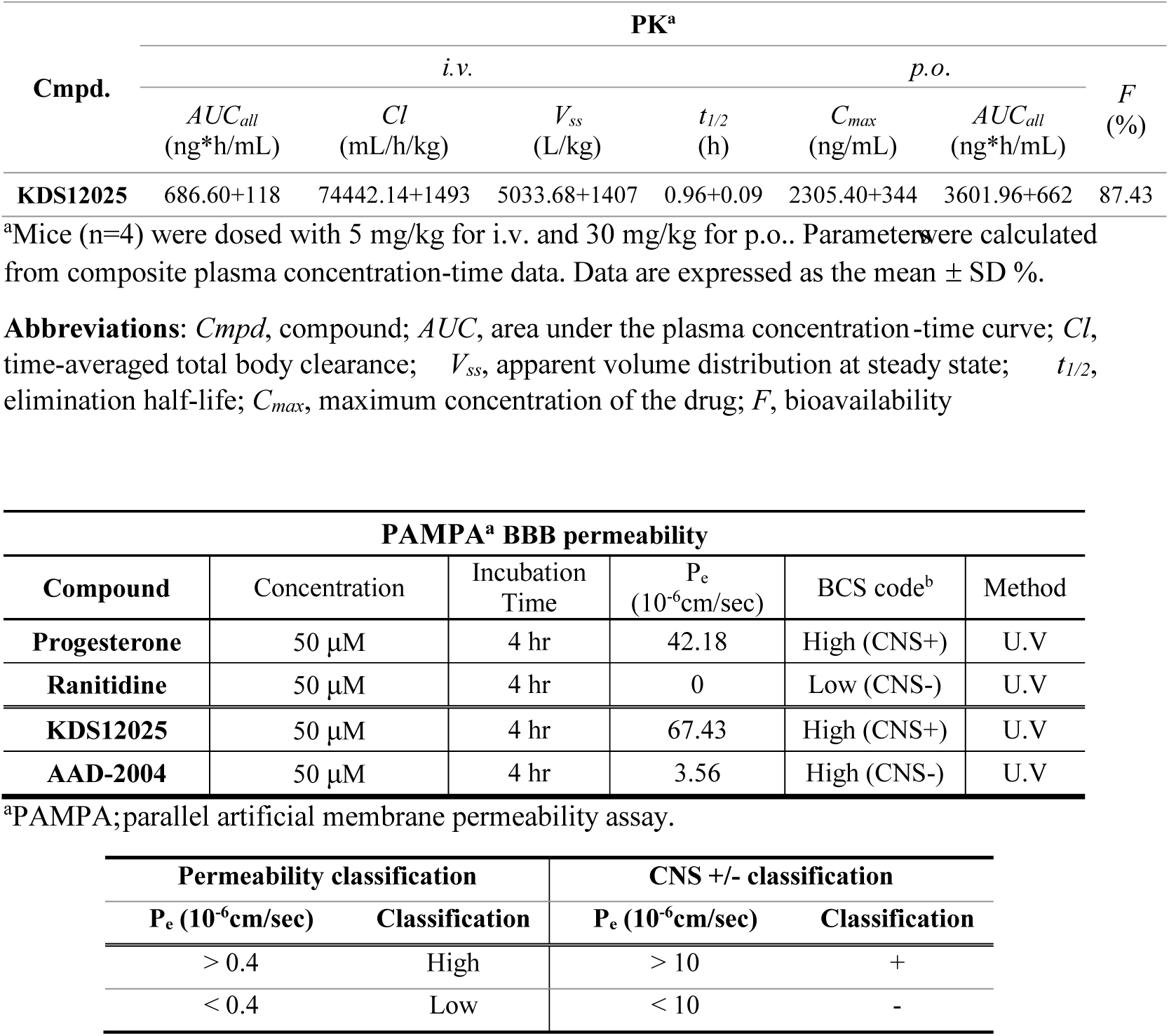
*In vivo* pharmacokinetic parameters of KDS12025.

**Supplementary Table 4.**
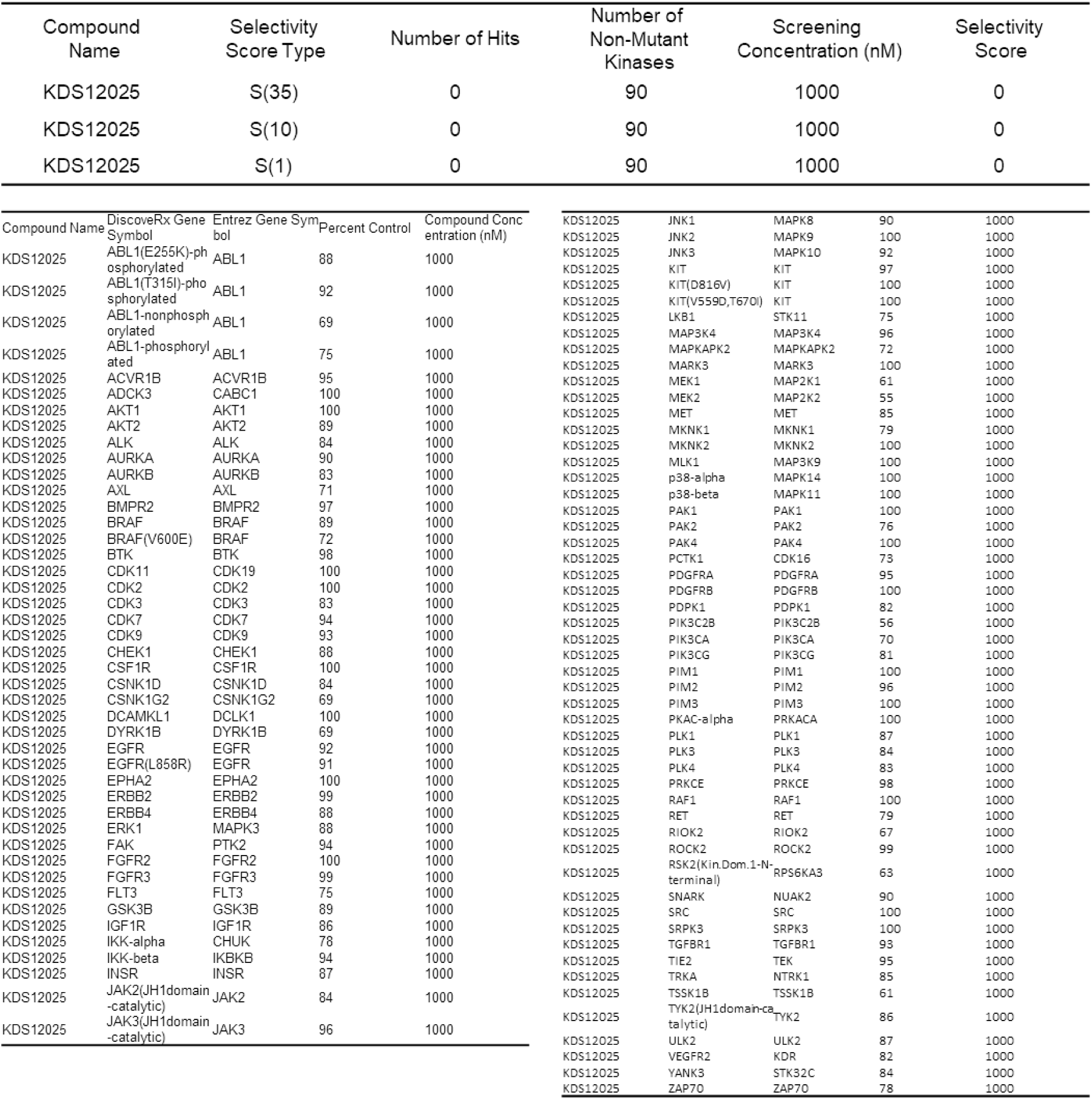
KINOMEscan screening results for KDS12025: interactions with 97 kinases.

**Supplementary Table 5.** KDS12025 interactions with 87 primary molecular targets, including G protein-coupled receptors (GPCRs), kinases, non-kinase enzymes, nuclear receptors, transporters, and various ion channels *(continued)*

| Compound I.D. | Client Compound I.D. | Test Concentration | % Inhibition of Control Specific Binding |  |  |
| --- | --- | --- | --- | --- | --- |
|  |  |  | 1<sup>st</sup> | 2<sup>nd</sup> | Mean |
| **A<sub>2A</sub>(h) (agonist radioligand)** |  |  |  |  |  |
| 100059830-2 | KDS12025 | 1.0E-06 M | 9.7 | 8.8 | 9.3 |
| **α<sub>1A</sub>(h) (antagonist radioligand)** |  |  |  |  |  |
| 100059830-2 | KDS12025 | 1.0E-06 M | 12.4 | 13.5 | 12.9 |
| **α<sub>1B</sub>(h) (antagonist radioligand)** |  |  |  |  |  |
| 100059830-2 | KDS12025 | 1.0E-06 M | 38.6 | 30.6 | 34.6 |
| **α<sub>1D</sub>(h) (antagonist radioligand)** |  |  |  |  |  |
| 100059830-2 | KDS12025 | 1.0E-06 M | 24.7 | 23.7 | 24.2 |
| **α<sub>2A</sub>(h) (antagonist radioligand)** |  |  |  |  |  |
| 100059830-2 | KDS12025 | 1.0E-06 M | 13.1 | 14.6 | 13.9 |
| **α<sub>2B</sub>(h) (antagonist radioligand)** |  |  |  |  |  |
| 100059830-2 | KDS12025 | 1.0E-06 M | 18.1 | 13.4 | 15.8 |
| **β<sub>1</sub>(h) (agonist radioligand)** |  |  |  |  |  |
| 100059830-2 | KDS12025 | 1.0E-06 M | 13.3 | 5.7 | 9.5 |
| **β<sub>2</sub>(h) (antagonist radioligand)** |  |  |  |  |  |
| 100059830-2 | KDS12025 | 1.0E-06 M | 3.2 | -3.9 | -0.3 |
| **AT<sub>1</sub> (h) (antagonist radioligand)** |  |  |  |  |  |
| 100059830-2 | KDS12025 | 1.0E-06 M | 17.7 | 7.5 | 12.6 |
| **BZD (central) (agonist radioligand)** |  |  |  |  |  |
| 100059830-2 | KDS12025 | 1.0E-06 M | -2.5 | 3.4 | 0.5 |
| **Cl<sup>-</sup> channel (GABA-gated) (TBOB site) (antagonist radioligand)** |  |  |  |  |  |
| 100059830-2 | KDS12025 | 1.0E-06 M | -3.8 | 11.0 | 3.6 |
| **B<sub>2</sub> (h) (agonist radioligand)** |  |  |  |  |  |
| 100059830-2 | KDS12025 | 1.0E-06 M | -4.5 | 0.6 | -2.0 |
| **CB<sub>1</sub>(h) (agonist radioligand)** |  |  |  |  |  |
| 100059830-2 | KDS12025 | 1.0E-06 M | 18.3 | 21.6 | 20.0 |
| **CB<sub>2</sub>(h) (agonist radioligand)** |  |  |  |  |  |
| 100059830-2 | KDS12025 | 1.0E-06 M | 21.6 | 9.2 | 15.4 |
| **CCK<sub>1</sub> (CCK<sub>A</sub>) (h) (agonist radioligand)** |  |  |  |  |  |
| 100059830-2 | KDS12025 | 1.0E-06 M | -3.7 | -4.5 | -4.1 |
| **CCK<sub>2</sub> (CCK<sub>B</sub>) (h) (agonist radioligand)** |  |  |  |  |  |
| 100059830-2 | KDS12025 | 1.0E-06 M | -7.1 | 3.0 | -2.1 |
| **D<sub>1</sub>(h) (antagonist radioligand)** |  |  |  |  |  |
| 100059830-2 | KDS12025 | 1.0E-06 M | 8.9 | -3.8 | 2.5 |
| **D<sub>2S</sub>(h) (agonist radioligand)** |  |  |  |  |  |
| 100059830-2 | KDS12025 | 1.0E-06 M | 36.0 | 40.2 | 38.1 |
| **D<sub>2L</sub>(h) (antagonist radioligand)** |  |  |  |  |  |
| 100059830-2 | KDS12025 | 1.0E-06 M | 25.2 | 6.8 | 16.0 |
| **ET<sub>A</sub>(h) (agonist radioligand)** |  |  |  |  |  |
| 100059830-2 | KDS12025 | 1.0E-06 M | -8.0 | 9.2 | 0.6 |
| **GABA<sub>A</sub>1 (h) (α1,β2,γ2) (agonist radioligand)** |  |  |  |  |  |
| 100059830-2 | KDS12025 | 1.0E-06 M | 7.6 | 2.2 | 4.9 |
| **GABA<sub>B</sub>(1b)(h) (antagonist radioligand)** |  |  |  |  |  |
| 100059830-2 | KDS12025 | 1.0E-06 M | -6.5 | -0.7 | -3.6 |
| **AMPA (agonist radioligand)** |  |  |  |  |  |
| 100059830-2 | KDS12025 | 1.0E-06 M | 2.0 | -2.8 | -0.4 |
| **kainate (agonist radioligand)** |  |  |  |  |  |
| 100059830-2 | KDS12025 | 1.0E-06 M | 23.2 | -18.1 | 2.6 |
| **NMDA (antagonist radioligand)** |  |  |  |  |  |
| 100059830-2 | KDS12025 | 1.0E-06 M | 2.2 | 0.4 | 1.3 |
| **mGluR5 (h) (agonist radioligand)** |  |  |  |  |  |
| 100059830-2 | KDS12025 | 1.0E-06 M | -13.6 | -15.8 | -14.7 |
|  |  |  | 1<sup>st</sup> | 2<sup>nd</sup> | Mean |
| glycine (strychnine-sensitive) (antagonist radioligand) |  |  |  |  |  |
| 100059830-2 | KDS12025 | 1.0E-06 M | -17.4 | -17.6 | -17.5 |
| glycine (strychnine-insensitive) (antagonist radioligand) |  |  |  |  |  |
| 100059830-2 | KDS12025 | 1.0E-06 M | 3.2 | -0.5 | 1.4 |
| CXCR2 (IL-8B) (h) (agonist radioligand) |  |  |  |  |  |
| 100059830-2 | KDS12025 | 1.0E-06 M | 8.2 | 16.7 | 12.4 |
| CCR1 (h) (agonist radioligand) |  |  |  |  |  |
| 100059830-2 | KDS12025 | 1.0E-06 M | 9.6 | 11.8 | 10.7 |
| H<sub>1</sub>(h) (antagonist radioligand) |  |  |  |  |  |
| 100059830-2 | KDS12025 | 1.0E-06 M | 12.9 | 10.6 | 11.8 |
| H<sub>2</sub>(h) (antagonist radioligand) |  |  |  |  |  |
| 100059830-2 | KDS12025 | 1.0E-06 M | -12.8 | 5.8 | -3.5 |
| CysLT<sub>1</sub> (LTD<sub>4</sub>)(h) (agonist radioligand) |  |  |  |  |  |
| 100059830-2 | KDS12025 | 1.0E-06 M | 16.9 | 9.0 | 12.9 |
| MC<sub>1</sub> (agonist radioligand) |  |  |  |  |  |
| 100059830-2 | KDS12025 | 1.0E-06 M | -16.6 | -1.8 | -9.2 |
| MC<sub>4</sub>(h) (agonist radioligand) |  |  |  |  |  |
| 100059830-2 | KDS12025 | 1.0E-06 M | 11.3 | 14.5 | 12.9 |
| MAO-A (antagonist radioligand) |  |  |  |  |  |
| 100059830-2 | KDS12025 | 1.0E-06 M | 4.5 | -5.1 | -0.3 |
| M<sub>1</sub>(h) (antagonist radioligand) |  |  |  |  |  |
| 100059830-2 | KDS12025 | 1.0E-06 M | 2.2 | 14.1 | 8.2 |
| M<sub>2</sub> (h) (antagonist radioligand) |  |  |  |  |  |
| 100059830-2 | KDS12025 | 1.0E-06 M | -4.1 | -10.2 | -7.1 |
| M<sub>3</sub>(h) (antagonist radioligand) |  |  |  |  |  |
| 100059830-2 | KDS12025 | 1.0E-06 M | 11.8 | 17.3 | 14.5 |
| M<sub>4</sub>(h) (antagonist radioligand) |  |  |  |  |  |
| 100059830-2 | KDS12025 | 1.0E-06 M | 1.0 | -4.5 | -1.7 |
| NK<sub>1</sub>(h) (agonist radioligand) |  |  |  |  |  |
| 100059830-2 | KDS12025 | 1.0E-06 M | 7.8 | 9.9 | 8.8 |
| Y<sub>1</sub>(h) (agonist radioligand) |  |  |  |  |  |
| 100059830-2 | KDS12025 | 1.0E-06 M | -5.5 | -2.1 | -3.8 |
| N neuronal α4β2 (h) (agonist radioligand) |  |  |  |  |  |
| 100059830-2 | KDS12025 | 1.0E-06 M | 1.5 | 1.7 | 1.6 |
| N muscle-type (h) (antagonist radioligand) |  |  |  |  |  |
| 100059830-2 | KDS12025 | 1.0E-06 M | 31.5 | 20.0 | 25.7 |
| δ (DOP) (h) (agonist radioligand) |  |  |  |  |  |
| 100059830-2 | KDS12025 | 1.0E-06 M | 10.6 | 30.0 | 20.3 |
| kappa (h) (KOP) (agonist radioligand) |  |  |  |  |  |
| 100059830-2 | KDS12025 | 1.0E-06 M | 22.1 | 12.4 | 17.3 |
| μ (MOP) (h) (agonist radioligand) |  |  |  |  |  |
| 100059830-2 | KDS12025 | 1.0E-06 M | 14.5 | 16.5 | 15.5 |
| PPAR<sub>γ</sub> (h) (agonist radioligand) |  |  |  |  |  |
| 100059830-2 | KDS12025 | 1.0E-06 M | 5.3 | -4.6 | 0.3 |
| PAF (h) (agonist radioligand) |  |  |  |  |  |
| 100059830-2 | KDS12025 | 1.0E-06 M | -6.0 | -6.1 | -6.0 |
| PCP (antagonist radioligand) |  |  |  |  |  |
| 100059830-2 | KDS12025 | 1.0E-06 M | 5.4 | 10.6 | 8.0 |
| RARα (h) (agonist radioligand) |  |  |  |  |  |
| 100059830-2 | KDS12025 | 1.0E-06 M | -5.4 | -5.0 | -5.2 |
| 5-HT<sub>1A</sub>(h) (agonist radioligand) |  |  |  |  |  |
| 100059830-2 | KDS12025 | 1.0E-06 M | 34.9 | 40.4 | 37.6 |
|  |  |  | 1<sup>st</sup> | 2<sup>nd</sup> | Mean |
| 5-HT<sub>1B</sub> (h) (antagonist radioligand) |  |  |  |  |  |
| 100059830-2 | KDS12025 | 1.0E-06 M | 13.2 | 5.3 | 9.2 |
| 5-HT<sub>2A</sub>(h) (agonist radioligand) |  |  |  |  |  |
| 100059830-2 | KDS12025 | 1.0E-06 M | 22.5 | 13.9 | 18.2 |
| 5-HT<sub>2B</sub>(h) (agonist radioligand) |  |  |  |  |  |
| 100059830-2 | KDS12025 | 1.0E-06 M | 34.1 | 38.2 | 36.1 |
| 5-HT<sub>2C</sub>(h) (antagonist radioligand) |  |  |  |  |  |
| 100059830-2 | KDS12025 | 1.0E-06 M | 21.0 | 26.2 | 23.6 |
| 5-HT<sub>3</sub>(h) (antagonist radioligand) |  |  |  |  |  |
| 100059830-2 | KDS12025 | 1.0E-06 M | 8.6 | -0.5 | 4.0 |
| GR (h) (agonist radioligand) |  |  |  |  |  |
| 100059830-2 | KDS12025 | 1.0E-06 M | -4.3 | -3.4 | -3.9 |
| Estrogen ER alpha (h) (agonist radioligand) |  |  |  |  |  |
| 100059830-2 | KDS12025 | 1.0E-06 M | -9.2 | 18.5 | 4.7 |
| PR (h) (agonist radioligand) |  |  |  |  |  |
| 100059830-2 | KDS12025 | 1.0E-06 M | -5.3 | -2.8 | -4.1 |
| AR(h) (agonist radioligand) |  |  |  |  |  |
| 100059830-2 | KDS12025 | 1.0E-06 M | 5.8 | 12.5 | 9.1 |
| V<sub>1a</sub>(h) (agonist radioligand) |  |  |  |  |  |
| 100059830-2 | KDS12025 | 1.0E-06 M | -12.5 | -7.3 | -9.9 |
| Ca<sup>2+</sup> channel (L, dihydropyridine site) (antagonist radioligand) |  |  |  |  |  |
| 100059830-2 | KDS12025 | 1.0E-06 M | -37.5 | -21.0 | -29.3 |
| Ca<sup>2+</sup> channel (L, diltiazem site) (benzothiazepines) (antagonist radioligand) |  |  |  |  |  |
| 100059830-2 | KDS12025 | 1.0E-06 M | 11.9 | 18.3 | 15.1 |
| Ca<sup>2+</sup> channel (L, verapamil site) (phenylalkylamine) (antagonist radioligand) |  |  |  |  |  |
| 100059830-2 | KDS12025 | 1.0E-06 M | -0.6 | 4.5 | 2.0 |
| Ca<sup>2+</sup> channel (N) (antagonist radioligand) |  |  |  |  |  |
| 100059830-2 | KDS12025 | 1.0E-06 M | -3.6 | -17.7 | -10.7 |
| Potassium Channel hERG (human)- [3H] Dofetilide |  |  |  |  |  |
| 100059830-2 | KDS12025 | 1.0E-06 M | 29.7 | 33.2 | 31.5 |
| K<sub>V</sub> channel (antagonist radioligand) |  |  |  |  |  |
| 100059830-2 | KDS12025 | 1.0E-06 M | 14.7 | -7.8 | 3.5 |
| Na<sup>+</sup> channel (site 2) (antagonist radioligand) |  |  |  |  |  |
| 100059830-2 | KDS12025 | 1.0E-06 M | 16.4 | 34.5 | 25.5 |
| adenosine transporter (antagonist radioligand) |  |  |  |  |  |
| 100059830-2 | KDS12025 | 1.0E-06 M | 1.4 | -1.4 | 0.0 |
| norepinephrine transporter(h) (antagonist radioligand) |  |  |  |  |  |
| 100059830-2 | KDS12025 | 1.0E-06 M | 10.5 | 3.0 | 6.7 |
| dopamine transporter(h) (antagonist radioligand) |  |  |  |  |  |
| 100059830-2 | KDS12025 | 1.0E-06 M | -1.9 | -11.4 | -6.6 |
| GABA transporter (antagonist radioligand) |  |  |  |  |  |
| 100059830-2 | KDS12025 | 1.0E-06 M | -1.0 | 8.9 | 4.0 |
| 5-HT transporter (h) (antagonist radioligand) |  |  |  |  |  |
| 100059830-2 | KDS12025 | 1.0E-06 M | 37.1 | 53.6 | 45.4 |

**Supplementary Table 6.** Demographic information about human postmortem brain samples from normal subjects and AD patients.

| Number | Case | Age | Sex | Braak Stage |
| --- | --- | --- | --- | --- |
| 1 | Normal | 88 | M | I |
| 2 | Normal | 82 | M | I |
| 3 | Normal | 70 | M | I |
| 1 | AD | 89 | M | V |
| 2 | AD | 83 | M | VI |
| 3 | AD | 70 | M | VI |

**Supplementary Table 7.** Additional statics table.

| Figure No. | Detailed information for statistical analysis |
| --- | --- |
| 2b | Two-way ANOVA with Tukey's multiple comparison test, $F(1,21) = 0.6535$, $p = 0.43$<br>HRP: Vehicle (1.001±0.02269, n=3), HTPeB (3.769±0.7911, n=3), KDS12025 (5.642±0.5747, n=3)<br>Vehicle vs. HTPeB, $p = 0.03$; Vehicle vs. KDS12025, $p < 0.001$; HTPeB vs. KDS12025, $p = 0.17$<br>Hb: Vehicle (1.001±0.01076, n=6), HTPeB (2.529±0.6645, n=6), KDS12025 (5.676±0.6374, n=6)<br>Vehicle vs. HTPeB, $p = 0.10$; Vehicle vs. KDS12025, $p < 0.001$; HTPeB vs. KDS12025, $p < 0.001$ |
| 2d | Hb (0.01 U/ml): (-) (100.0±1.860, n=3), Hb+DMSO (97.77±0.9081, n=3),<br>Hb+HTPeB (84.92±3.034, n=3), Hb+KDS12025 (82.57±2.025, n=3)<br>One-way ANOVA with Tukey's multiple comparisons test, $F(3,8) = 17.77$, $p < 0.001$<br>(-) vs. Hb+DMSO, $p = 0.87$; (-) vs. Hb+HTPeB, $p = 0.004$; (-) vs. Hb+KDS12025, $p = 0.002$;<br>Hb+DMSO vs. Hb+HTPeB, $p = 0.01$; Hb+DMSO vs. Hb+KDS12025, $p = 0.004$; Hb+HTPeB vs. Hb+KDS12025, $p = 0.86$<br>Hb (0.1 U/ml): (-) (100.0±1.860, n=3), Hb+DMSO (95.19±1.581, n=3),<br>Hb+HTPeB (81.24±5.607, n=3), Hb+KDS12025 (75.36±1.113, n=3)<br>One-way ANOVA with Tukey's multiple comparisons test, $F(3,8) = 13.84$, $p = 0.002$<br>(-) vs. Hb+DMSO, $p = 0.7$; (-) vs. Hb+HTPeB, $p = 0.01$; (-) vs. Hb+KDS12025, $p = 0.002$;<br>Hb+DMSO vs. Hb+HTPeB, $p = 0.05$; Hb+DMSO vs. Hb+KDS12025, $p = 0.009$; Hb+HTPeB vs. Hb+KDS12025, $p = 0.57$<br>Hb (1 U/ml): (-) (100.0±5.136, n=3), Hb+DMSO (91.00±3.892, n=6),<br>Hb+HTPeB (70.82±3.83, n=6), Hb+KDS12025 (62.38±2.923, n=6)<br>One-way ANOVA with Tukey's multiple comparisons test, $F(3,17) = 18.34$, $p < 0.001$<br>(-) vs. Hb+DMSO, $p = 0.49$; (-) vs. Hb+HTPeB, $p = 0.001$; (-) vs. Hb+KDS12025, $p < 0.001$;<br>Hb+DMSO vs. Hb+HTPeB, $p = 0.005$; Hb+DMSO vs. Hb+KDS12025, $p < 0.001$; Hb+HTPeB vs. Hb+KDS12025, $p = 0.37$<br>Hb (11 U/ml): (-) (100.0±5.009, n=6), Hb+DMSO (64.68±6.780, n=9),<br>Hb+HTPeB (48.10±5.928, n=9), Hb+KDS12025 (30.17±2.742, n=9)<br>One-way ANOVA with Tukey's multiple comparisons test, $F(3,29) = 25.49$, $p < 0.001$<br>(-) vs. Hb+DMSO, $p = 0.001$; (-) vs. Hb+HTPeB, $p < 0.001$; (-) vs. Hb+KDS12025, $p < 0.001$;<br>Hb+DMSO vs. Hb+HTPeB, $p = 0.14$; Hb+DMSO vs. Hb+KDS12025, $p < 0.001$; Hb+HTPeB vs. Hb+KDS12025, $p = 0.09$<br>Hb (112 U/ml): (-) (100.0±5.968, n=3), Hb+DMSO (22.45±2.044, n=6),<br>Hb+HTPeB (13.83±1.029, n=6), Hb+KDS12025 (8.191±0.4716, n=6)<br>One-way ANOVA with Tukey's multiple comparisons test, $F(3,17) = 289.3$, $p < 0.001$<br>(-) vs. Hb+DMSO, $p < 0.001$; (-) vs. Hb+HTPeB, $p < 0.001$; (-) vs. Hb+KDS12025, $p < 0.001$;<br>Hb+DMSO vs. Hb+HTPeB, $p = 0.03$; Hb+DMSO vs. Hb+KDS12025, $p < 0.001$; Hb+HTPeB vs. Hb+KDS12025, $p = 0.20$ |
| 2g | Two-way ANOVA with Sidak's multiple comparison test, $F(6,24) = 0.1095$, $p > 0.99$<br>Full day: CTRL (3927, n=4), KDS12025 (10mpk) (3593, n=4), KDS12025 (1mpk) (3871, n=4), KDS12025 (0.1mpk) (4161, n=4)<br>CTRL vs. KDS12025 (10mpk), $p > 0.99$; CTRL vs. KDS12025 (1mpk), $p > 0.99$; CTRL vs. KDS12025 (0.1mpk), $p > 0.99$; KDS12025 (10mpk) vs. KDS12025 (1mpk), $p > 0.99$; KDS12025 (10mpk) vs. KDS12025 (0.1mpk), $p > 0.99$; KDS12025 (1mpk) vs. KDS12025 (0.1mpk), $p > 0.99$<br>Dark: CTRL (4162, n=4), KDS12025 (10mpk) (4216, n=4), KDS12025 (1mpk) (4063, n=4), KDS12025 (0.1mpk) (4228, n=4)<br>CTRL vs. KDS12025 (10mpk), $p > 0.99$; CTRL vs. KDS12025 (1mpk), $p > 0.99$; CTRL vs. KDS12025 (0.1mpk), $p > 0.99$; KDS12025 (10mpk) vs. KDS12025 (1mpk), $p > 0.99$; KDS12025 (10mpk) vs. KDS12025 (0.1mpk), $p > 0.99$; KDS12025 (1mpk) vs. KDS12025 (0.1mpk), $p > 0.99$<br>Light: CTRL (3752, n=4), KDS12025 (10mpk) (3753, n=4), KDS12025 (1mpk) (3584, n=4), KDS12025 (0.1mpk) (3923, n=4)<br>CTRL vs. KDS12025 (10mpk), $p > 0.99$; CTRL vs. KDS12025 (1mpk), $p > 0.99$; CTRL vs. KDS12025 (0.1mpk), $p > 0.99$; KDS12025 (10mpk) vs. KDS12025 (1mpk), $p > 0.99$; KDS12025 (10mpk) vs. KDS12025 (0.1mpk), $p > 0.99$; KDS12025 (1mpk) vs. KDS12025 (0.1mpk), $p > 0.99$ |
| 2h | Two-way ANOVA with Sidak's multiple comparison test, $F(6,24) = 0.1095$, $p > 0.99$<br>Full day: CTRL (3072, n=4), KDS12025 (10mpk) (3120, n=4), KDS12025 (1mpk) (3075, n=4), KDS12025 (0.1mpk) (3292, n=4)<br>CTRL vs. KDS12025 (10mpk), $p > 0.99$; CTRL vs. KDS12025 (1mpk), $p > 0.99$; CTRL vs. KDS12025 (0.1mpk), $p > 0.99$; KDS12025 (10mpk) vs. KDS12025 (1mpk), $p > 0.99$; KDS12025 (10mpk) vs. KDS12025 (0.1mpk), $p > 0.99$; KDS12025 (1mpk) vs. KDS12025 (0.1mpk), $p > 0.99$<br>Dark: CTRL (3299, n=4), KDS12025 (10mpk) (3344, n=4), KDS12025 (1mpk) (3167, n=4), KDS12025 (0.1mpk) (3345, n=4)<br>CTRL vs. KDS12025 (10mpk), $p > 0.99$; CTRL vs. KDS12025 (1mpk), $p > 0.99$; CTRL vs. KDS12025 (0.1mpk), $p > 0.99$; KDS12025 (10mpk) vs. KDS12025 (1mpk), $p > 0.99$; KDS12025 (10mpk) vs. KDS12025 (0.1mpk), $p > 0.99$; KDS12025 (1mpk) vs. KDS12025 (0.1mpk), $p > 0.99$<br>Light: CTRL (2850, n=4), KDS12025 (10mpk) (2900, n=4), KDS12025 (1mpk) (2916, n=4), KDS12025 (0.1mpk) (3065, n=4)<br>CTRL vs. KDS12025 (10mpk), $p > 0.99$; CTRL vs. KDS12025 (1mpk), $p > 0.99$; CTRL vs. KDS12025 (0.1mpk), $p > 0.99$; KDS12025 (10mpk) vs. KDS12025 (1mpk), $p > 0.99$; KDS12025 (10mpk) vs. KDS12025 (0.1mpk), $p > 0.99$; KDS12025 (1mpk) vs. KDS12025 (0.1mpk), $p > 0.99$ |
| 2i | Two-way ANOVA with Sidak's multiple comparison test, $F(6,24) = 0.1095$, $p > 0.99$<br>Full day: CTRL (0.834, n=4), KDS12025 (10mpk) (0.857, n=4), KDS12025 (1mpk) (0.883, n=4), KDS12025 (0.1mpk) (0.857, n=4)<br>CTRL vs. KDS12025 (10mpk), $p > 0.99$; CTRL vs. KDS12025 (1mpk), $p > 0.99$; CTRL vs. KDS12025 (0.1mpk), $p > 0.99$; KDS12025 (10mpk) vs. KDS12025 (1mpk), $p > 0.99$; KDS12025 (10mpk) vs. KDS12025 (0.1mpk), $p > 0.99$; KDS12025 (1mpk) vs. KDS12025 (0.1mpk), $p > 0.99$<br>Dark: CTRL (0.858, n=4), KDS12025 (10mpk) (0.877, n=4), KDS12025 (1mpk) (0.891, n=4), KDS12025 (0.1mpk) (0.849, n=4)<br>CTRL vs. KDS12025 (10mpk), $p > 0.99$; CTRL vs. KDS12025 (1mpk), $p > 0.99$; CTRL vs. KDS12025 (0.1mpk), $p > 0.99$; KDS12025 (10mpk) vs. KDS12025 (1mpk), $p > 0.99$; KDS12025 (10mpk) vs. KDS12025 (0.1mpk), $p > 0.99$; KDS12025 (1mpk) vs. KDS12025 (0.1mpk), $p > 0.99$<br>Light: CTRL (0.8033, n=4), KDS12025 (10mpk) (0.831, n=4), KDS12025 (1mpk) (0.860, n=4), KDS12025 (0.1mpk) (0.833, n=4)<br>CTRL vs. KDS12025 (10mpk), $p > 0.99$; CTRL vs. KDS12025 (1mpk), $p > 0.99$; CTRL vs. KDS12025 (0.1mpk), $p > 0.99$; KDS12025 (10mpk) vs. KDS12025 (1mpk), $p > 0.99$; KDS12025 (10mpk) vs. KDS12025 (0.1mpk), $p > 0.99$; KDS12025 (1mpk) vs. KDS12025 (0.1mpk), $p > 0.99$ |
| 3b | Two-way ANOVA with Tukey's multiple comparison test, $F(2,27) = 1.662$, $p = 0.21$<br>Acquisition: WT+GFP (60.65±15.92, n=13), APP+GFP (37.86±9.449, n=7), APP+DTR+KDS (29.44±7.549, n=10)<br>WT+GFP vs. APP+GFP, $p = 0.96$; WT+GFP vs. APP+DTR+KDS, $p = 0.88$; APP+DTR vs. APP+DTR+KDS, $p > 0.99$<br>Retrieval: WT (364.8±47.97, n=13), APP+GFP (107.6±44.34, n=7), APP+DTR+KDS (281.8±47.93, n=9)<br>WT+GFP vs. APP+GFP, $p < 0.001$; WT+GFP vs. APP+DTR+KDS, $p = 0.25$; APP+DTR vs. APP+DTR+KDS, $p = 0.009$ |
| 3c | WT+GFP (0.1807±0.06871, n=12), APP+DTR (-0.04320±0.04714, n=14), APP+DTR+KDS (0.3678±0.1108, n=6)<br>One-way ANOVA with Tukey's multiple comparisons test, $F(2,29) = 8.158$, $p = 0.002$<br>WT+GFP vs. APP+DTR, $p = 0.04$; WT+GFP vs. APP+DTR+KDS, $p = 0.22$; APP+DTR vs. APP+DTR+KDS, $p = 0.002$ |
| 3e | WT+GFP (49.18±2.4685, n=9), APP+DTR (82.53±3.645, n=9), APP+DTR+KDS (58.99±2.456, n=9)<br>One-way ANOVA with Tukey's multiple comparisons test, $F(2,24) = 33.23$, $p < 0.001$<br>WT+GFP vs. APP+DTR, $p < 0.001$; WT+GFP vs. APP+DTR+KDS, $p = 0.07$; APP+DTR vs. APP+DTR+KDS, $p = 0.002$ |
| 3f | WT+GFP (58.95±1.050, n=11), APP+DTR (41.15±1.393, n=10), APP+DTR+KDS (54.19±2.663, n=11)<br>One-way ANOVA with Tukey's multiple comparisons test, $F(2,29) = 23.72$, $p < 0.01$<br>WT+GFP vs. APP+DTR, $p < 0.001$; WT+GFP vs. APP+DTR+KDS, $p = 0.18$; APP+DTR vs. APP+DTR+KDS, $p < 0.001$ |
| 3h | Control (224.3±30.59, n=6), A53T (93.57±19.55, n=7), A53T+KDS (223.8±23.51, n=8)<br>One-way ANOVA with Tukey's multiple comparisons test, $F(2,18) = 9.52$, $p = 0.02$<br>Control vs. A53T, $p = 0.005$; Control vs. A53T+KDS, $p > 0.99$; A53T vs. A53T+KDS, $p = 0.003$ |
| 3j | Control (12.16±1.171.59, n=35), A53T ipsi (73.73±10.13, n=34), A53T ipsi+KDS (5.531±0.9318, n=16), A53T contra (4.673±1.833, n=8), A53T contra+KDS (10.18±1.460, n=18)<br>One-way ANOVA with Tukey's multiple comparisons test, F(4,106) = 22.01, p < 0.001<br>Control vs. A53T ipsi, p < 0.001; Control vs. A53T ipsi+KDS, p = 0.96; Control vs. A53T contra, p = 0.98; Control vs. A53T contra+KDS, p > 0.99; A53T ipsi vs. A53T ipsi+KDS, p < 0.001; A53T ipsi vs. A53T contra, p < 0.001; A53T ipsi vs. A53T contra+KDS, p < 0.001; A53T ipsi+KDS vs. A53T contra, p > 0.99; A53T ipsi+KDS vs. A53T contra+KDS, p > 0.99; A53T contra vs. A53T contra+KDS, p > 0.99 |
| 3k | Control (26.81±3.748, n=35), A53T ipsi (57.94±4.409, n=91), A53T ipsi+KDS (34.18±3.214, n=53), A53T contra (36±6.065, n=31), A53T contra+KDS (25.28±2.892, n=39)<br>One-way ANOVA with Tukey's multiple comparisons test, F(4,244) = 11.18, p < 0.001<br>Control vs. A53T ipsi, p < 0.001; Control vs. A53T ipsi+KDS, p = 0.83; Control vs. A53T contra, p = 0.77; Control vs. A53T contra+KDS, p > 0.99; A53T ipsi vs. A53T ipsi+KDS, p < 0.001; A53T ipsi vs. A53T contra, p = 0.001; A53T ipsi vs. A53T contra+KDS, p < 0.001; A53T ipsi+KDS vs. A53T contra, p > 0.99; A53T ipsi+KDS vs. A53T contra+KDS, p = 0.68; A53T contra vs. A53T contra+KDS, p = 0.63 |
| 3l | Control (30.24±4.6, n=5), A53T ipsi (7.112±2.209, n=5), A53T ipsi+KDS (20.96±2.68, n=7), A53T contra (23.62±2.751, n=5), A53T contra+KDS (23.20±2.685, n=6)<br>One-way ANOVA with Tukey's multiple comparisons test, F(4,23) = 7.027, p < 0.001<br>Control vs. A53T ipsi, p < 0.001; Control vs. A53T ipsi+KDS, p = 0.21; Control vs. A53T contra, p = 0.60; Control vs. A53T contra+KDS, p = 0.50; A53T ipsi vs. A53T ipsi+KDS, p = 0.02; A53T ipsi vs. A53T contra, p = 0.01; A53T ipsi vs. A53T contra+KDS, p = 0.01; A53T ipsi+KDS vs. A53T contra, p = 0.97; A53T ipsi+KDS vs. A53T contra+KDS, p = 0.98; A53T contra vs. A53T contra+KDS, p > 0.99 |
| 3n | Two-way ANOVA with Tukey's multiple comparison test, F (42,303) = 3.645, p < 0.001<br>15wk: Control (n=5), SOD1 (n=6), SOD1+KDS12025 1mpk (n=7), SOD1+KDS12025 10mpk (n=7)<br>Control vs. SOD1, p = 0.0014; Control vs. SOD1+KDS12025 1mpk, p = 0.9366; Control vs. SOD1+KDS12025 10mpk, p = 0.306<br>10wk: Control vs. SOD1, p > 0.9999; Control vs. SOD1+KDS12025 1mpk, p > 0.9999; Control vs. SOD1+KDS12025 10mpk, p = 0.7843<br>11wk: Control vs. SOD1, p = 0.4748; Control vs. SOD1+KDS12025 1mpk, p = 0.1726; Control vs. SOD1+KDS12025 10mpk, p > 0.9999<br>12wk: Control vs. SOD1, p = 0.0727; Control vs. SOD1+KDS12025 1mpk, p > 0.9999; Control vs. SOD1+KDS12025 10mpk, p = 0.3824<br>13wk: Control vs. SOD1, p = 0.5437; Control vs. SOD1+KDS12025 1mpk, p = 0.9744; Control vs. SOD1+KDS12025 10mpk, p > 0.9999<br>14wk: Control vs. SOD1, p = 0.3696; Control vs. SOD1+KDS12025 1mpk, p > 0.9999; Control vs. SOD1+KDS12025 10mpk, p = 0.8831<br>15wk: Control vs. SOD1, p = 0.0014; Control vs. SOD1+KDS12025 1mpk, p = 0.9366; Control vs. SOD1+KDS12025 10mpk, p = 0.3060<br>16wk: Control vs. SOD1, p = 0.0201; Control vs. SOD1+KDS12025 1mpk, p = 0.9991; Control vs. SOD1+KDS12025 10mpk, p = 0.2009<br>17wk: Control vs. SOD1, p = 0.0007; Control vs. SOD1+KDS12025 1mpk, p = 0.9962; Control vs. SOD1+KDS12025 10mpk, p > 0.9999<br>18wk: Control vs. SOD1, p = 0.0201; Control vs. SOD1+KDS12025 1mpk, p = 0.9991; Control vs. SOD1+KDS12025 10mpk, p = 0.2009<br>19wk: Control vs. SOD1, p = 0.0007; Control vs. SOD1+KDS12025 1mpk, p = 0.9962; Control vs. SOD1+KDS12025 10mpk, p > 0.9999<br>20wk: Control vs. SOD1, p < 0.0001; Control vs. SOD1+KDS12025 1mpk, p = 0.7111; Control vs. SOD1+KDS12025 10mpk, p = 0.3060<br>21wk: Control vs. SOD1, p < 0.0001; Control vs. SOD1+KDS12025 1mpk, p = 0.2127; Control vs. SOD1+KDS12025 10mpk, p = 0.1706<br>22wk: Control vs. SOD1, p < 0.0001; Control vs. SOD1+KDS12025 1mpk, p < 0.0001; Control vs. SOD1+KDS12025 10mpk, p = 0.0002<br>23wk: Control vs. SOD1, p < 0.0001; Control vs. SOD1+KDS12025 1mpk, p < 0.0001; Control vs. SOD1+KDS12025 10mpk, p < 0.0001<br>24wk: Control vs. SOD1, p = 0.0002; Control vs. SOD1+KDS12025 1mpk, p < 0.0001; Control vs. SOD1+KDS12025 10mpk, p < 0.0001 |
| 3o | Log-rank (Mantel-Cox) test, p < 0.001<br>Control (n=4, median survival = undefined), SOD1 (n=7, median survival = 140), SOD1+KDS12025 1mpk (n=7, median survival = 168), SOD1+KDS12025 10mpk (n=7, median survival = 168) |
| 4b | Two-way ANOVA with Tukey's multiple comparison test, F (1,350) = 600.9, p < 0.001<br>CA1: Normal (1.0±0.1025, n=30), AD (0.1101±0.01655, n=30); Normal vs. AD, p < 0.001<br>CA2: Normal (1.0±0.0761, n=30), AD (0.1514±0.02163, n=30); Normal vs. AD, p < 0.001<br>CA3: Normal (1.0±0.09249, n=30), AD (0.0888±0.01051, n=30); Normal vs. AD, p < 0.001<br>CA4: Normal (1.0±0.09124, n=30), AD (0.1165±0.01455, n=30); Normal vs. AD, p < 0.001<br>DG: Normal (1.0±0.07573, n=30), AD (0.07965±0.01313, n=30); Normal vs. AD, p < 0.001<br>Ent Cx: Normal (1.0±0.08698, n=30), AD (0.1148±0.01699, n=30); Normal vs. AD, p < 0.001 |
| 4d | Two-way ANOVA with Tukey's multiple comparison test, F (2,378) = 37.72, p < 0.001<br>Acquisition: WT (18.68±8.178, n=6), APP (19.40±5.736, n=5), APP+KDS12025 (ad libitum) (21.17±3.843, n=6)<br>WT vs. APP, p > 0.99; WT vs. APP+KDS12025 (ad libitum), p > 0.99; APP vs. APP+KDS12025 (ad libitum), p > 0.99<br>Retrieval: WT (271.5±76.57, n=6), APP (47.64±16.04, n=5), APP+KDS12025 (ad libitum) (254.3±56.96, n=6)<br>WT vs. APP, p = 0.003; WT vs. APP+KDS12025 (ad libitum), p = 0.95; APP vs. APP+KDS12025 (ad libitum), p = 0.006 |
| 4e | WT (0.2560±0.07604, n=6), APP (-0.09560±0.05233, n=5), APP+KDS12025 (ad libitum) (0.2057±0.04843, n=6)<br>One-way ANOVA with Tukey's multiple comparisons test, F(2,14) = 8.997, p = 0.003<br>WT vs. APP, p = 0.004; WT vs. APP+KDS12025 (ad libitum), p = 0.82; APP vs. APP+KDS12025 (ad libitum), p = 0.01 |
| 4g | Two-way ANOVA with Tukey's multiple comparison test, F (2,378) = 37.72, p < 0.001<br>WT (0.8145±0.09924, n=11), APP (0.4109±0.1036, n=11), APP+KDS12025 (ad libitum) (0.7455±0.1216, n=11)<br>WT vs. APP, p < 0.001; WT vs. APP+KDS12025 (ad libitum), p = 0.16; APP vs. APP+KDS12025 (ad libitum), p < 0.001 |
| 4h | WT (0.96±0.04, n=6), APP (0.08±0.08, n=5), APP+KDS12025 (ad libitum) (0.733±0.1085, n=6)<br>One-way ANOVA with Tukey's multiple comparisons test, F(2,13) = 26.68, p < 0.001<br>WT vs. APP, p < 0.001; WT vs. APP+KDS12025 (ad libitum), p = 0.18; APP vs. APP+KDS12025 (ad libitum), p < 0.001 |
| 4j | WT (70.03±1.523, n=55), APP (62.3±1.219, n=65), APP+KDS12025 (ad libitum) (68.67±2.358, n=54)<br>One-way ANOVA with Tukey's multiple comparisons test, F(2,171) = 6.123, p = 0.003<br>WT vs. APP, p = 0.004; WT vs. APP+KDS12025 (ad libitum), p = 0.85; APP vs. APP+KDS12025 (ad libitum), p = 0.02 |
| 4k | WT (59.70±2.280, n=5), APP (114.3±9.536, n=6), APP+KDS12025 (ad libitum) (64.48±6.979, n=6)<br>One-way ANOVA with Tukey's multiple comparisons test, F(2,14) = 17.24, p < 0.001<br>WT vs. APP, p < 0.001; WT vs. APP+KDS12025 (ad libitum), p = 0.89; APP vs. APP+KDS12025 (ad libitum), p < 0.001 |
| 4m | Two-way ANOVA with Tukey's multiple comparison test, F (1,42) = 37.70, p < 0.001<br>Sc-sh: Vehicle (100.0±0.9850, n=8), Aβ42 (125.7±2.521, n=8), Aβ42+KDS12025 (105.8±2.173, n=8)<br>Vehicle vs. Aβ42, p < 0.001; Vehicle vs. Aβ42+KDS12025, p = 0.40; APP vs. APP+KDS12025, p < 0.001<br>Hbβ-Sh: Vehicle (100.0±2.506, n=8), Aβ42 (144.7±3.744, n=8), Aβ42+KDS12025 (134.1±5.185, n=8)<br>Vehicle vs. Aβ42, p < 0.001; Vehicle vs. Aβ42+KDS12025, p < 0.001; APP vs. APP+KDS12025, p = 0.05 |
| 4o | Two-way ANOVA with Tukey's multiple comparison test, F (3,310) = 8.024, p < 0.001<br>WT (4.037±0.7779, n=9), APP (4.614±0.8108, n=12), APP+shHbβ+KDS12025 (5.314±0.9626, n=11), APP+ScRNA+KDS12025 (4.984±0.7027, n=8)<br>WT vs. APP, p = 0.01; WT vs. APP+shHbβ+KDS12025, p < 0.001; WT vs. APP+ScRNA+KDS12025, p > 0.99; APP vs. APP+shHbβ+KDS12025, p = 0.64; APP vs. APP+ScRNA+KDS12025, p = 0.03; APP+shHbβ+KDS12025 vs. APP+ScRNA+KDS12025, p = 0.001 |
| 4q | WT (11.25±0.6627, n=47), APP (8.798±0.5302, n=54), APP+shHbβ+KDS12025 (6.275±0.4473, n=83), APP+ScRNA+KDS12025 (10.10±0.4531, n=96)<br>One-way ANOVA with Tukey's multiple comparisons test, F(3,276) = 17.89, p < 0.001<br>WT vs. APP, p = 0.02; WT vs. APP+shHbβ+KDS12025, p < 0.001; WT vs. APP+ScRNA+KDS12025, p = 0.42; APP vs. APP+shHbβ+KDS12025, p = 0.004; APP vs. APP+ScRNA+KDS12025, p = 0.28; APP+shHbβ+KDS12025 vs. APP+ScRNA+KDS12025, p < 0.001 |
| 4r | WT (117.1±19.54, n=21), APP (203.7±22.90, n=18), APP+shHbβ+KDS12025 (234.3±16.93, n=26), APP+ScRNA+KDS12025 (142.4±8.715, n=35)<br>One-way ANOVA with Tukey's multiple comparisons test, F(3,99) = 61.21, p < 0.001<br>WT vs. APP, p = 0.005; WT vs. APP+shHbβ+KDS12025, p < 0.001; WT vs. APP+ScRNA+KDS12025, p = 0.65; APP vs. APP+shHbβ+KDS12025, p = 0.59; APP vs. APP+ScRNA+KDS12025, p = 0.04; APP+shHbβ+KDS12025 vs. APP+ScRNA+KDS12025, p < 0.001 |
| 4s | WT (117.8±2.617, n=21), APP (156.1±2.347, n=20), APP+shHbβ+KDS12025 (146.8±2.262, n=27), APP+ScRNA+KDS12025 (124.3±1.773, n=35)<br>One-way ANOVA with Tukey's multiple comparisons test, F(3,99) = 61.21, p < 0.001<br>WT vs. APP, p < 0.001; WT vs. APP+shHbβ+KDS12025, p < 0.001; WT vs. APP+ScRNA+KDS12025, p = 0.16; APP vs. APP+shHbβ+KDS12025, p = 0.03; APP vs. APP+ScRNA+KDS12025, p < 0.001; APP+shHbβ+KDS12025 vs. APP+ScRNA+KDS12025, p < 0.001 |
| S6d | CAT: Vehicle (93.8±11.09, n=8), HTPEB (84.5±9.957, n=8), KDS12025 (92.0±6.622, n=8)<br>One-way ANOVA with Tukey's multiple comparisons test, F(2,21) = 0.2723, p = 0.76<br>Vehicle vs. HTPEB, p = 0.77; Vehicle vs. KDS12025, p > 0.99; HTPEB vs. KDS12025, p = 0.84<br>HRP: Vehicle (7.25±0.479, n=4), HTPEB (9.0±1.0, n=4), KDS12025 (7.3±0.947, n=4)<br>One-way ANOVA with Tukey's multiple comparisons test, F(2,9) = 1.441, p = 0.29<br>Vehicle vs. HTPEB, p = 0.35; Vehicle vs. KDS12025, p > 0.99; HTPEB vs. KDS12025, p = 0.35<br>Hb: Vehicle (13.0±1.643, n=5), HTPEB (12.5±3.969, n=4), KDS12025 (13.5±2.533, n=4)<br>One-way ANOVA with Tukey's multiple comparisons test, F(2,10) = 0.03125, p = 0.96<br>Vehicle vs. HTPEB, p > 0.99; Vehicle vs. KDS12025, p > 0.99; HTPEB vs. KDS12025, p = 0.97 |
| S7b | Two-way ANOVA with Sidak's multiple comparison test, F(6,24) = 0.2512, p = 0.95<br>Full day: CTRL (18.99±0.3193, n=4), KDS12025 (10mpk) (19.14±0.6082, n=4), KDS12025 (1mpk) (21.26±0.07088, n=4), KDS12025 (0.1mpk) (22.21±1.751, n=4)<br>CTRL vs. KDS12025 (10mpk), p > 0.99; CTRL vs. KDS12025 (1mpk), p = 0.83; CTRL vs. KDS12025 (0.1mpk), p = 0.31; KDS12025 (10mpk) vs. KDS12025 (1mpk), p = 0.89; KDS12025 (10mpk) vs. KDS12025 (0.1mpk), p = 0.38; KDS12025 (1mpk) vs. KDS12025 (0.1mpk), p > 0.99<br>Dark: CTRL (20.68±0.4930, n=4), KDS12025 (10mpk) (20.73±0.9735, n=4), KDS12025 (1mpk) (21.29±0.7258, n=4), KDS12025 (0.1mpk) (22.37±1.215, n=4)<br>CTRL vs. KDS12025 (10mpk), p > 0.99; CTRL vs. KDS12025 (1mpk), p > 0.99; CTRL vs. KDS12025 (0.1mpk), p = 0.98; KDS12025 (10mpk) vs. KDS12025 (1mpk), p > 0.99; KDS12025 (10mpk) vs. KDS12025 (0.1mpk), p = 0.99; KDS12025 (1mpk) vs. KDS12025 (0.1mpk), p > 0.99<br>Light: CTRL (18.81±0.6740, n=4), KDS12025 (10mpk) (18.91±1.057, n=4), KDS12025 (1mpk) (19.95±0.3562, n=4), KDS12025 (0.1mpk) (20.90±1.556, n=4)<br>CTRL vs. KDS12025 (10mpk), p > 0.99; CTRL vs. KDS12025 (1mpk), p > 0.99; CTRL vs. KDS12025 (0.1mpk), p = 0.90; KDS12025 (10mpk) vs. KDS12025 (1mpk), p > 0.99; KDS12025 (10mpk) vs. KDS12025 (0.1mpk), p = 0.93; KDS12025 (1mpk) vs. KDS12025 (0.1mpk), p > 0.99 |
| S7c | Two-way ANOVA with Sidak's multiple comparison test, F(6,24) = 0.2059, p = 0.97<br>Full day: CTRL (0.1391±0.01165, n=4), KDS12025 (10mpk) (0.1288±0.01088, n=4), KDS12025 (1mpk) (0.1669±0.03508, n=4), KDS12025 (0.1mpk) (0.1260±0.01213, n=4)<br>CTRL vs. KDS12025 (10mpk), p > 0.99; CTRL vs. KDS12025 (1mpk), p > 0.99; CTRL vs. KDS12025 (0.1mpk), p > 0.99; KDS12025 (10mpk) vs. KDS12025 (1mpk), p = 0.93; KDS12025 (10mpk) vs. KDS12025 (0.1mpk), p > 0.99; KDS12025 (1mpk) vs. KDS12025 (0.1mpk), p = 0.96<br>Dark: CTRL (0.1680±0.01588, n=4), KDS12025 (10mpk) (0.1572±0.01881, n=4), KDS12025 (1mpk) (0.1761±0.01642, n=4), KDS12025 (0.1mpk) (0.1543±0.01806, n=4)<br>CTRL vs. KDS12025 (10mpk), p > 0.99; CTRL vs. KDS12025 (1mpk), p > 0.99; CTRL vs. KDS12025 (0.1mpk), p > 0.99; KDS12025 (10mpk) vs. KDS12025 (1mpk), p > 0.99; KDS12025 (10mpk) vs. KDS12025 (0.1mpk), p > 0.99; KDS12025 (1mpk) vs. KDS12025 (0.1mpk), p > 0.99<br>Light: CTRL (0.1056±0.01805, n=4), KDS12025 (10mpk) (0.09464±0.0095, n=4), KDS12025 (1mpk) (0.1032±0.0093, n=4), KDS12025 (0.1mpk) (0.07281±0.0039, n=4)<br>CTRL vs. KDS12025 (10mpk), p > 0.99; CTRL vs. KDS12025 (1mpk), p > 0.99; CTRL vs. KDS12025 (0.1mpk), p = 0.98; KDS12025 (10mpk) vs. KDS12025 (1mpk), p > 0.99; KDS12025 (10mpk) vs. KDS12025 (0.1mpk), p > 0.99; KDS12025 (1mpk) vs. KDS12025 (0.1mpk), p > 0.99 |
| S7d | Two-way ANOVA with Sidak's multiple comparison test, F(6,16) = 0.3547, p = 0.90<br>Full day: CTRL (0.07±0.01165, n=4), KDS12025 (10mpk) (0.07760±0.01206, n=4), KDS12025 (1mpk) (0.09027±0.02488, n=4), KDS12025 (0.1mpk) (0.1081±0.0057, n=4)<br>CTRL vs. KDS12025 (10mpk), p > 0.99; CTRL vs. KDS12025 (1mpk), p > 0.99; CTRL vs. KDS12025 (0.1mpk), p > 0.99; KDS12025 (10mpk) vs. KDS12025 (1mpk), p > 0.99; KDS12025 (10mpk) vs. KDS12025 (0.1mpk), p > 0.99; KDS12025 (1mpk) vs. KDS12025 (0.1mpk), p > 0.99<br>Dark: CTRL (0.08623±0.02568, n=4), KDS12025 (10mpk) (0.1041±0.03944, n=4), KDS12025 (1mpk) (0.1220±0.02080, n=4), KDS12025 (0.1mpk) (0.1272±0.009, n=4)<br>CTRL vs. KDS12025 (10mpk), p > 0.99; CTRL vs. KDS12025 (1mpk), p > 0.99; CTRL vs. KDS12025 (0.1mpk), p = 0.99; KDS12025 (10mpk) vs. KDS12025 (1mpk), p > 0.99; KDS12025 (10mpk) vs. KDS12025 (0.1mpk), p > 0.99; KDS12025 (1mpk) vs. KDS12025 (0.1mpk), p > 0.99<br>Light: CTRL (0.05996±0.01772, n=4), KDS12025 (10mpk) (0.05856±0.0075, n=4), KDS12025 (1mpk) (0.07906±0.01497, n=4), KDS12025 (0.1mpk) (0.06931±0.00048, n=4)<br>CTRL vs. KDS12025 (10mpk), p > 0.99; CTRL vs. KDS12025 (1mpk), p > 0.99; CTRL vs. KDS12025 (0.1mpk), p > 0.99; KDS12025 (10mpk) vs. KDS12025 (1mpk), p > 0.99; KDS12025 (10mpk) vs. KDS12025 (0.1mpk), p > 0.99; KDS12025 (1mpk) vs. KDS12025 (0.1mpk), p > 0.99 |
| S8b | Control (100.0±4.136, n=6), Aβ 0μM (89.89±4.975, n=3), Aβ 1μM (101.9±6.217, n=6), Aβ 5μM (137.3±8.218, n=6), Aβ 10μM (155.2±9.442, n=6)<br>One-way ANOVA with Tukey's multiple comparisons test, F(4,22) = 13.74, p < 0.0001<br>Control vs. Aβ 0μM, p = 0.9182; Control vs. Aβ 1μM, p = 0.9997; Control vs. Aβ 5μM, p = 0.0088; Control vs. Aβ 10μM, p = 0.0001; Aβ 0μM vs. Aβ 1μM, p = 0.8587; Aβ 0μM vs. Aβ 5μM, p = 0.0064; Aβ 0μM vs. Aβ 10μM, p = 0.0002; Aβ 1μM vs. Aβ 5μM, p = 0.0137; Aβ 1μM vs. Aβ 10μM, p = 0.0002; Aβ 5μM vs. Aβ 10μM, p = 0.3971 |
| S8c | Control (100.0±3.409, n=12), Aβ (5μM) (138.5±4.632, n=10), Aβ+KDS12008 (10μM) (96.78±7.989, n=10), Aβ+KDS12017 (10μM) (109.6±6.466, n=10), Aβ+KDS12025 (10μM) (110.3±5.804, n=11), Aβ+HTPEB (10μM) (112.0±8.321, n=8), Aβ+Sodium pyruvate (1mM) (149.1±6.194, n=5)<br>One-way ANOVA with Tukey's multiple comparisons test, F(6,59) = 8.225, p < 0.0001<br>CTRL vs. Aβ (5μM), p = 0.0003; CTRL vs. Aβ+KDS12008 (10μM), p = 0.99976; CTRL vs. Aβ+KDS12017 (10μM), p = 0.8993; CTRL vs. Aβ+KDS12025 (10μM), p = 0.8492; Aβ+HTPEB (10μM), p = 0.8079; Aβ+Sodium pyruvate (1mM), p = 0.0002; Aβ (5μM) vs. Aβ+KDS12008 (10μM), p = 0.0002; Aβ (5μM) vs. Aβ+KDS12017 (10μM), p = 0.0198; Aβ (5μM) vs. Aβ+KDS12025 (10μM), p = 0.0202; Aβ (5μM) vs. Aβ+HTPEB (10μM), p = 0.0672; Aβ (5μM) vs. Aβ+Sodium pyruvate (1mM), p = 0.9471; Aβ+KDS12008 (10μM) vs. Aβ+KDS12017 (10μM), p = 0.7391; Aβ+KDS12008 (10μM) vs. Aβ+KDS12025 (10μM), p = 0.6638; Aβ+KDS12008 (10μM) vs. Aβ+HTPEB (10μM), p = 0.6257; Aβ+KDS12008 (10μM) vs. Aβ+Sodium pyruvate (1mM), p < 0.0001; Aβ+KDS12017 (10μM) vs. Aβ+KDS12025 (10μM), p > 0.9999; Aβ+KDS12017 (10μM) vs. Aβ+HTPEB (10μM), p > 0.9999; Aβ+KDS12017 (10μM) vs. Aβ+Sodium pyruvate (1mM), p = 0.0061; Aβ+KDS12025 (10μM) vs. Aβ+HTPEB (10μM), p = 0.0064; Aβ+KDS12025 (10μM) vs. Aβ+Sodium pyruvate (10μM), p = 0.0184 |
| S8e | Control (100.0±3.367, n=4), Aβ 0μM (109.8±5.776, n=4), Aβ 1μM (109.4±4.271, n=4), Aβ 5μM (140.5±4.407, n=4), Aβ 10μM (208.1±3.662, n=4)<br>One-way ANOVA with Tukey's multiple comparisons test, F(4,15) = 102.9, p < 0.0001<br>Control vs. Aβ 0μM, p = 0.5290; Control vs. Aβ 1μM, p = 0.5641; Control vs. Aβ 5μM, p < 0.0001; Control vs. Aβ 10μM, p < 0.0001; Aβ 0μM vs. Aβ 1μM, p > 0.9999; Aβ 0μM vs. Aβ 5μM, p = 0.0014; Aβ 0μM vs. Aβ 10μM, p < 0.0001; Aβ 1μM vs. Aβ 5μM, p = 0.0012; Aβ 1μM vs. Aβ 10μM, p < 0.0001; Aβ 5μM vs. Aβ 10μM, p < 0.0001 |
| S8i | Vehicle (91.88±9.525, n=12), Aβ (203.1±11.93, n=11), Aβ+KDS12025 (111.8±4.968, n=12), Aβ+Sodium Pyruvate (137.5±9.751, n=10)<br>One-way ANOVA with Tukey's multiple comparisons test, F(3,41) = 27.64, p < 0.001<br>Vehicle vs. Aβ, p < 0.001; Vehicle vs. Aβ+KDS12025, p = 0.40; Vehicle vs. Aβ+Sodium Pyruvate, p = 0.007; Aβ vs. Aβ+KDS12025, p < 0.001; Aβ vs. Aβ+Sodium Pyruvate, p < 0.001; Aβ+KDS12025 vs. Aβ+Sodium Pyruvate, p = 0.23 |
| S8j | Vehicle (104.8±6.634, n=10), Putrescine (163.6±15.84, n=16), Put.+KDS12025 (96.12±6.634, n=28), Put.+Sodium Pyruvate (189.0±23.55, n=10)<br>One-way ANOVA with Tukey's multiple comparisons test, F(3,60) = 13.61, p < 0.001<br>Vehicle vs. Putrescine, p = 0.02; Vehicle vs. Put.+KDS12025, p = 0.96; Vehicle vs. Put.+Sodium Pyruvate, p = 0.001; Putrescine vs. Put.+KDS12025, p < 0.001; Putrescine vs. Put.+Sodium Pyruvate, p = 0.55; Put.+KDS12025 vs. Put.+Sodium Pyruvate, p < 0.001 |
| S9b | Two-way ANOVA with Sidak's multiple comparison test, F(4,58) = 2.950, p = 0.0275<br>Acquisition: WT (45.24±14.0, n=9), APP/PS1 (56.45±20.33, n=6), APP/PS1+KDS12008 (44.42±13.85, n=6), APP/PS1+KDS12017 (71.53±23.24, n=4), APP/PS1+KDS12025 (29.03±6.431, n=9)<br>WT vs. APP/PS1, p > 0.9999; WT vs. APP/PS1+KDS12008, p > 0.9999; WT vs. APP/PS1+KDS12017, p > 0.9999; WT vs. APP/PS1+KDS12025, p > 0.9999; APP/PS1 vs. APP/PS1+KDS12008, p > 0.9999; APP/PS1 vs. APP/PS1+KDS12017, p > 0.9999; APP/PS1 vs. APP/PS1+KDS12025, p > 0.9999; APP/PS1+KDS12008 vs. APP/PS1+KDS12017, p > 0.9999; APP/PS1+KDS12008 vs. APP/PS1+KDS12025, p > 0.9999; APP/PS1+KDS12017 vs. APP/PS1+KDS12025, p > 0.9999<br>Retrieval: WT (414.1±63.65, n=9), APP/PS1 (59.87±22.89, n=6), APP/PS1+KDS12008 (233.6±102.3, n=6), APP/PS1+KDS12017 (305.6±137.5, n=4), APP/PS1+KDS12025 (291.0±68.80, n=9)<br>WT vs. APP/PS1, p = 0.0002; WT vs. APP/PS1+KDS12008, p = 0.2005; WT vs. APP/PS1+KDS12017, p = 0.9166; WT vs. APP/PS1+KDS12025, p = 0.5572; APP/PS1 vs. APP/PS1+KDS12008, p = 0.3581; APP/PS1 vs. APP/PS1+KDS12017, p = 0.1085; APP/PS1 vs. APP/PS1+KDS12025, p = 0.0379; APP/PS1+KDS12008 vs. APP/PS1+KDS12017, p = 0.9973; APP/PS1+KDS12008 vs. APP/PS1+KDS12025, p = 0.9978; APP/PS1+KDS12017 vs. APP/PS1+KDS12025, p > 0.9999 |
| S9e | WT (1557±63.18, n=19), APP/PS1 (2513±115.0, n=41), APP/PS1+KDS12008 (2063±93.76, n=32), APP/PS1+KDS12017 (1711±145.9, n=23), APP/PS1+KDS12025 (1190±87.29, n=19)<br>One-way ANOVA with Tukey's multiple comparisons test, F(4,129) = 20.43, p < 0.001<br>WT vs. APP/PS1, p < 0.001; WT vs. APP/PS1+KDS12008, p = 0.03; WT vs. APP/PS1+KDS12017, p = 0.92; WT vs. APP/PS1+KDS12025, p = 0.32; APP/PS1 vs. APP/PS1+KDS12008, p = 0.01; APP/PS1 vs. APP/PS1+KDS12017, p < 0.001; APP/PS1 vs. APP/PS1+KDS12025, p < 0.001; APP/PS1+KDS12008 vs. APP/PS1+KDS12017, p = 0.19; APP/PS1+KDS12008 vs. APP/PS1+KDS12025, p < 0.001; APP/PS1+KDS12017 vs. APP/PS1+KDS12025, p = 0.04 |
| S9f | WT (1213±22.64, n=20), APP/PS1 (1512±43.96, n=43), APP/PS1+KDS12008 (1562±64.27, n=32), APP/PS1+KDS12017 (1556±53.65, n=21), APP/PS1+KDS12025 (1343±38.44, n=36)<br>One-way ANOVA with Tukey's multiple comparisons test, F(4,147) = 7.937, p < 0.001<br>WT vs. APP/PS1, p < 0.001; WT vs. APP/PS1+KDS12008, p < 0.001; WT vs. APP/PS1+KDS12017, p < 0.001; WT vs. APP/PS1+KDS12025, p = 0.43; APP/PS1 vs. APP/PS1+KDS12008, p = 0.94; APP/PS1 vs. APP/PS1+KDS12017, p = 0.97; APP/PS1 vs. APP/PS1+KDS12025, p = 0.05; APP/PS1+KDS12008 vs. APP/PS1+KDS12017, p > 0.99; APP/PS1+KDS12008 vs. APP/PS1+KDS12025, p = 0.01; APP/PS1+KDS12017 vs. APP/PS1+KDS12025, p = 0.04 |
| S9h | Two-way ANOVA with Tukey's multiple comparison test, F(3,66) = 13.58, p < 0.001<br>Acquisition: WT+Saline (40.67±6.90, n=11), APP/PS1+Saline (22.17±3.617, n=7), APP/PS1+KDS12025 (3mpk) (36.47±13.39, n=10), APP/PS1+KDS12025 (10mpk) (66.00±15.30, n=9)<br>WT+Saline vs. APP/PS1+Saline, p = 0.99; WT+Saline vs. APP/PS1+KDS12025 (3mpk), p > 0.99; WT+Saline vs. APP/PS1+KDS12025 (10mpk), p = 0.96; APP/PS1+Saline vs. APP/PS1+KDS12025 (3mpk), p > 0.99; APP/PS1+Saline vs. APP/PS1+KDS12025 (10mpk), p = 0.86; APP/PS1+KDS12025 (3mpk) vs. APP/PS1+KDS12025 (10mpk), p = 0.94<br>Retrieval: WT+Saline (556.0±31.47, n=11), APP/PS1+Saline (83.69±15.91, n=7), APP/PS1+KDS12025 (3mpk) (341.7±71.16, n=10), APP/PS1+KDS12025 (10mpk) (508.8±49.93, n=9)<br>WT+Saline vs. APP/PS1+Saline, p < 0.001; WT+Saline vs. APP/PS1+KDS12025 (3mpk), p < 0.001; WT+Saline vs. APP/PS1+KDS12025 (10mpk), p = 0.77; APP/PS1+Saline vs. APP/PS1+KDS12025 (3mpk), p < 0.001; APP/PS1+Saline vs. APP/PS1+KDS12025 (10mpk), p < 0.001; APP/PS1+KDS12025 (3mpk) vs. APP/PS1+KDS12025 (10mpk), p = 0.008 |
| S9j | WT (54.04±4.850, n=6), APP/PS1 (84.53±6.827, n=11), APP/PS1+KDS (3mpk) (59.33±6.459, n=10), APP/PS1+KDS (10mpk) (56.71±4.775, n=9)<br>One-way ANOVA with Tukey's multiple comparisons test, F(3,39) = 8.841, p < 0.001<br>WT vs. APP/PS1, p = 0.02; WT vs. APP/PS1+KDS (3mpk), p = 0.95; WT vs. APP/PS1+KDS (10mpk), p > 0.99; APP/PS1 vs. APP/PS1+KDS (3mpk), p = 0.02; APP/PS1 vs. APP/PS1+KDS (10mpk), p = 0.01; APP/PS1+KDS (3mpk) vs. APP/PS1+KDS (10mpk), p = 0.99 |
| S9l | WT (4.001±0.4549, n=15), APP/PS1 (11.24±1.731, n=16), APP/PS1+KDS (3mpk) (4.160±0.8926, n=7), APP/PS1+KDS (10mpk) (3.113±0.7279, n=5)<br>One-way ANOVA with Tukey's multiple comparisons test, F(3,39) = 8.841, p < 0.001<br>WT vs. APP/PS1, p < 0.001; WT vs. APP/PS1+KDS (3mpk), p > 0.99; WT vs. APP/PS1+KDS (10mpk), p = 0.98; APP/PS1 vs. APP/PS1+KDS (3mpk), p = 0.007; APP/PS1 vs. APP/PS1+KDS (10mpk), p = 0.006; APP/PS1+KDS (3mpk) vs. APP/PS1+KDS (10mpk), p = 0.98 |
| S9m | WT (34.57±1.908, n=10), APP/PS1 (41.39±4.835, n=16), APP/PS1+KDS (3mpk) (36.09±4.601, n=7), APP/PS1+KDS (10mpk) (26.02±1.734, n=5)<br>One-way ANOVA with Tukey's multiple comparisons test, F(3,34) = 1.596, p = 0.21<br>WT vs. APP/PS1, p = 0.64; WT vs. APP/PS1+KDS (3mpk), p > 0.99; WT vs. APP/PS1+KDS (10mpk), p = 0.60; APP/PS1 vs. APP/PS1+KDS (3mpk), p = 0.84; APP/PS1 vs. APP/PS1+KDS (10mpk), p = 0.17; APP/PS1+KDS (3mpk) vs. APP/PS1+KDS (10mpk), p = 0.63 |
| S9n | WT (1.983±0.4121, n=11), APP/PS1 (2.428±0.3485, n=15), APP/PS1+KDS (3mpk) (1.355±0.2694, n=7), APP/PS1+KDS (10mpk) (1.177±0.5985, n=5)<br>One-way ANOVA with Tukey's multiple comparisons test, F(3,34) = 1.596, p = 0.21<br>WT vs. APP/PS1, p = 0.81; WT vs. APP/PS1+KDS (3mpk), p = 0.73; WT vs. APP/PS1+KDS (10mpk), p = 0.64; APP/PS1 vs. APP/PS1+KDS (3mpk), p = 0.27; APP/PS1 vs. APP/PS1+KDS (10mpk), p = 0.24; APP/PS1+KDS (3mpk) vs. APP/PS1+KDS (10mpk), p > 0.99 |
| S10a | Total distance: WT+GFP (2308±148.9, n=9), APP+DTR (2339±561.4, n=6), APP+DTR+KDS (2372±293.8, n=11)<br>One-way ANOVA with Tukey's multiple comparisons test, F(2,23) = 0.01147, p = 0.99<br>WT+GFP vs. APP+DTR, p > 0.99; WT+GFP vs. APP+DTR+KDS, p = 0.99; APP+DTR vs. APP+DTR+KDS, p > 0.99<br>Center frequency: WT+GFP (38.67±7.446, n=9), APP+DTR (37.25±10.31, n=6), APP+DTR+KDS (34.80±7.729, n=11)<br>One-way ANOVA with Tukey's multiple comparisons test, F(2,12) = 0.05969, p = 0.94<br>WT+GFP vs. APP+DTR, p > 0.99; WT+GFP vs. APP+DTR+KDS, p = 0.94; APP+DTR vs. APP+DTR+KDS, p = 0.98<br>Center duration: WT+GFP (80.41±22.51, n=9), APP+DTR (53.97±11.90, n=6), APP+DTR+KDS (75.75±13.63, n=11)<br>One-way ANOVA with Tukey's multiple comparisons test, F(2,22) = 0.5221, p = 0.60<br>WT+GFP vs. APP+DTR, p = 0.60; WT+GFP vs. APP+DTR+KDS, p = 0.98; APP+DTR vs. APP+DTR+KDS, p = 0.69 |
| S10b | Total distance: WT (2371±460.1, n=6), APP (2336±373.4, n=4), APP+KDS12025 (ad libitum) (2987±763.8, n=4)<br>One-way ANOVA with Tukey's multiple comparisons test, F(2,11) = 0.4168, p = 0.67<br>WT vs. APP, p > 0.99; WT vs. APP+KDS12025 (ad libitum), p = 0.70; APP vs. APP+KDS12025 (ad libitum), p = 0.72<br>Center frequency: WT (34.17±5.224, n=6), APP (34.25±7.598, n=4), APP+KDS12025 (ad libitum) (30.75±8.025, n=4)<br>One-way ANOVA with Tukey's multiple comparisons test, F(2,11) = 0.08192, p = 0.92<br>WT vs. APP, p > 0.99; WT vs. APP+KDS12025 (ad libitum), p = 0.93; APP vs. APP+KDS12025 (ad libitum), p = 0.94<br>Center duration: WT (69.22±14.49, n=6), APP (57.80±9.964, n=4), APP+KDS12025 (ad libitum) (65.64±21.86, n=4)<br>One-way ANOVA with Tukey's multiple comparisons test, F(2,11) = 0.1314, p = 0.88<br>WT vs. APP, p = 0.87; WT vs. APP+KDS12025 (ad libitum), p = 0.99; APP vs. APP+KDS12025 (ad libitum), p = 0.95 |
| S10c | WT (0.2256±0.02750, n=3), GiD (-0.02438±0.07989, n=3), GiD+KDS12025 (0.1mpk) (0.2255±0.02347, n=4)<br>One-way ANOVA with Tukey's multiple comparisons test, F(2,7) = 9.285, p = 0.01<br>WT vs. GiD, p = 0.02; WT vs. GiD+KDS12025 (0.1mpk), p > 0.99; GiD vs. GiD+KDS12025 (0.1mpk), p = 0.01 |
| S10d | Total distance: WT (2043±158.7, n=6), APP (2159±465.6, n=3), APP+shHbβ+KDS12025 (2873±187.1, n=7), APP+ScRNA+KDS12025 (2220±326.5, n=6)<br>One-way ANOVA with Tukey's multiple comparisons test, F(3,18) = 2.390, p = 0.10<br>WT vs. APP, p > 0.99; WT vs. APP+shHbβ+KDS12025, p = 0.10; WT vs. APP+ScRNA+KDS12025, p = 0.96; APP vs. APP+shHbβ+KDS12025, p = 0.36; APP vs. APP+ScRNA+KDS12025, p > 0.99; APP+shHbβ+KDS12025 vs. APP+ScRNA+KDS12025, p = 0.26<br>Center frequency: WT (34.00±2.380, n=6), APP (29.33±8.192, n=3), APP+shHbβ+KDS12025 (36.43±5.237, n=7), APP+ScRNA+KDS12025 (31.33±7.065, n=6)<br>One-way ANOVA with Tukey's multiple comparisons test, F(3,18) = 0.2637, p = 0.85<br>WT vs. APP, p > 0.96; WT vs. APP+shHbβ+KDS12025, p = 0.99; WT vs. APP+ScRNA+KDS12025, p = 0.99; APP vs. APP+shHbβ+KDS12025, p = 0.87; APP vs. APP+ScRNA+KDS12025, p > 0.99; APP+shHbβ+KDS12025 vs. APP+ScRNA+KDS12025, p = 0.90<br>Center frequency: WT (84.14±7.605, n=6), APP (50.72±9.915, n=3), APP+shHbβ+KDS12025 (68.36±13.11, n=7), APP+ScRNA+KDS12025 (60.94±13.14, n=6)<br>One-way ANOVA with Tukey's multiple comparisons test, F(3,18) = 1.123, p = 0.37<br>WT vs. APP, p = 0.38; WT vs. APP+shHbβ+KDS12025, p = 0.76; WT vs. APP+ScRNA+KDS12025, p = 0.51; APP vs. APP+shHbβ+KDS12025, p = 0.81; APP vs. APP+ScRNA+KDS12025, p = 0.96; APP+shHbβ+KDS12025 vs. APP+ScRNA+KDS12025, p = 0.97 |
| S11d | WT (41.73±1.1951, n=19), APP (50.90±1.074, n=41), APP+KDS12025 (ad libitum) (44.45±1.015, n=28)<br>One-way ANOVA with Tukey's multiple comparisons test, F(2,85) = 14.24, p < 0.001<br>WT vs. APP, p < 0.001; WT vs. APP+KDS12025 (ad libitum), p = 0.38; APP vs. APP+KDS12025 (ad libitum), p < 0.001 |
| S11e | WT (1903±72.86, n=32), APP (2435±66.11, n=58), APP+KDS12025 (ad libitum) (1951±48.61, n=57)<br>One-way ANOVA with Tukey's multiple comparisons test, F(2,144) = 23.42, p < 0.001<br>WT vs. APP, p < 0.001; WT vs. APP+KDS12025 (ad libitum), p = 0.88; APP vs. APP+KDS12025 (ad libitum), p < 0.001 |
| S11f | WT (31.45±3.606, n=11), APP (63.19±5.017, n=16), APP+KDS12025 (ad libitum) (33.69±3.3, n=13)<br>One-way ANOVA with Tukey's multiple comparisons test, F(2,37) = 18.10, p < 0.001<br>WT vs. APP, p < 0.001; WT vs. APP+KDS12025 (ad libitum), p = 0.94; APP vs. APP+KDS12025 (ad libitum), p < 0.001 |
| S11g | WT (1.617±0.2177, n=11), APP (7.134±1.523, n=16), APP+KDS12025 (ad libitum) (2.808±0.6584, n=13)<br>One-way ANOVA with Tukey's multiple comparisons test, F(2,37) = 6.945, p = 0.003<br>WT vs. APP, p = 0.004; WT vs. APP+KDS12025 (ad libitum), p = 0.76; APP vs. APP+KDS12025 (ad libitum), p = 0.02 |
| S11h | WT (110.0±6.606, n=11), APP (135.6±5.625, n=16), APP+KDS12025 (ad libitum) (103.1±7.794, n=13)<br>One-way ANOVA with Tukey's multiple comparisons test, F(2,37) = 7.249, p = 0.002<br>WT vs. APP, p = 0.03; WT vs. APP+KDS12025 (ad libitum), p = 0.77; APP vs. APP+KDS12025 (ad libitum), p = 0.003 |
| S11i | Two-way ANOVA with Tukey's multiple comparison test, F(2,378) = 37.72, p < 0.001<br>WT (2.028, n=12), APP (4.318, n=17), APP+KDS12025 (ad libitum) (2.392, n=12)<br>WT vs. APP, p < 0.001; WT vs. APP+KDS12025 (ad libitum), p = 0.52; APP vs. APP+KDS12025 (ad libitum), p < 0.001 |
| S12c | Control (89.59±3.31, n=35), A53T ipsi (186.7±1.985, n=57), A53T ipsi+KDS12025 (106.7±3.202, n=32), A53T contra (79.40±5.141, n=31), A53T contra+KDS12025 (82.69±4.999, n=28)<br>One-way ANOVA with Tukey's multiple comparisons test, F(4,178) = 214.4, p < 0.001<br>Control vs. A53T ipsi, p < 0.001; Control vs. A53T ipsi+KDS12025, p = 0.009; Control vs. A53T contra, p = 0.29; Control vs. A53T contra+KDS12025, p = 0.70; A53T ipsi vs. A53T ipsi+KDS12025, p < 0.001; A53T ipsi vs. A53T contra, p < 0.001; A53T ipsi vs. A53T contra+KDS12025, p < 0.0001; A53T ipsi+KDS12025 vs. A53T contra, p < 0.001; A53T ipsi+KDS12025 vs. A53T contra+KDS12025, p < 0.001; A53T contra vs. A53T contra+KDS12025, p = 0.98 |
| S13b | Normal (0.0±0, n=6), CIA+Vehicle (11.08±1.274, n=6), CIA+KDS12025 0.1 MPK (9.5±1.133, n=6), CIA+KDS12025 1 MPK (6.583±0.7002, n=6)<br>One-way ANOVA with Tukey's multiple comparisons test, F(3,20) = 28.23, p < 0.001<br>Normal vs. CIA+Vehicle, p < 0.001; Normal vs. CIA+KDS12025 0.1 MPK, p < 0.001; Normal vs. CIA+KDS12025 1 MPK, p < 0.001; CIA+Vehicle vs. CIA+KDS12025 0.1 MPK, p = 0.62; CIA+Vehicle vs. CIA+KDS12025 1 MPK, p = 0.01; CIA+KDS12025 0.1 MPK vs. CIA+KDS12025 1 MPK, p = 0.15 |
| S13c | Normal (0.0±0, n=6), CIA+Vehicle (70.83±11.93, n=6), CIA+KDS12025 0.1 MPK (54.17±15.02, n=6), CIA+KDS12025 1 MPK (25.0±9.129, n=6)<br>One-way ANOVA with Tukey's multiple comparisons test, F(3,20) = 8.718, p < 0.001<br>Normal vs. CIA+Vehicle, p < 0.001; Normal vs. CIA+KDS12025 0.1 MPK, p = 0.009; Normal vs. CIA+KDS12025 1 MPK, p = 0.37; CIA+Vehicle vs. CIA+KDS12025 0.1 MPK, p = 0.69; CIA+Vehicle vs. CIA+KDS12025 1 MPK, p = 0.03; CIA+KDS12025 0.1 MPK vs. CIA+KDS12025 1 MPK, p = 0.24 |
| S13d | Normal (0.09±0.0, n=6), CIA+Vehicle (0.1533±0.01085, n=6), CIA+KDS12025 0.1 MPK (0.1383±0.0074, n=6), CIA+KDS12025 1 MPK (0.1183±0.0047, n=6)<br>One-way ANOVA with Tukey's multiple comparisons test, F(3,20) = 15.25, p < 0.001<br>Normal vs. CIA+Vehicle, p < 0.001; Normal vs. CIA+KDS12025 0.1 MPK, p < 0.001; Normal vs. CIA+KDS12025 1 MPK, p = 0.04; CIA+Vehicle vs. CIA+KDS12025 0.1 MPK, p = 0.45; CIA+Vehicle vs. CIA+KDS12025 1 MPK, p = 0.01; CIA+KDS12025 0.1 MPK vs. CIA+KDS12025 1 MPK, p = 0.22 |
| S13f | Normal (0.0±0, n=5), CIA+Vehicle (2.964±0.024, n=5), CIA+KDS12025 0.1 MPK (1.338±0.2985, n=5), CIA+KDS12025 1 MPK (0.7620±0.3263, n=5)<br>One-way ANOVA with Tukey's multiple comparisons test, $F(3,16) = 32.25$, $p < 0.001$<br>Normal vs. CIA+Vehicle, $p < 0.001$; Normal vs. CIA+KDS12025 0.1 MPK, $p = 0.003$; Normal vs. CIA+KDS12025 1 MPK, $p = 0.11$; CIA+Vehicle vs. CIA+KDS12025 0.1 MPK, $p < 0.001$; CIA+Vehicle vs. CIA+KDS12025 1 MPK, $p < 0.001$; CIA+KDS12025 0.1 MPK vs. CIA+KDS12025 1 MPK, $p = 0.29$ |
| S13g | Normal (0.0±0, n=5), CIA+Vehicle (3.0±0, n=5), CIA+KDS12025 0.1 MPK (1.5±0.2949, n=5), CIA+KDS12025 1 MPK (0.966±0.2183, n=5)<br>One-way ANOVA with Tukey's multiple comparisons test, $F(3,16) = 46.69$, $p < 0.001$<br>Normal vs. CIA+Vehicle, $p < 0.001$; Normal vs. CIA+KDS12025 0.1 MPK, $p < 0.001$; Normal vs. CIA+KDS12025 1 MPK, $p = 0.009$; CIA+Vehicle vs. CIA+KDS12025 0.1 MPK, $p < 0.001$; CIA+Vehicle vs. CIA+KDS12025 1 MPK, $p < 0.001$; CIA+KDS12025 0.1 MPK vs. CIA+KDS12025 1 MPK, $p = 0.21$ |
| S13h | Normal (0.0±0, n=5), CIA+Vehicle (2.664±0.1938, n=5), CIA+KDS12025 0.1 MPK (0.9±0.3929, n=5), CIA+KDS12025 1 MPK (0.338±0.2928, n=5)<br>One-way ANOVA with Tukey's multiple comparisons test, $F(3,16) = 20.24$, $p < 0.001$<br>Normal vs. CIA+Vehicle, $p < 0.001$; Normal vs. CIA+KDS12025 0.1 MPK, $p = 0.11$; Normal vs. CIA+KDS12025 1 MPK, $p = 0.80$; CIA+Vehicle vs. CIA+KDS12025 0.1 MPK, $p < 0.001$; CIA+Vehicle vs. CIA+KDS12025 1 MPK, $p < 0.001$; CIA+KDS12025 0.1 MPK vs. CIA+KDS12025 1 MPK, $p = 0.46$ |
| S13i | Normal (0.0±0, n=5), CIA+Vehicle (2.64±0.1974, n=5), CIA+KDS12025 0.1 MPK (0.852±0.3905, n=5), CIA+KDS12025 1 MPK (0.338±0.2928, n=5)<br>One-way ANOVA with Tukey's multiple comparisons test, $F(3,16) = 19.93$, $p < 0.001$<br>Normal vs. CIA+Vehicle, $p < 0.001$; Normal vs. CIA+KDS12025 0.1 MPK, $p = 0.14$; Normal vs. CIA+KDS12025 1 MPK, $p = 0.80$; CIA+Vehicle vs. CIA+KDS12025 0.1 MPK, $p < 0.001$; CIA+Vehicle vs. CIA+KDS12025 1 MPK, $p < 0.001$; CIA+KDS12025 0.1 MPK vs. CIA+KDS12025 1 MPK, $p = 0.53$ |
| S13k | Normal (0.0±0, n=5), CIA+Vehicle (3.0±0, n=5), CIA+KDS12025 0.1 MPK (1.504±0.3303, n=5), CIA+KDS12025 1 MPK (0.988±0.2547, n=5)<br>One-way ANOVA with Tukey's multiple comparisons test, $F(3,16) = 19.93$, $p < 0.001$<br>Normal vs. CIA+Vehicle, $p < 0.001$; Normal vs. CIA+KDS12025 0.1 MPK, $p = 0.14$; Normal vs. CIA+KDS12025 1 MPK, $p = 0.80$; CIA+Vehicle vs. CIA+KDS12025 0.1 MPK, $p < 0.001$; CIA+Vehicle vs. CIA+KDS12025 1 MPK, $p < 0.001$; CIA+KDS12025 0.1 MPK vs. CIA+KDS12025 1 MPK, $p = 0.53$ |
| S13l | Normal (0.0±0, n=5), CIA+Vehicle (2.976±0.024, n=5), CIA+KDS12025 0.1 MPK (1.352±0.3083, n=5), CIA+KDS12025 1 MPK (0.95±0.2009, n=5)<br>One-way ANOVA with Tukey's multiple comparisons test, $F(3,16) = 45.32$, $p < 0.001$<br>Normal vs. CIA+Vehicle, $p < 0.001$; Normal vs. CIA+KDS12025 0.1 MPK, $p < 0.001$; Normal vs. CIA+KDS12025 1 MPK, $p = 0.01$; CIA+Vehicle vs. CIA+KDS12025 0.1 MPK, $p < 0.001$; CIA+Vehicle vs. CIA+KDS12025 1 MPK, $p < 0.001$; CIA+KDS12025 0.1 MPK vs. CIA+KDS12025 1 MPK, $p = 0.44$ |
| S13m | Normal (0.0±0, n=5), CIA+Vehicle (2.188±0.1666, n=5), CIA+KDS12025 0.1 MPK (0.85±0.2018, n=5), CIA+KDS12025 1 MPK (0.652±0.1825, n=5)<br>One-way ANOVA with Tukey's multiple comparisons test, $F(3,16) = 30.86$, $p < 0.001$<br>Normal vs. CIA+Vehicle, $p < 0.001$; Normal vs. CIA+KDS12025 0.1 MPK, $p = 0.01$; Normal vs. CIA+KDS12025 1 MPK, $p = 0.06$; CIA+Vehicle vs. CIA+KDS12025 0.1 MPK, $p < 0.001$; CIA+Vehicle vs. CIA+KDS12025 1 MPK, $p < 0.001$; CIA+KDS12025 0.1 MPK vs. CIA+KDS12025 1 MPK, $p = 0.83$ |
| S13o | Control (0.2808±0.02283, n=5), LPS-2hrs (0.5301±0.04939, n=7), LPS-6hrs (1.204±0.05891, n=5), LPS-24hrs (0.2503±0.03503, n=8)<br>One-way ANOVA with Tukey's multiple comparisons test, $F(3,21) = 89.41$, $p < 0.001$<br>Control vs. LPS-2hrs, $p = 0.004$; Control vs. LPS-6hrs, $p < 0.001$; Control vs. LPS-24hrs, $p = 0.96$; LPS-2hrs vs. LPS-6hrs, $p < 0.001$; LPS-2hrs vs. LPS-24hrs, $p < 0.001$; LPS-6hrs vs. LPS-24hrs, $p < 0.001$ |
| S13p | Control (0.3325±0.02161, n=6), LPS-6hrs (1.295±0.05323, n=6), LPS-6hrs+KDS (0.7847±0.08078, n=6)<br>One-way ANOVA with Tukey's multiple comparisons test, $F(2,15) = 70.74$, $p < 0.001$<br>Control vs. LPS-6hrs, $p < 0.001$; Control vs. LPS-6hrs+KDS, $p < 0.001$; LPS-6hrs vs. LPS-6hrs+KDS, $p < 0.001$ |
| S14b | Unpaired t-test, one-tailed<br>Normal (1.0±0.3789, n=8) vs. AD (0.3084±0.02120, n=8), $p = 0.04$, $t = 1.822$, $df = 14$ |
| S14c | Unpaired t-test, two-tailed<br>WT (0.5166±0.08026, n=4) vs. APP/PS1 (0.2328±0.03508, n=4), $p = 0.02$, $t = 3.241$, $df = 6$ |
| S15c | Control (26.40±1.082, n=61), A53T ipsi (44.57±2.459, n=31), A53T ipsi+KDS12025 (34.98±2.564, n=20), A53T contra (45.24±4.064, n=19), A53T contra+KDS12025 (46.92±4.141, n=12)<br>One-way ANOVA with Tukey's multiple comparisons test, $F(4,138) = 18.62$, $p < 0.001$<br>Control vs. A53T ipsi, $p < 0.001$; Control vs. A53T ipsi+KDS12025, $p = 0.98$; Control vs. A53T contra, $p = 0.06$; Control vs. A53T contra+KDS12025, $p > 0.99$; A53T ipsi vs. A53T ipsi+KDS12025, $p < 0.001$; A53T ipsi vs. A53T contra, $p = 0.05$; A53T ipsi vs. A53T contra+KDS12025, $p < 0.0001$; A53T ipsi+KDS12025 vs. A53T contra, $p = 0.05$; A53T ipsi+KDS12025 vs. A53T contra+KDS12025, $p > 0.99$; A53T contra vs. A53T contra+KDS12025, $p = 0.07$ |
| S15d | Control (65.22±6.586, n=14), A53T ipsi (60.57±4.01, n=13), A53T ipsi+KDS12025 (55.65±3.378, n=28), A53T contra (60.30±3.44, n=39), A53T contra+KDS12025 (61.58±2.401, n=38)<br>One-way ANOVA with Tukey's multiple comparisons test, $F(4,127) = 0.8159$, $p = 0.58$<br>Control vs. A53T ipsi, $p = 0.97$; Control vs. A53T ipsi+KDS12025, $p = 0.53$; Control vs. A53T contra, $p = 0.92$; Control vs. A53T contra+KDS12025, $p = 0.97$; A53T ipsi vs. A53T ipsi+KDS12025, $p = 0.94$; A53T ipsi vs. A53T contra, $p > 0.99$; A53T ipsi vs. A53T contra+KDS12025, $p > 0.99$; A53T ipsi+KDS12025 vs. A53T contra, $p = 0.85$; A53T ipsi+KDS12025 vs. A53T contra+KDS12025, $p = 0.71$; A53T contra vs. A53T contra+KDS12025, $p > 0.99$ |
| S16e | WT (117.1±19.54, n=21), APP (203.7±22.90, n=18), APP+shHbβ+KDS12025 (234.3±16.93, n=26), APP+ScRNA+KDS12025 (142.4±8.715, n=35)<br>One-way ANOVA with Tukey's multiple comparisons test, $F(3,99) = 61.21$, $p < 0.001$<br>WT vs. APP, $p = 0.005$; WT vs. APP+shHbβ+KDS12025, $p < 0.001$; WT vs. APP+ScRNA+KDS12025, $p = 0.65$; APP vs. APP+shHbβ+KDS12025, $p = 0.59$; APP vs. APP+ScRNA+KDS12025, $p = 0.04$; APP+shHbβ+KDS12025 vs. APP+ScRNA+KDS12025, $p < 0.001$ |
| S16f | WT (72.00±7.810, n=10), APP (113.1±6.305, n=14), APP+shHbβ+KDS12025 (100.4±6.075, n=28), APP+ScRNA+KDS12025 (54.48±2.319, n=23)<br>One-way ANOVA with Tukey's multiple comparisons test, $F(3,61) = 28.97$, $p < 0.001$<br>WT vs. APP, $p < 0.001$; WT vs. APP+shHbβ+KDS12025, $p = 0.006$; WT vs. APP+ScRNA+KDS12025, $p = 0.13$; APP vs. APP+shHbβ+KDS12025, $p = 0.33$; APP vs. APP+ScRNA+KDS12025, $p < 0.001$; APP+shHbβ+KDS12025 vs. APP+ScRNA+KDS12025, $p < 0.001$ |
| S16g | Two-way ANOVA with Tukey's multiple comparison test, $F(3,310) = 8.024$, $p < 0.001$<br>WT ($4.037 \pm 0.7779$, $n=9$), APP ($4.614 \pm 0.8108$, $n=12$),<br>APP+shHbβ+KDS12025 ($5.314 \pm 0.9626$, $n=11$), APP+ScRNA+KDS12025 ($4.984 \pm 0.7027$, $n=8$)<br>WT vs. APP, $p = 0.01$; WT vs. APP+shHbβ+KDS12025, $p < 0.001$; WT vs. APP+ScRNA+KDS12025, $p > 0.99$; APP vs. APP+shHbβ+KDS12025, $p = 0.64$; APP vs. APP+ScRNA+KDS12025, $p = 0.03$; APP+shHbβ+KDS12025 vs. APP+ScRNA+KDS12025, $p = 0.001$ |
| S16i | Unaired t-test, two tailed<br>Ctrl ($1.191 \pm 0.3225$, $n=6$), D1 ($4.588 \pm 0.9107$, $n=6$), $p = 0.006$; $t = 3.517$, $df = 10$ |
| S16j | Ctrl ($1.0 \pm 0.1$, $n=3$), H<sub>2</sub>O<sub>2</sub> 200 μM, 1hr ($1.853 \pm 0.2$, $n=3$), H<sub>2</sub>O<sub>2</sub> 200 μM, 3hr ($2.544 \pm 0.2$, $n=3$)<br>One-way ANOVA with Tukey's multiple comparisons test, $F(2,6) = 19.94$, $p = 0.002$<br>Ctrl vs. H<sub>2</sub>O<sub>2</sub> 200 μM, 1hr, $p = 0.03$; Ctrl vs. H<sub>2</sub>O<sub>2</sub> 200 μM, 3hr, $p = 0.002$; H<sub>2</sub>O<sub>2</sub> 200 μM, 1hr vs. H<sub>2</sub>O<sub>2</sub> 200 μM, 3hr, $p = 0.07$ |

## Notes

### Summary of Updates

The main text and figures have been revised. We have added the results of PD, ALS, and RA.

## References and Notes

1 Bulters, D. et al. Haemoglobin scavenging in intracranial bleeding: biology and clinical implications. Nat Rev Neurol 14, 416–432 (2018). 10.1038/s41582-018-0020-0

2 Biagioli, M. et al. Unexpected expression of alpha- and beta-globin in mesencephalic dopaminergic neurons and glial cells. Proc Natl Acad Sci U S A 106, 15454–15459 (2009). 10.1073/pnas.0813216106

3 Altinoz, M. A. et al. Involvement of hemoglobins in the pathophysiology of Alzheimer’s disease. Exp Gerontol 126, 110680 (2019). 10.1016/j.exger.2019.110680

4 Codrich, M. et al. Neuronal hemoglobin affects dopaminergic cells’ response to stress. Cell Death Dis 8, e2538 (2017). 10.1038/cddis.2016.458

5 Zheng, R., Yan, Y., Pu, J. & Zhang, B. Physiological and Pathological Functions of Neuronal Hemoglobin: A Key Underappreciated Protein in Parkinson’s Disease. Int J Mol Sci 23 (2022). 10.3390/ijms23169088

6 Sankar, S. B., Donegan, R. K., Shah, K. J., Reddi, A. R. & Wood, L. B. Heme and hemoglobin suppress amyloid beta-mediated inflammatory activation of mouse astrocytes. J Biol Chem 293, 11358–11373 (2018). 10.1074/jbc.RA117.001050

7 Amri, F., Ghouili, I., Tonon, M. C., Amri, M. & Masmoudi-Kouki, O. Hemoglobin-Improved Protection in Cultured Cerebral Cortical Astroglial Cells: Inhibition of Oxidative Stress and Caspase Activation. Front Endocrinol (Lausanne) 8, 67 (2017). 10.3389/fendo.2017.00067

8 Chun, H. et al. Severe reactive astrocytes precipitate pathological hallmarks of Alzheimer’s disease via H(2)O(2)(-) production. Nat Neurosci 23, 1555–1566 (2020). 10.1038/s41593-020-00735-y

9 Chun, H. & Lee, C. J. Reactive astrocytes in Alzheimer’s disease: A double-edged sword. Neurosci Res 126, 44–52 (2018). 10.1016/j.neures.2017.11.012

10 Nam, M. H., Sa, M., Ju, Y. H., Park, M. G. & Lee, C. J. Revisiting the Role of Astrocytic MAOB in Parkinson’s Disease. Int J Mol Sci 23 (2022). 10.3390/ijms23084453

11 Woo, T. G. et al. Novel chemical inhibitor against SOD1 misfolding and aggregation protects neuron-loss and ameliorates disease symptoms in ALS mouse model. Commun Biol 4, 1397 (2021). 10.1038/s42003-021-02862-z

12 Forman, H. J. & Zhang, H. Targeting oxidative stress in disease: promise and limitations of antioxidant therapy. Nat Rev Drug Discov 20, 689–709 (2021). 10.1038/s41573-021-00233-1

13 Won, W. et al. Inhibiting peripheral and central MAO-B ameliorates joint inflammation and cognitive impairment in rheumatoid arthritis. Exp Mol Med 54, 1188–1200 (2022). 10.1038/s12276-022-00830-z

14 Butterfield, D. A. & Halliwell, B. Oxidative stress, dysfunctional glucose metabolism and Alzheimer disease. Nat Rev Neurosci 20, 148–160 (2019). 10.1038/s41583-019-0132-6

15 Banaszak, M., Gorna, I., Wozniak, D., Przyslawski, J. & Drzymala-Czyz, S. The Impact of Curcumin, Resveratrol, and Cinnamon on Modulating Oxidative Stress and Antioxidant Activity in Type 2 Diabetes: Moving beyond an Anti-Hyperglycaemic Evaluation. Antioxidants (Basel) 13 (2024). 10.3390/antiox13050510

16 Shin, J. H. et al. Concurrent blockade of free radical and microsomal prostaglandin E synthase-1-mediated PGE2 production improves safety and efficacy in a mouse model of amyotrophic lateral sclerosis. J Neurochem 122, 952–961 (2012). 10.1111/j.1471-4159.2012.07771.x

17 Gajhede, M., Schuller, D. J., Henriksen, A., Smith, A. T. & Poulos, T. L. Crystal structure of horseradish peroxidase C at 2.15 A resolution. Nat Struct Biol 4, 1032–1038 (1997). 10.1038/nsb1297-1032

18 Vidossich, P. et al. On the role of water in peroxidase catalysis: a theoretical investigation of HRP compound I formation. J Phys Chem B 114, 5161–5169 (2010). 10.1021/jp911170b

19 Casado, A., Encarnacion Lopez-Fernandez, M., Concepcion Casado, M. & de La Torre, R. Lipid peroxidation and antioxidant enzyme activities in vascular and Alzheimer dementias. Neurochem Res 33, 450–458 (2008). 10.1007/s11064-007-9453-3

20 Ighodaro, O. & Akinloye, O. First line defence antioxidants-superoxide dismutase (SOD), catalase (CAT) and glutathione peroxidase (GPX): Their fundamental role in the entire antioxidant defence grid. Alexandria journal of medicine 54, 287–293 (2018).

21 Goyal, M. M. & Basak, A. Human catalase: looking for complete identity. Protein Cell 1, 888–897 (2010). 10.1007/s13238-010-0113-z

22 Wang, X. et al. Pyruvate protects mitochondria from oxidative stress in human neuroblastoma SK-N-SH cells. Brain Res 1132, 1–9 (2007). 10.1016/j.brainres.2006.11.032

23 Lee, J. D. et al. Structure-guided engineering of a fast genetically encoded sensor for real-time H_2_O_2_ monitoring. bioRxiv, 2024.2001.2031.578117 (2024). 10.1101/2024.01.31.578117

24 Yoon, B. E. et al. Glial GABA, synthesized by monoamine oxidase B, mediates tonic inhibition. J Physiol 592, 4951–4968 (2014). 10.1113/jphysiol.2014.278754

25 Haettig, J., Sun, Y., Wood, M. A. & Xu, X. Cell-type specific inactivation of hippocampal CA1 disrupts location-dependent object recognition in the mouse. Learn Mem 20, 139–146 (2013). 10.1101/lm.027847.112

26 Lee, S. et al. Channel-mediated tonic GABA release from glia. Science 330, 790–796 (2010). 10.1126/science.1184334

27 An, H. et al. Adenovirus-induced Reactive Astrogliosis Exacerbates the Pathology of Parkinson’s Disease. Exp Neurobiol 30, 222–231 (2021). 10.5607/en21013

28 Schonberger, S. J., Edgar, P. F., Kydd, R., Faull, R. L. & Cooper, G. J. Proteomic analysis of the brain in Alzheimer’s disease: molecular phenotype of a complex disease process. Proteomics 1, 1519–1528 (2001). 10.1002/1615-9861(200111)1:12<1519::aid-prot1519>3.0.co;2-l

29 Jo, S. et al. GABA from reactive astrocytes impairs memory in mouse models of Alzheimer’s disease. Nat Med 20, 886–896 (2014). 10.1038/nm.3639

30 Grigorieva, D. V. et al. Measurement of plasma hemoglobin peroxidase activity. Bull Exp Biol Med 155, 118–121 (2013). 10.1007/s10517-013-2095-3

31 Derat, E., Shaik, S., Rovira, C., Vidossich, P. & Alfonso-Prieto, M. The effect of a water molecule on the mechanism of formation of compound 0 in horseradish peroxidase. J Am Chem Soc 129, 6346–6347 (2007). 10.1021/ja0676861

32 Somin, S., Kulasiri, D. & Samarasinghe, S. Alleviating the unwanted effects of oxidative stress on Abeta clearance: a review of related concepts and strategies for the development of computational modelling. Transl Neurodegener 12, 11 (2023). 10.1186/s40035-023-00344-2

33 Koivisto, H. et al. Chronic Pyruvate Supplementation Increases Exploratory Activity and Brain Energy Reserves in Young and Middle-Aged Mice. Front Aging Neurosci 8, 41 (2016). 10.3389/fnagi.2016.00041

34 More, J. et al. N-Acetylcysteine Prevents the Spatial Memory Deficits and the Redox-Dependent RyR2 Decrease Displayed by an Alzheimer’s Disease Rat Model. Front Aging Neurosci 10, 399 (2018). 10.3389/fnagi.2018.00399

35 Vanni, S. et al. Hemoglobin mRNA Changes in the Frontal Cortex of Patients with Neurodegenerative Diseases. Front Neurosci 12, 8 (2018). 10.3389/fnins.2018.00008

36 Orre, M. et al. Acute isolation and transcriptome characterization of cortical astrocytes and microglia from young and aged mice. Neurobiol Aging 35, 1–14 (2014). 10.1016/j.neurobiolaging.2013.07.008

37 Spellman, D. S. et al. Development and evaluation of a multiplexed mass spectrometry based assay for measuring candidate peptide biomarkers in Alzheimer’s Disease Neuroimaging Initiative (ADNI) CSF. Proteomics Clin Appl 9, 715–731 (2015). 10.1002/prca.201400178

38 Zille, M. et al. Neuronal Death After Hemorrhagic Stroke In Vitro and In Vivo Shares Features of Ferroptosis and Necroptosis. Stroke 48, 1033–1043 (2017). 10.1161/STROKEAHA.116.015609

39 Straub, A. C. et al. Endothelial cell expression of haemoglobin alpha regulates nitric oxide signalling. Nature 491, 473–477 (2012). 10.1038/nature11626

40 Li, Y. P. et al. Erythropoietin attenuates Alzheimer-like memory impairments and pathological changes induced by amyloid beta42 in mice. Brain Res 1618, 159–167 (2015). 10.1016/j.brainres.2015.05.031

41 Freed, J. & Chakrabarti, L. Defining a role for hemoglobin in Parkinson’s disease. NPJ Parkinsons Dis 2, 16021 (2016). 10.1038/npjparkd.2016.21

42 Dassault Systèmes BIOVIA, Discovery Studio Modeling Environment, Release 2021, San Diego: Dassault Systèmes, 2023

43 Henriksen, A., Smith, A. T. & Gajhede, M. The Structures of the Horseradish Peroxidase C-Ferulic Acid Complex and the Ternary Complex with Cyanide Suggest How Peroxidases Oxidize Small Phenolic Substrates*. Journal of Biological Chemistry 274, 35005–35011 (1999). 10.1074/jbc.274.49.35005

44 Park, S.-Y., Yokoyama, T., Shibayama, N., Shiro, Y. & Tame, J. R. H. 1.25 Å Resolution Crystal Structures of Human Haemoglobin in the Oxy, Deoxy and Carbonmonoxy Forms. Journal of Molecular Biology 360, 690–701 (2006). 10.1016/j.jmb.2006.05.036

45 Berglund, G. I. et al. The catalytic pathway of horseradish peroxidase at high resolution. Nature 417, 463–468 (2002). 10.1038/417463a

46 Kamaljeet, Bansal, S. & SenGupta, U. A Study of the Interaction of Bovine Hemoglobin with Synthetic Dyes Using Spectroscopic Techniques and Molecular Docking. Frontiers in Chemistry 4 (2017). 10.3389/fchem.2016.00050

47 Pathak, K. V., Chiu, T.-L., Amin, E. A. & Turesky, R. J. Methemoglobin Formation and Characterization of Hemoglobin Adducts of Carcinogenic Aromatic Amines and Heterocyclic Aromatic Amines. Chemical Research in Toxicology 29, 255–269 (2016). 10.1021/acs.chemrestox.5b00418

48 Puscas, C. et al. The high affinity of small-molecule antioxidants for hemoglobin. Free Radical Biology and Medicine 124, 260–274 (2018). 10.1016/j.freeradbiomed.2018.06.019

49 Reeder, B. J. et al. Tyrosine Residues as Redox Cofactors in Human Hemoglobin: IMPLICATIONS FOR ENGINEERING NONTOXIC BLOOD SUBSTITUTES*. Journal of Biological Chemistry 283, 30780–30787 (2008). 10.1074/jbc.M804709200

50 Sharma, M., Farhat, N., Khan, A. U., Khan, F. H. & Mahmood, R. Studies on the interaction of 2,4-dibromophenol with human hemoglobin using multi-spectroscopic, molecular docking and molecular dynamics techniques. Journal of Biomolecular Structure and Dynamics, 1–11 (2023). 10.1080/07391102.2023.2264975

51 Faller, B. Artificial membrane assays to assess permeability. Curr Drug Metab 9, 886–892 (2008). 10.2174/138920008786485227

52 Lee, C. J. et al. Astrocytic control of synaptic NMDA receptors. J Physiol 581, 1057–1081 (2007). 10.1113/jphysiol.2007.130377

53 Stine, W. B., Jr., Dahlgren, K. N., Krafft, G. A. & LaDu, M. J. In vitro characterization of conditions for amyloid-beta peptide oligomerization and fibrillogenesis. J Biol Chem 278, 11612–11622 (2003). 10.1074/jbc.M210207200

54 Ju, Y. H. et al. Astrocytic urea cycle detoxifies Abeta-derived ammonia while impairing memory in Alzheimer’s disease. Cell Metab 34, 1104–1120 e1108 (2022). 10.1016/j.cmet.2022.05.011

55 Park, J. H. et al. Newly developed reversible MAO-B inhibitor circumvents the shortcomings of irreversible inhibitors in Alzheimer’s disease. Sci Adv 5, eaav0316 (2019). 10.1126/sciadv.aav0316

56 Chun, H. et al. Severe reactive astrocytes precipitate pathological hallmarks of Alzheimer’s disease via H2O2(-) production. Nat Neurosci 23, 1555–1566 (2020). 10.1038/s41593-020-00735-y

57 Ventura, A. et al. Cre-lox-regulated conditional RNA interference from transgenes. Proc Natl Acad Sci U S A 101, 10380–10385 (2004). 10.1073/pnas.0403954101

